# Synthetic Mucins as Glycan-Defined Prebiotics

**DOI:** 10.1101/2025.01.27.635133

**Authors:** Jill W. Alty, Carolyn E. Barnes, Agnese Nicoli, Bradley S. Turner, Ekua A. Beneman, Amanda E. Dugan, Spencer D. Brucks, Austin G. Kruger, Richard R. Schrock, Katharina Ribbeck, Laura L. Kiessling

## Abstract

The human microbiome contains at least as many bacterial cells as human cells. The mucosal layer that lines all epithelial cells organizes, cultivates, and regulates these bacterial inhabitants. Some commensal bacteria offer benefits, like improving gut barrier function, suppressing pathobiont growth, and modulating host immunity. These health benefits have fueled the popularity of probiotics, but their retention is often hindered by their low colonization efficiency and mucosal adhesion. Mucins, the primary structural components of the mucosal layer, are essential for the organization and regulation of microbial populations, promoting growth and offering sites for adhesion through their multivalent presentation of *O*-glycosylation. The molecular mechanisms of mucin– probiotic interactions remain understudied due, in part, to the inability to incisively manipulate native mucin sequences or the glycans they bear. In this investigation, we developed synthetic mucins with defined glycan presentations to interrogate glycan-dependent interactions between mucus and probiotic *Lactobacillus* species. Though synthetic mucins can dampen the effects of pathogens, toxins, and viruses, their impact on probiotic bacteria as prebiotics or binding sites is unclear. We synthesized mucin surrogates that bind to three investigated *Lactobacillus* species. The nutrient conditions under which bacteria were cultured influenced glycan binding preferences, suggesting mucin–probiotic interactions change with nutrient availability. The addition of synthetic mucins to native mucin increased *Lactobacillus fermentum* adherence. Additionally, an increase in *Lactobacillus* glycosidase activity indicated that native and synthetic mucins both function as prebiotics, as probiotic bacteria can cleave the displayed *O*-glycans. Thus, synthetic mucins can cultivate target probiotic bacteria and increase adhesion as binding sites, highlighting their value as tools for elucidating native mucin functions and as promising agents for promoting human health.

## Introduction

The human microbiome is primarily housed within the mucosal layer lining all epithelial cells and consists of trillions of bacteria, encompassing disease-causing pathogens or beneficial commensals. Some commensal bacteria are probiotic, conferring health benefits to the host. Regarded as valuable probiotics, bacteria from the genus *Lactobacillus* are often found in exogenous sources like fermented food products. *Lactobacillus* spp. mitigate irritable bowel syndrome, alter the pathogenicity of other microbes, and regulate host immunity by reducing inflammation and activating immune cells.^1-3^ *Lactobacillus* spp. can produce short-chain fatty acids through the fermentation of carbohydrates, improving gut microbiota dysbiosis, promoting phage production, and, in some cases, eliciting anti-tumor responses.^4-7^ Supplementation of probiotic *Lactobacillus* to augment these populations is a promising strategy for maintaining a healthy gut microbiome and treating diseases such as ulcerative colitis. However, the retention of probiotic bacteria is challenging as they are typically cleared from the body in three to ten days due to their low mucosal adhesion and colonization efficiency.^8, 9^

Mucins, the main structural components of the mucosal layer, are a family of proteins characterized by their extended backbone conformation and dense *O*-glycosylation. The structural characteristics of mucins and their *O*-glycosylation patterns are essential for organizing, cultivating, and regulating microbial populations by adhering microbes and serving as nutrient sources (Figure 1A).^10^ These functions support the growth of probiotic, mucus-binding commensals such as *Lactobacillus* spp.^11-13^

**Figure 1.**
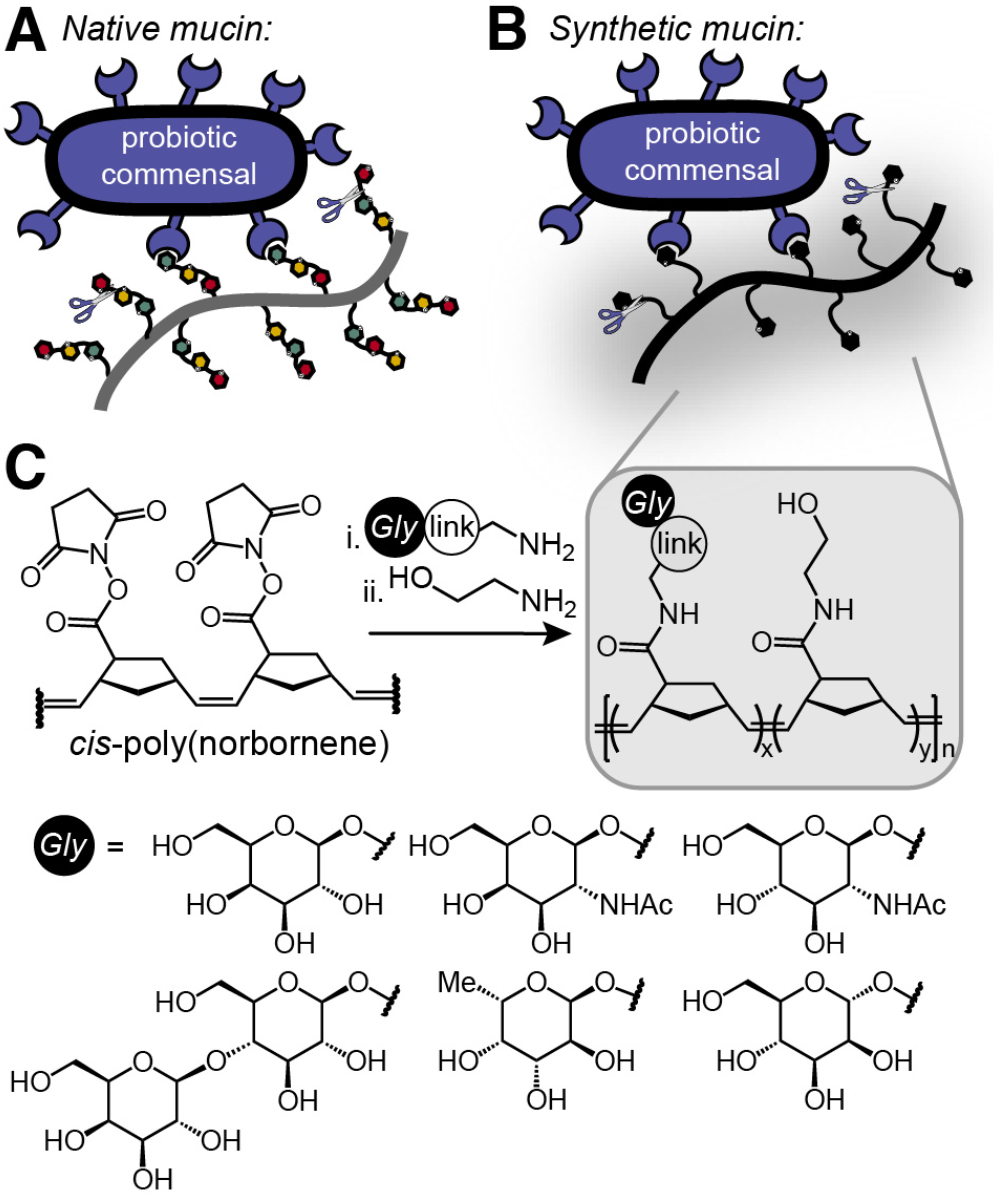
A) Native mucins as prebiotics for adhesion and nutrition of probiotics. B) Synthetic mucins as prebiotics. C) Mucin surrogate synthesis. Link is ethanolamine, triethylene glycol, or aryl.

As key nutrient sources and adhesion sites for gut bacteria, mucins may be regarded as prebiotics, non-digestible materials that stimulate the growth and activity of probiotics.^14-16^ Prebiotics serve as nutrients for probiotics, encouraging bacterial production of short-chain fatty acids and modulating immune function through their interactions with toll-like receptors and commensal bacteria.^17, 18^ Common prebiotics include galacto- and fructo-oligosaccharides, though mucin glycans have also been shown to be effective prebiotics.^19^ The use of mucins as prebiotics has emerged as a strategy to manipulate the gut microbiome. This approach may more effectively diversify microbial community composition than live probiotic supplementation.^16^

A critical feature of mucins is their multivalent display of glycans. While it is understood that bacteria bind native mucins via their *O*-glycans, the inability to manipulate the chemical structure of native mucins complicates efforts to identify the features responsible. Despite advances, challenges remain, including difficulties in recombinantly expressing native mucins, the inability to genetically control mucin glycosylation, and the complexity of synthesizing mucin glycans.^20-23^ Synthetic polymers displaying defined glycans can function as mucin mimetics, overcoming the challenges of unknown and heterogeneous glycosylation patterns.^24-27^ To date, studies with synthetic mucins have focused on inhibiting pathogenic bacteria or virulence pathways. We envisioned that synthetic mucins could act as prebiotics to enhance probiotic adhesion and inform the binding preferences of commensal bacteria (Figure 1B).

Herein, we synthesize a suite of chemically defined synthetic mucins to probe their function as prebiotics for probiotic *Lactobacillus* species. These mimics display native mucin saccharides on an extended synthetic backbone (Figure 1C). We identified agents that bound probiotic commensals, including *L. plantarum, L. fermentum*, and *L. reuteri*. In each case, we reveal the glycan specificity for retaining each organism. Bacterial binding to synthetic mucins was highly dependent on the nutrient sources available during growth, with nutrient-rich media leading to glycan-specific binding. In contrast, nutrient-poor media resulted in more promiscuous binding, highlighting how the environment influences the ability of probiotics to adhere to these mucin-like compounds. The multivalent display of the mimetics induced clustering and adhesion of the bacterial species. When *L. fermentum* was incubated in the presence of both native and synthetic mucins, more commensal bacteria adhered to the mucosa, indicating synthetic mucins were prebiotic. We also detected glycosidase production, which leads to *O*-glycan cleavage, presumably further cultivating probiotic growth. This report presents a platform to investigate how native mucins bind probiotics and function as prebiotics through tailored synthetic mucin surrogates.

## Results

### Design of synthetic mucins

To identify the monosaccharides involved in the interactions between mucins and *Lactobacillus* spp., a collection of amine-terminated, *O*-glycan epitopes was synthesized (Figure 1C). These epitopes represented the most prevalent groups within native mucin glycan sequences: α-fucose (α-Fuc), β-galactose (β-Gal), β-*N*-acetyl galactosamine (β-GalNAc), β-*N*-acetyl glucosamine (β-GlcNAc), as well as the disaccharide lactose (β-Lac).^28, 29^ Though present in trace quantities on native mucins in *N*-linked glycans,^30^ we included α-mannose (α-Man) residues in this survey as the epitope has been identified to bind *L. plantarum* and *L. fermentum*.^31-36^ We typically linked the saccharide residues to the polymer scaffold through a triethylene glycol (PEG_3_) group to minimize steric effects and thereby facilitate bacterial binding.^37^

The polymeric backbone used for the synthetic mucins was synthesized using a *cis*-selective ring-opening metathesis polymerization of a norbornene derivative to provide an extended scaffold that mimics the morphology of native mucins, increases the steric accessibility of the glycans, and improves water solubility.^24^ The glycan was installed by exposing a glycan building block with an amine-terminated anomeric linker to the polymer bearing pendent *N*-hydroxy succinimidyl esters (Figure 1C).^38^ The degree of glycan substitution was varied (20–80%) and as anticipated, lower glycan densities tend to result in optimal binding of the bacteria (Figure S1).^24, 39, 40^ For visualizing the mimetics, approximately one fluorophore per chain was added using Alexa Fluor 405 (AF405) cadaverine prior to quenching any remaining succinimidyl ester with ethanolamine. We focused on 100-mer synthetic mucins decorated with ∼20-50% glycan (Figure S2).

### Glycan-specific binding of synthetic mucins to *L. plantarum*

To test whether bacteria interact with the polymers, we first examined *L. plantarum*. Because *L. plantarum* binds native mucus and α-mannose through mucus-binding proteins and mannose-specific adhesins, we used this probiotic to test the utility of a flow cytometry-based assay to assess mucin-mimetic binding to bacteria.^31, 32, 41-44^ Native or synthetic mucins were incubated with live bacteria, and then the cells were washed to remove any unbound glycopolymer. The percentage of cells bound to the AF405-containing polymer was then determined by cellular fluorescence.

Mannosylated synthetic mucins served as positive controls and exhibited dose-dependent binding to *L. plantarum* (Figure 2A, Figure S3). Polymers substituted with an aryl-mannoside (α-ArMan) were compared to those with an α-mannoside (α-PEG_3_Man) appended through an anomeric substituent to probe the linker’s impact. Indeed, different affinities were observed for α-ArMan polymers versus α-PEG_3_Man polymers binding to *L. plantarum*. To ensure that ligands occupied similar binding sites to mucin *O*-glycans, we generally employed glycan substituents with oligoethylene glycol linkers. Therefore, we validated that glycan binding preferences could be assessed using flow cytometry with *L. plantarum* and mannose-functionalized mucin surrogates.

**Figure 2.**
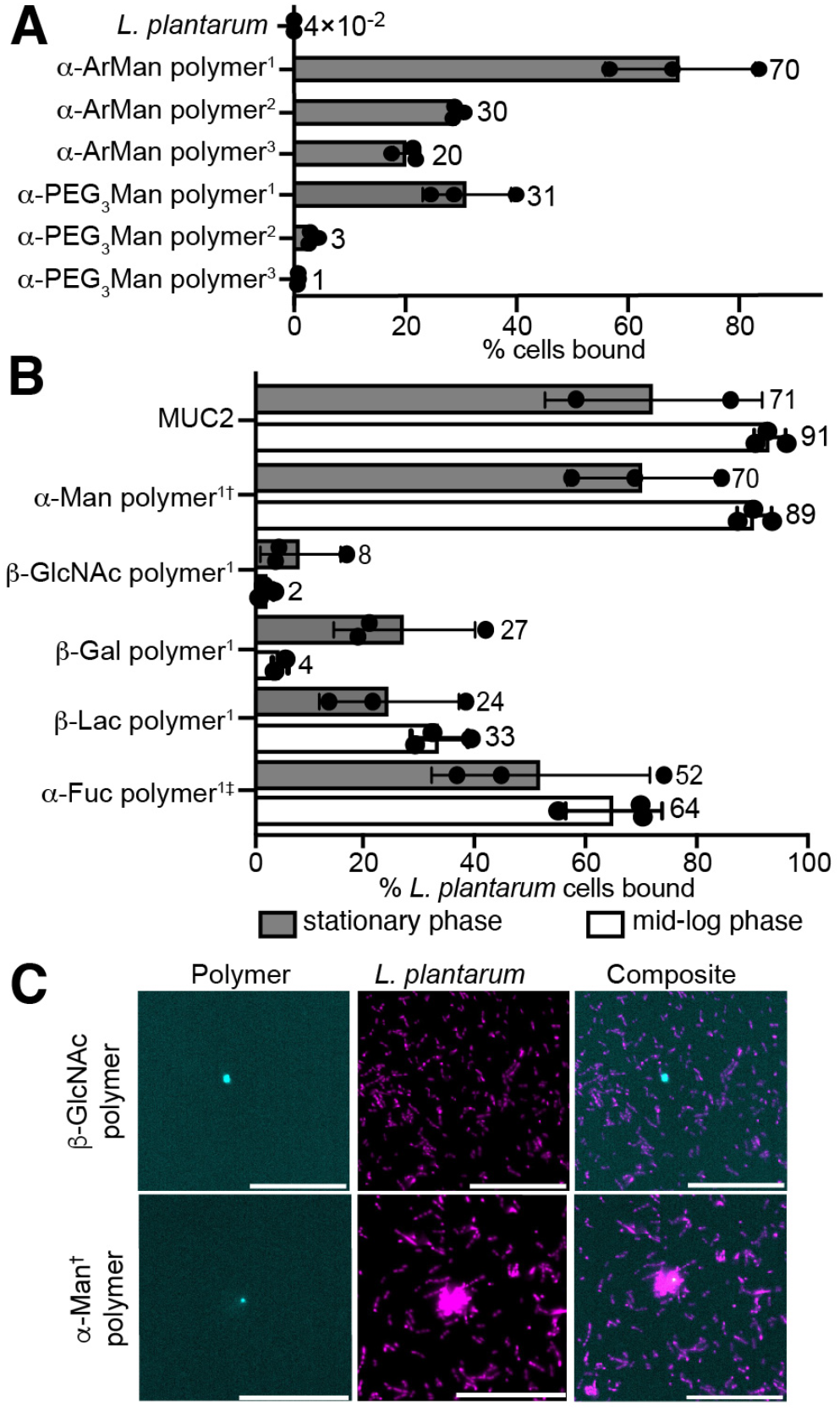
A & B) Percentage of cells bound calculated from flow cytometry of native and synthetic mucins with *L. plantarum* grown to stationary or mid-log phase. Concentrations: ^1^2 mM, ^2^0.5 mM, and ^3^0.05 mM glycan for polymers; 0.04-0.1 wt% for MUC2. Saccharide linker is PEG_3_ unless † for aryl or ‡ for ethanolamine. Functionalizations: β-Gal = 40%, β-Lac = 20%, β-GlcNAc = 40-50%, α-Fuc = 30%, α-Man = 20%, α-Man = 15-25%. C) Confocal microscopy of synthetic mucin and bacteria. Magenta: bacteria in stationary phase; Cyan: polymer. Scale bar, 40 mm. Data are representative of biological replicates (n β 3).

*L. plantarum* binding to the collection of synthetic mucins was compared to its interaction with native gut mucin, MUC2 (Figure 2B, Figure S5–S7). Growth phase and available nutrients may affect probiotic bacteria’s fitness and their ability to colonize the human gut.^45^ Hence, we compared the probiotic binding when grown to stationary phase, or when growth has reached saturation, versus in mid-logarithmic (mid-log) phase, when growth is exponential. In both phases, *L. plantarum* cells showed excellent binding to MUC2 with increased binding in mid-log phase. For synthetic mucins bearing a single glycan epitope, α-Man polymer and α-Fuc polymer exhibited the strongest binding, agnostic of growth phase. Visualizing by confocal microscopy, the α-Man and α-Fuc synthetic mucins displayed the most coincidence of polymer and *L. plantarum* fluorescence, whereas no colocalization was observed for those synthetic mucins that flow cytometry indicated could not bind (Figure 2C).

### Growth conditions impact the binding of synthetic mucins to *L. reuteri*

We hypothesized that the mucin surrogates could be used to determine whether glycan-binding preferences change under different growth conditions. When grown in nutrient-rich conditions (MRS broth) to stationary or mid-log phase, *L. reuteri* bound specifically to synthetic mucins displaying β-Lac or α-Fuc as judged by flow cytometry (Figure 3A, Figure S8-S9) and confocal microscopy (Figure 3C). Physiological settings can be more nutrient-poor. We hypothesized bacteria grown in such conditions would bind a greater variety of synthetic mucins to enhance their ability to colonize a nutrient-scarce gut.^45^ Indeed, when bacteria were cultured in minimal media (Table S1), mucin binding by *L. reuteri* was enriched. Moreover, the previously observed glycan preferences were not apparent, as *L. reuteri* binding was enhanced for all mucin surrogates (Figure 3C, Figure S10).

**Figure 3.**
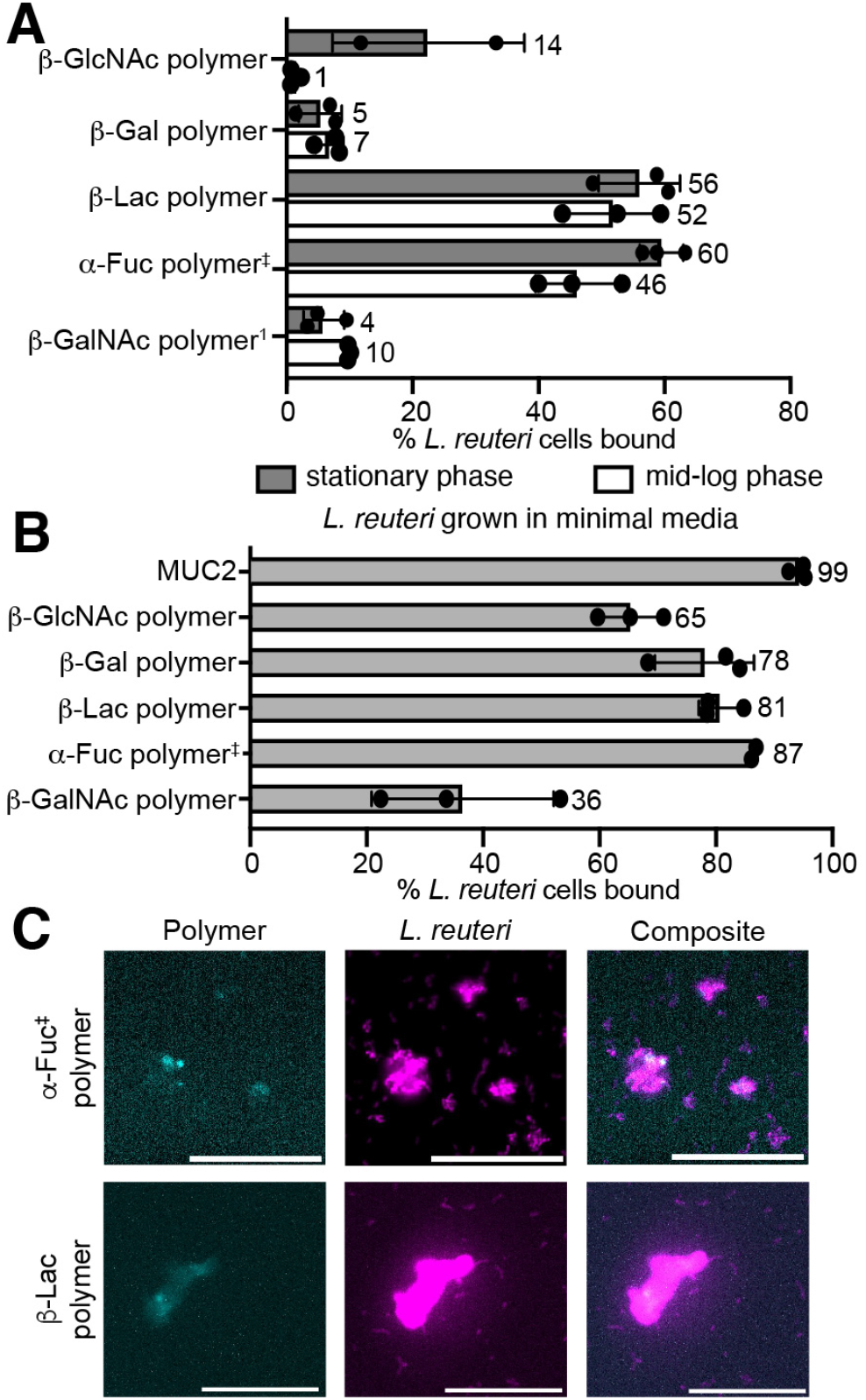
Percentage of cells bound calculated from flow cytometry of native and synthetic mucins with *L. reuteri* in MRS broth (A) and in minimal media (B). Concentrations: 2 mM glycan for polymers, 0.02-0.06 wt% for MUC2. Saccharide linker is PEG_3_ unless ‡ for ethanolamine. Functionalizations: β-Gal = 40-45%, β-Lac = 10-20%, β-GlcNAc = 40-50%, α-Fuc = 25-50%, β-GalNAc = 30%. ^1^Concentration in stationary phase: 0.5 mM. C) Confocal microscopy of synthetic mucin and bacteria. Magenta: bacteria in stationary phase; Cyan: polymer. Scale bar, 40 mm. Data are representative of biological replicates (n ≥ 3).

### Manipulation of mucin interactions of *L. fermentum*

Probiotic *L. fermentum* is closely related to *L. reuteri* and is known to express mucin-binding proteins.^11, 46-48^ Like *L. reuteri, L. fermentum* bound most avidly to synthetic mucins bearing β-lactose residues. When this species was grown to mid-log phase, it was less selective, binding most synthetic mucins (Figure 4A, Figure S12). Synthetic mucins presenting β-lactose bound 97% of *L. fermentum*, outperforming native mucin.

**Figure 4.**
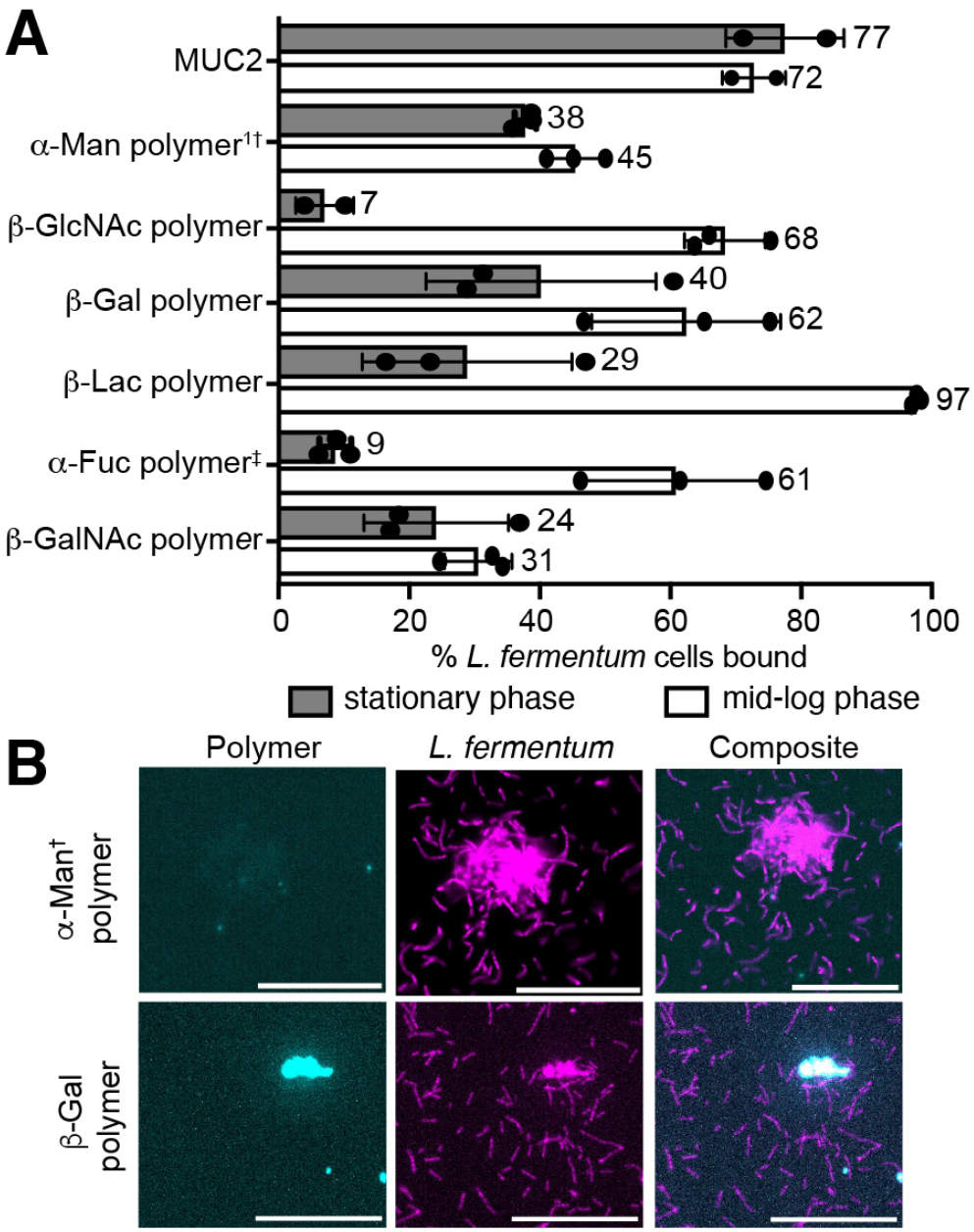
A) Percentage of cells bound calculated from flow cytometry of native or synthetic mucins with *L. fermentum* cultured in stationary and mid-log phases. Concentrations: 2 mM glycan for polymers, 0.02-0.04 wt% for MUC2. Saccharide linker is PEG_3_ unless † for aryl or ‡ for ethanolamine. Functionalizations: β*-*Gal = 40-45%, β*-*Lac = 15-20%, β*-*GlcNAc = 40-50%, α*-*Fuc= 25-30%, β*-*GalNAc = 30%, α-Man = 20-40%. ^1^Concentration in stationary phase: 0.5 mM. B) Confocal microscopy shows colocalization of synthetic mucin and bacteria. Magenta: bacteria in stationary phase; Cyan: polymer. Scale bar, 40 mm. Data are representative of biological replicates (n β 3).

Confocal microscopy revealed that synthetic mucins bearing α-Man and β-Gal colocalize with *L. fermentum*, further supporting the glycan specificity observed with flow cytometry (Figure 4B). As bacterial growth slowed in the stationary phase, the commensal’s affinity for synthetic mucins diminished, while native mucins retained excellent binding (Figure 4A, Figure S13).

### Clustering of *Lactobacillus* spp. through multivalency

Because native and synthetic mucins both present multivalent arrays of glycans, they can engage multiple bacteria to promote cell clustering or agglutination. To quantify the extent of bacterial clustering by native and synthetic mucins, we analyzed the size of cell clusters using microscopy.^49^ For each species, the synthetic mucin that bound the most bacterial cells also produced the largest average cluster size (Figure S15-S17). When *L. plantarum* was exposed to polymers bearing α-Fuc or α-Man, the population of single cells decreased by 50% or 60%, respectively, and the cell cluster size ranged up to 5 microns (Figure S15). In contrast, the non-binding synthetic mucins had no effect on microbial clustering. MUC2 modestly agglutinated *L. fermentum*, while synthetic mucins presenting β-Gal or α-Man decreased the population of single cells, giving rise to a broad dispersity of bacterial clusters (Figure 5A, Figure S17). We envisioned leveraging this outcome to enhance bacterial attachment to mucosal membranes.

**Figure 5.**
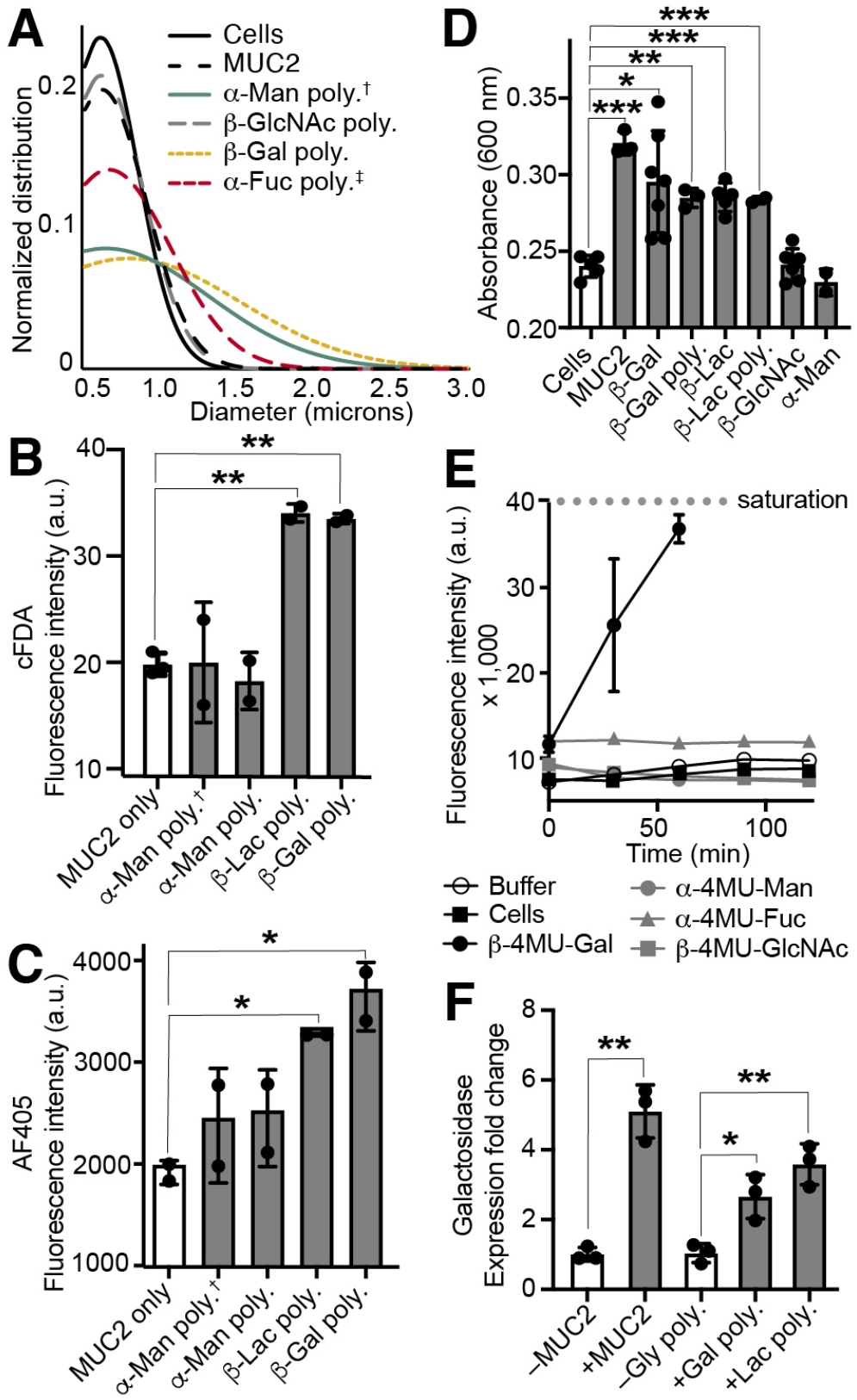
A) *L. fermentum* cellular clustering by native and synthetic mucins visualized by confocal microscopy and quantified using Fiji software. Saccharide linker is PEG_3_ unless † for aryl or ‡ for ethanolamine. B) *L. fermentum* adhesion to MUC2-coated wells in the presence or absence of synthetic mucins measured by cFDA fluorescence. C) Synthetic mucin retention to MUC2-coated wells with *L. fermentum* measured by AF405 fluorescence. D) Nutritional glycan preferences of *L. fermentum* in minimal media with mono- or di-saccharides as monitored by adhesion assay. E) Glycosidase activity of *L. fermentum* as monitored by fluorogenic glycans. F) qPCR analysis of β-galactosidase expression for *L. fermentum* grown in the presence or absence of native or synthetic mucins. Statistical analysis was performed using ANOVA with multiple comparisons test: *P<0.1, **P<0.01, ***P<0.001, ****P<0.0001. Data are representative of biological replicates (n β 2).

### Synthetic mucins as prebiotics for *L. fermentum* through enhanced adhesion

As *Lactobacillus* species are typically cleared by the body in three to ten days, improving the retention of probiotics in the gut is crucial to maximize their beneficial effects.^17, 18^ Thus, emerging prebiotics that enhance the adhesion of probiotic *Lactobacillus* spp. to native mucins are under investigation.^50^ To test whether the synthetic mucins could function as prebiotics, *L. fermentum* was incubated on a MUC2-modified plate surface to generate a pseudo-mucosa.^51^ AF405 fluorescence confirmed that the synthetic mucin remained post-washing (Figure 5C, Figure S19). Supplementation with synthetic mucin led to enhanced adhesion. The retentive activity of the synthetic mucins was glycan-specific, with enhanced adhesion when polymers with either β-Lac or β-Gal substituents were added (Figure 5B, Figure S19). In contrast, the non-binding polymer with β-GlcNAc residues failed to increase *L. fermentum* adhesion.

### Nutrition of *L. fermentum* via monovalent and multivalent glycans

For best results as a prebiotic, the synthetic mucin would increase probiotic adhesion and serve as a nutrient source to cultivate probiotic microbes within the mucosa. Therefore, we assessed microbial growth on the synthetic mucins. *L. fermentum* increased proliferation in the presence of free β-lactose or β-galactose, the same epitopes that increased bacterial binding and adhesion to MUC2 when appended to synthetic mucins (Figure 5D, Figure S20). This trend translated to multivalent scaffolds: MUC2 and the synthetic mucins displaying β-Gal or β-Lac significantly enhanced bacterial growth.

### Glycosidase expression and manipulation

Upon binding to endogenous mucins, bacteria upregulate the production of glycosidases to harvest glycans. We assessed the presence of genes encoding glycosidases of the three *Lactobacillus* species. The genomes of *L. reuteri, L. fermentum*, and *L. plantarum* were analyzed using the Kyoto Encyclopedia of Genes and Genomes database^52^ to determine the presence of genes encoding glycosidases of interest. All three species encode at least one predicted β-galactosidase. To determine whether these glycosidases were expressed and possess the expected enzymatic activity, we used monosaccharides conjugated to the reporter 4-methylumbelliferone (4MU). These substrates were incubated with each bacterial species, and the release of fluorescent 4MU was monitored. When bacteria were grown in nutrient-poor media supplemented with Gal-4MU, they retained efficient galactosidase activity (Figure 5E and Figure S21). No glycosidase activity for α-Man, α-Fuc, or β-GlcNAc was detected with *L. reuteri* or *L. fermentum*. We did observe efficient β-GlcNAc-ase activity with *L. plantarum*, though no binding to the GlcNAc-substituted synthetic mucin was detected (Figure S21).

Because nutrient-rich growth conditions gave rise to nutrient-specific fitness, we investigated if there were changes in galactosidase expression on a transcriptional level based on growth conditions. We grew *L. fermentum* in nutrient-rich media supplemented with native or synthetic mucin. Quantitative real-time PCR (qPCR) was used to quantify the relative change in β-galactosidase expression. We found the expression of β-galactosidase genes by *L. fermentum* was increased in either MUC2- or synthetic mucin-doped media (Figure 5F, Figure S22-23). Native and synthetic mucins bearing the preferred glycan, β-Gal or β-Lac, enhanced the bacteria’s ability to procure nutrients by increasing its galactosidase expression.

## Discussion

Bacteria in the mucosal layer often engage with mucins via glycan-binding enzymes and proteins.^53^ We first examined *L. plantarum* as a model probiotic due to its known mannose- and mucin-binding proteins.^31, 32, 41-44^ In assessing the activity of chemically defined synthetic mucins, we found that *L. plantarum* preferentially binds to α-Man and α-Fuc polymers, regardless of growth phase. These observations are consistent with the similar arrangements of the hydroxyl groups within these two glycan epitopes. Across all studies, α-ArMan polymers outperformed α-PEG_3_Man polymers. Initially designed for binding to dendritic cell lectin DC-SIGN, the arylated α-Man ligand engages in π–π interactions with aromatic amino acid residues in the binding pocket, which may be occurring in the described mucin- or mannose-binding proteins expressed by bacteria.^54^

Aromatic substituents can enhance the glycan affinity for many lectins.^55-57^ However, to ensure that our synthetic mucins occupied similar binding sites to native mucins, we generally used oligoethylene glycol-linked *O*-glycans.

While it is known that growth medium and strain diversity can influence *L. reuteri* affinity for native mucins,^58, 59^ we reasoned that the mucin mimics could reveal differences in glycan specificity. Regardless of the growth phase, *L. reuteri* grown in nutrient-rich conditions preferentially bound polymers bearing α-Fuc or β-Lac, common native mucin termini. In contrast, when the bacteria were grown in nutrient-poor minimal media conditions, probiotic bacteria bound synthetic mucins indiscriminately. Taken together, these data indicate mucin binding of *L. reuteri* depends on bacterial growth conditions and favors promiscuous attachment in nutrient-poor physiological settings, likely contributing to the overall fitness of this probiotic.

Synthetic mucin binding to *L. fermentum* depended less on the glycan epitopes than did *L. reuteri* or *L. plantarum*, the latter of which bound preferentially to two distinct glycan epitopes. The isolated surface mucin-binding domain MBD_93_ of *L. fermentum* interacted with mucin glycans GalNAc, GlcNAc, and Gal.^48^ These findings align well with the binding profile we observed when *L. fermentum* was grown to mid-log phase.

However, the glycan epitope with the highest binding observed for *L. fermentum* was the β-Lac polymer in mid-log phase (97%), outperforming native mucin binding. As future work further characterizes binding profiles of probiotic bacteria, it will be interesting to understand how more complex glycans, such as disaccharides or core glycan sequences, impact binding.

Native mucins can function as prebiotics by increasing the adhesion of probiotic bacteria to the mucosa.^14-16^ Because the multivalent display of *O*-glycans induced clustering of the probiotic *Lactobacillus* bacteria, we hypothesized that the bacteria would better adhere to mucosa upon addition of synthetic mucin. Upon generation of a pseudo-mucosa through a mucin-coated plate, we observed that synthetic mucins increased the adhesion of *L. fermentum*. Interestingly, the increased retention of the probiotic to the pseudo-mucosa was glycan-dependent, corroborating our evidence that β-Lac and β-Gal polymers are effective binders of *L. fermentum*. These findings indicate that synthetic mucins can act as prebiotics and promote adhesion through glycan-specific binding to facilitate the retention of *Lactobacillus* spp.

In addition to improving bacterial adhesion, prebiotics are typically metabolized by probiotic commensal bacteria to encourage growth. To evaluate if the polymers were nutrient sources for *Lactobacillus* probiotics, we compared bacterial growth rates when grown in media supplemented with different glycan additives in the form of a free saccharide or the multivalent synthetic mucin substrates. The growth of *L. fermentum* depended on the identity of the glycan displayed by the mucin mimic, with more cellular growth observed when grown in the presence of β-Gal or β-Lac-bearing compounds. These polymers also effectively interacted with *L. fermentum*, suggesting that mucin mimics displaying these glycans could improve adhesion and function as a nutrient source, thus serving as novel prebiotics.

To access nutrients in the gut, bacteria produce glycosidases to degrade complex glycans to yield sustainable nutrients. Mucin-processing enzymes have been well-characterized for commensals, such as *Akkermansia muciniphila*.^60-63^ In contrast, few reports detail the expression of different glycosidases and their dependence on mucins for *Lactobacillus* species.^64, 65^ Using fluorogenic glycan substrates, glycosidase activity was monitored for *L. plantarum, L. reuteri*, and *L. fermentum*. The observed galactosidase activity suggests that *L. reuteri* and *L. fermentum* could use β-Gal or β-Lac synthetic mucins as prebiotics to bind mucin-binding proteins as an anchoring point and cleave these *O*-glycans for nutrients through glycosidases. In addition to galactosidase activity, *N*-acetylglucosaminidase activity was observed for *L. plantarum*. This observation could be relevant to prior reports that show *L. plantarum* encodes a protein with a chitin-binding domain, providing a mechanism to bind poly(β-GlcNAc).^66^ The glycosidase expression could be manipulated through growth conditions, with increased expression observed on a transcriptional level when native or synthetic mucins were added to the bacterial growth conditions. These data provide further support that the synthetic mucins can cultivate and manipulate *Lactobacillus* probiotics.

## Conclusion

Synthetic mucins are powerful, chemically tractable tools to characterize mucin–probiotic interactions. The glycan specificity was determined for three *Lactobacillus* species using chemically defined mucin surrogates. Mucin mimics bearing relevant binding agents facilitated enhanced adherence, retention, and growth of *Lactobacillus* spp. to mixtures of natural and synthetic mucins, suggesting synthetic mucins can serve as prebiotics. These findings underscore the value of elucidating mucin–bacteria interactions. In this way, mucin surrogates can facilitate the development of supplements that support a healthy microbiome.

## Abbreviations

4-MU: 4-methylumberriferyl
AF405: AlexaFluor 405
α-ArMan: aryl α-mannosylated
cFDA: carboxyfluorescein diacetate
DMSO: dimethylsulfoxide
α-Fuc: fucosylated
β-Gal: β-galactosylated
β-GalNAc: *N*-acetyl galactosaminated
β-GlcNAc: *N*-acetyl glucosaminated
HEPES: 4-(2-hydroxyethyl)-1-piperazineethanesulfonic acid buffer
β-Lac: lactosylated
MRS: Man,Rogosa, and Sharpe broth
OD600: optical density at 600 nm
PEG_3_: triethylene glycol
qPCR: quantitative polymerase chain reaction

## Supporting Information

Experimental procedures, materials, instrumentation, and additional tables and figures including minimal media conditions, bacterial strains, synthetic routes to polymers, flow cytometry data, microscopy and clustering data, adhesion results, OD600 data, glycosidase activity assays, qPCR data, and ^1^H NMR spectra.

## Author Information

### Notes

L.L.K. reports compensation for consulting and/or SAB membership from Exo Therapeutics, the ONO Pharmaceutical Foundation, and Coca Cola unrelated to this research. The remaining authors declare that they have no competing interests. Protocols involving samples from human participants were approved by the Massachusetts Institute of Technology’s Committee on the Use of Humans as Experimental Subjects.

## Acknowledgments

The authors thank Dr. George Degen for providing native mucin and Dr. René Riedel for synthesizing catalyst. This work was carried out in part through MIT.nano facilities. We also thank the MIT Department of Chemistry Instrumentation Facility.

