## Supplemental Information for "Synthetic Mucins as Glycan-Defined Prebiotics"

Supporting Information


1. General Procedures………………………………………………………………. 2
    a. Synthetic Materials…………………………………………………………. 2
    b. Strains and Growth Conditions……………………………………………... 2
    c. Synthetic Characterization………………………………………………….. 2
    d. Broth and Buffer Recipes…………………………………………………… 3
    e. Safety comment…………………………………………………………….. 3
2. Figures Referenced in the Main Text…………………………………………….. 4
    a. **Figure S1:** Polymer functionalization optimization……..……………………… 4
    b. **Figure S2:** Polymer molecular weight optimization……..……………………... 5
    c. **Figure S3:** Flow cytometry Gating strategy………..……..…………………….. 6
    d. **Figure S4:** *L. plantarum* and *L. fermentum* flow with mannosylated polymers… 7
    e. **Figure S5:** *L. plantarum* stationary flow cytometry…………………………….. 8
    f. **Figure S6:** *L. plantarum* stationary microscopy..……………………………… 11
    g. **Figure S7:** *L. plantarum* mid-log flow cytometry………………………………13
    h. **Figure S8:** *L. reuteri* stationary flow cytometry…………………….……..... 16
    i. **Figure S9:** *L. reuteri* mid-log flow cytometry……………………................... 18
    j. **Table S1:** Development of minimal media conditions for *L. reuteri*……………20
    k. **Figure S10:** *L. reuteri* minimal media flow cytometry……………………....... 21
    l. **Figure S11:** *L. reuteri* stationary microscopy..………………………………… 23
    m. **Figure S12:** *L. fermentum* mid-log flow cytometry……………………………. 24
    n. **Figure S13:** *L. fermentum* stationary flow cytometry………………………….. 27
    o. **Figure S14:** *L. fermentum* stationary microscopy..…………………………….. 29
    p. **Figure S15:** *L. plantarum* clustering..…………………………………...... 30
    q. **Figure S16**: *L. reuteri* clustering..……………..……………………………… 30
    r. **Figure S17:** *L. fermentum* clustering..…………………………………………. 31
    s. **Figure S18:** Adhesion optimization…………………………………………… 31
    t. **Figure S19:** *L. fermentum* adhesion..…………………………………………... 32
    u. **Figure S20:** OD600 for all bacteria …………………………………………… 33
    v. **Figure S21:** 4-methylumberriferyl glycans for glycosidase activity…………... 34
    w. **Table S2:** Real time qPCR primer sequences…………………………………35
    x. **Figure S22:** *L. fermentum* galactosidase expression with MUC2……………… 35
    y. **Figure S23:** *L. fermentum* galactosidase expression with cis-PNBs…………… 36
3. Ligand and Polymer Syntheses…………………………………………………..36
4. Flow Cytometry Protocol……………………………………………………….. 45
5. Microscopy Protocol ……………………………………………………………. 46
6. Clustering Protocol……………………………………………………………... 46
7. Adhesion Protocol ……………………………………………………………… 46
8. qPCR Protocol…………………………………………………………………..47
9. Ligand and Polymer NMRs and GPCs…………………………………………… 48
    a. **Figure S24:** *cis*-PNB 100-mer $^1$H NMR……………………………………...48
    b. **Figure S25:** *cis*-PNB 100-mer GPC……………………………………………..49
    c. **Figure S26**: *cis*-PNB 250-mer GPC……………………………………………..49

d. **Figure S27**: Gal *cis*-PNB $^1$H NMR……………………………………………...50
e. **Figure S28**: Lac *cis*-PNB $^1$H NMR……………………………………………...51
f. **Figure S29**: GlcNAc *cis*-PNB $^1$H NMR………………………………………52
g. **Figure S30**: GalNAc *cis*-PNB $^1$H NMR………………………………………53
h. **Figure S31**: Fuc *cis*-PNB $^1$H NMR……………………………………………...54
i. **Figure S32**: Man *cis*-PNB $^1$H NMR……………………………………………..55
j. **Figure S33**: ArMan *cis*-PNB $^1$H NMR…………………………………………..56
10. References………………………………………………………………………… 57

### 1. General Procedures

#### a) Synthetic Materials

Dichloromethane (DCM), methanol (MeOH), acetonitrile (ACN), tetrahydrofuran (THF), dimethylsulfoxide (DMSO), boron trifluoride diethyl etherate, β,D-galactose pentaacetate, exo-norbornene-5-carboxylic acid, sodium carbonate, *N*- hydroxysuccinimide (NHS), 1-Ethyl-3-(3-dimethylaminopropyl)carbodiimide (EDC), sodium metal, (1,3-Bis(2,4,6-trimethylphenyl)-2-imidazolidinylidene) dichloro(phenylmethylene)(tricyclohexylphosphine)ruthenium (Grubbs Catalyst 2nd generation), ethanolamine, Amberlyst A26 hydroxide form resin, 2-[4-(2-hydroxyethyl)piperazin-1-yl]ethanesulfonic acid (HEPES), sodium chloride, magnesium chloride hexahydrate, calcium chloride dihydrate, manganese chloride, Whatman PTFE 0.22 μm cutoff syringe filters, and 10,000 molecular weight cutoff (MWCO) cellulose dialysis tubing were purchased from Millipore Sigma. Diethyl ether (ether), hexanes, chloroform, acetone, ethyl acetate (EtOAc), 9-fluorenylmethoxycarbonyl chloride (Fmoc-Cl), piperidine, and *N*-methylmorpholine were purchased from Fisher Scientific. Dry solvents were obtained using a solvent purification system by Pure Process Technology. Flash chromatography (FC) was performed using SiliaFlash Irregular Silica Gel P60 from Silicycle and analytical thin layer chromatography (TLC) was performed using Siliaplate TLG-R10014B-323 with F254 fluorescence indicator. Porcine Muc2[1] was isolated and purified as previously described.

#### b) Strains and Growth Conditions

For bacterial assays, the strains employed include: *Limosilactobacillus reuteri* (Kandler et al.) F275 [DSM20016]; *Lactiplantibacillus plantarum* (Orla-Jensen) Zheng et al. NCIMB 8826 [Hayward 3A, WCFS1]; and *Lactobacillus fermentum* Beijerinck 36 [BUCSAV 233, NCDO 215, NCIB 6991, NCIB 8028, NCTC 6991]. *L. reuteri* was grown anaerobically in a falcon tube filled with *Lactobacilli* deMan, Rogosa, and Sharpe (MRS) broth (BD Difco™), minimizing air space in the tube, and incubated overnight at 37 °C without shaking. *L. plantarum* and *L. fermentum* were both grown aerobically in a 50 mL baffled Erlenmeyer flask with 15 mL MRS broth (BD, Difco™) at 37 °C with shaking at 190 rpm. Cell counts were determined by OD600 measurement on a BioMate 3S Spectrophotometer.

#### c) Synthetic Characterization

All nuclear magnetic resonance (NMR) spectra were obtained using a Bruker 400 MHz Avance Neo spectrometer, a Bruker 500 MHz Avance Neo spectrometer equipped with a BBFO

SmartProbe, or a Bruker 600 MHz Avance Neo spectrometer equipped with a 5mm helium-cooled QCI-F cryoprobe. Gel permeation chromatography (GPC) measurements were taken with a Tosoh EcoSEC HLC-8320 equipped with dual SuperH3000 columns running a chloroform mobile phase at 1 mL/min. All GPC instruments were calibrated to polystyrene standards. Absorbance and fluorescence measurements were made using a SpectraMax M5 plate reader.

**d) Broth and Buffer Recipes**

*HEPES with added calcium:*
750 mL MilliQ $H_2O$, 23.83 g HEPES (~0.1 M), 5.26 g sodium chloride, 95.2 mg magnesium chloride, 111 mg calcium chloride dihydrate, and 126 mg manganese chloride were combined. pH was adjusted from 5.36 to 7.4 using 10 M sodium hydroxide. The buffer was sterile- filtered prior to use.

*HEPES with EDTA:*
1 L MilliQ $H_2O$, 37.22 g EDTA (~100 mM), and 23.83 g HEPES (~0.1 M) were combined. pH was adjusted to 7.4 using 10 M sodium hydroxide. The buffer was sterile-filtered prior to use.

*MOD-MRS versus MRS:*
Media was autoclaved before use and only opened under a flame. MRS media was made according to instructions from BD Difco™ Lactobacillus broth, but the dry mixture should be composed of the contents described below. Modified MRS (MOD-MRS) was prepared using a recipe previously reported.[2]

| Component | MRS | MOD-MRS |
|---|---|---|
| Peptone | 10 g | 5 g |
| Beef/Meat Extract | 10 g | 4 g |
| Yeast Extract | 5 g | 2 g |
| Dextrose | 20 g | 0 g |
| Sorbitan Monooleate | 1 g | 0 g |
| Ammonium Citrate | 2 g | 0 g |
| Sodium Acetate | 5 g | 0.6 g |
| $MnSO_4$ x $H_2O$ | 0.05 g | 0.04 g |
| $Na_2HPO_4$ | 2.0 g | 3 g |
| Tween 80 | 0 | 0.5 mL |
| $K_2HPO_4$ | 0 | 1 g |
| $MgSO_4$ | 0 | 0.3 g |

**e) Safety comment:** No unexpected or unusually high safety hazards were encountered.

### 2. Figures Referenced in the Main Text

**Figure S1.** Flow cytometry optimization with *L. reuteri* after growth in minimal media with various degrees of functionalization. After overnight growth in MOD-MRS anaerobically, the ODs were measured as 0.122, 0.146, and 0.146. The assay was performed in biological triplicate

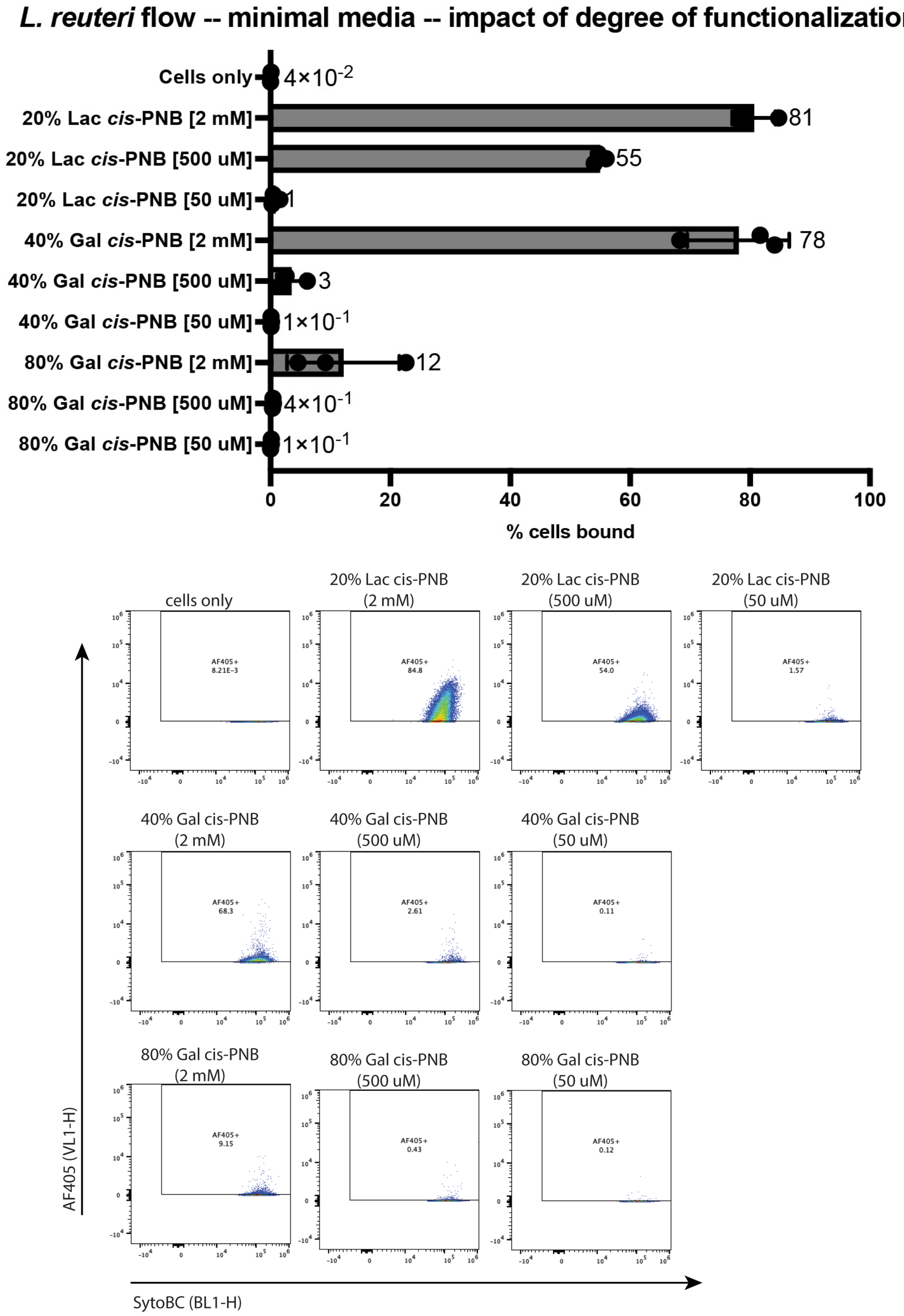

**Figure S2.** Flow cytometry optimization with *L. reuteri* after growth in minimal media with various molecular weights. After overnight growth in MOD-MRS anaerobically, the ODs were measured as 0.122, 0.146, and 0.146. The assay was performed in biological triplicate with technical triplicates in each plate.

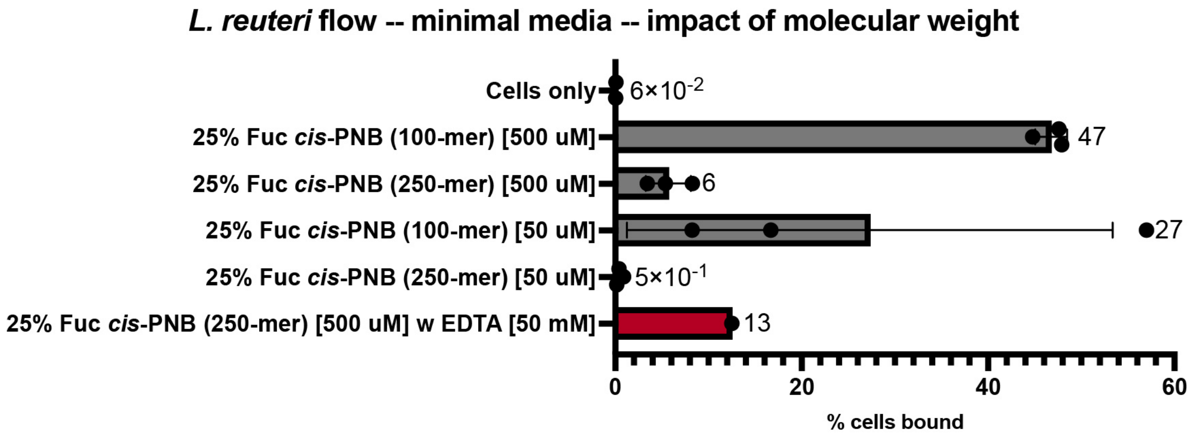


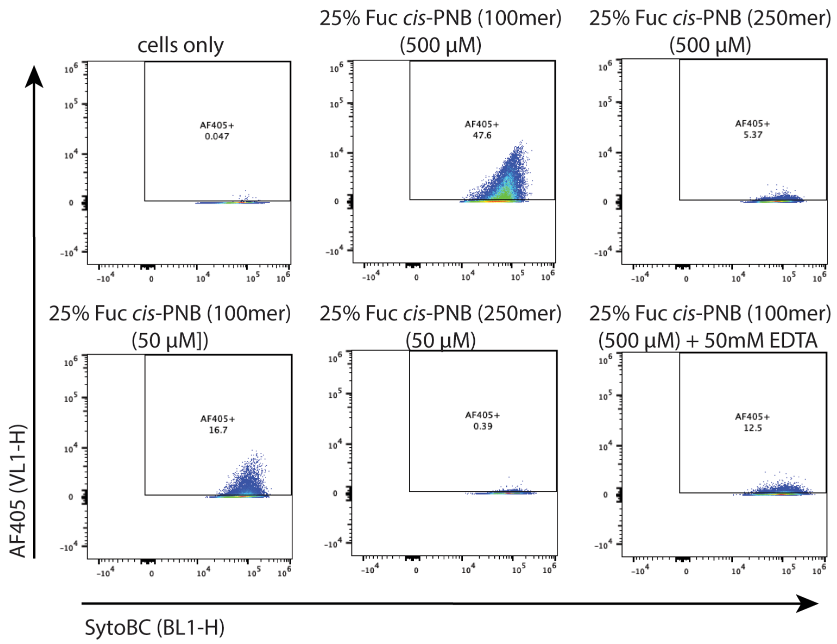

**Figure S3.** Gating strategy: The bacterial cells were first analyzed by FSC-H v. SSC-H to define the microbial population. Of the "Cells+" population, the cells were analyzed in SYTO-BC-H v. SSC-H to remove any debris that coincided with the population as only cells should be labeled by the nucleic acid stain. The "SYTO+" cells were then analyzed for polymer binding through gating SYTO-BC-H v. AF405-H. Any "AF405+" instances are considered "cells bound".

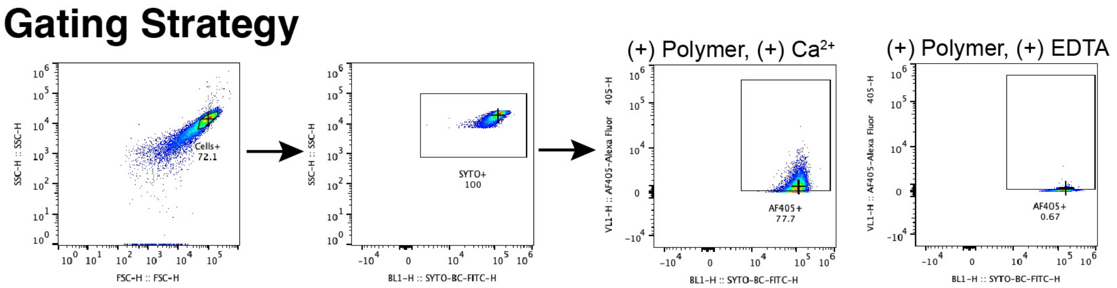

**Figure S4.** Flow cytometry gating sequence and dot plots from Mannosylated polymers binding *L. plantarum* and *L. fermentum* in stationary phase.

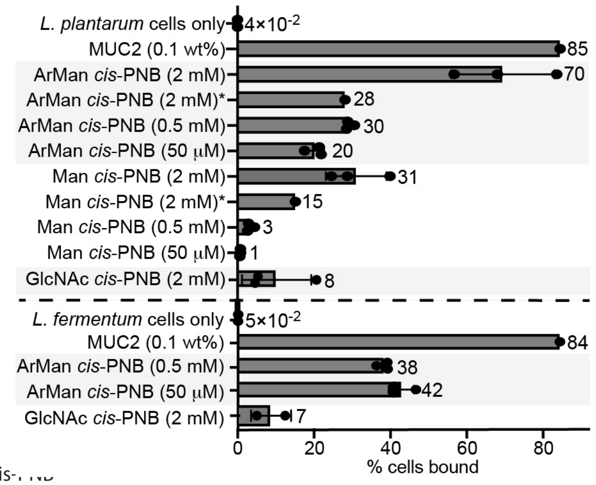


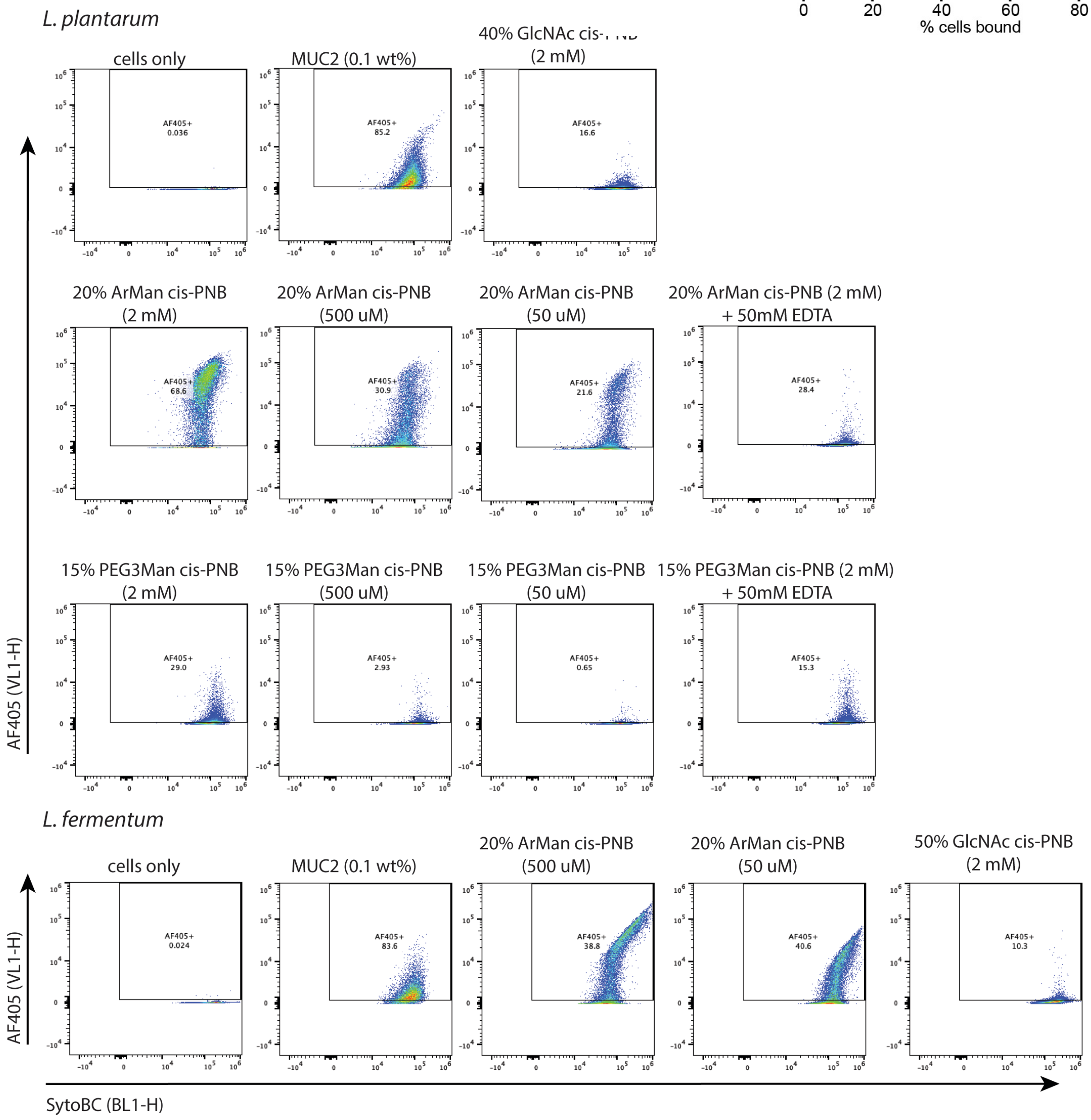

**Figure S5.** Flow cytometry of *L. plantarum* in stationary phase with various glycopolymer concentrations and glycan identities. The OD of *L. plantarum* after overnight growth was 1.85 and 1.79. The assay was performed in biological duplicate with technical triplicates in each plate.

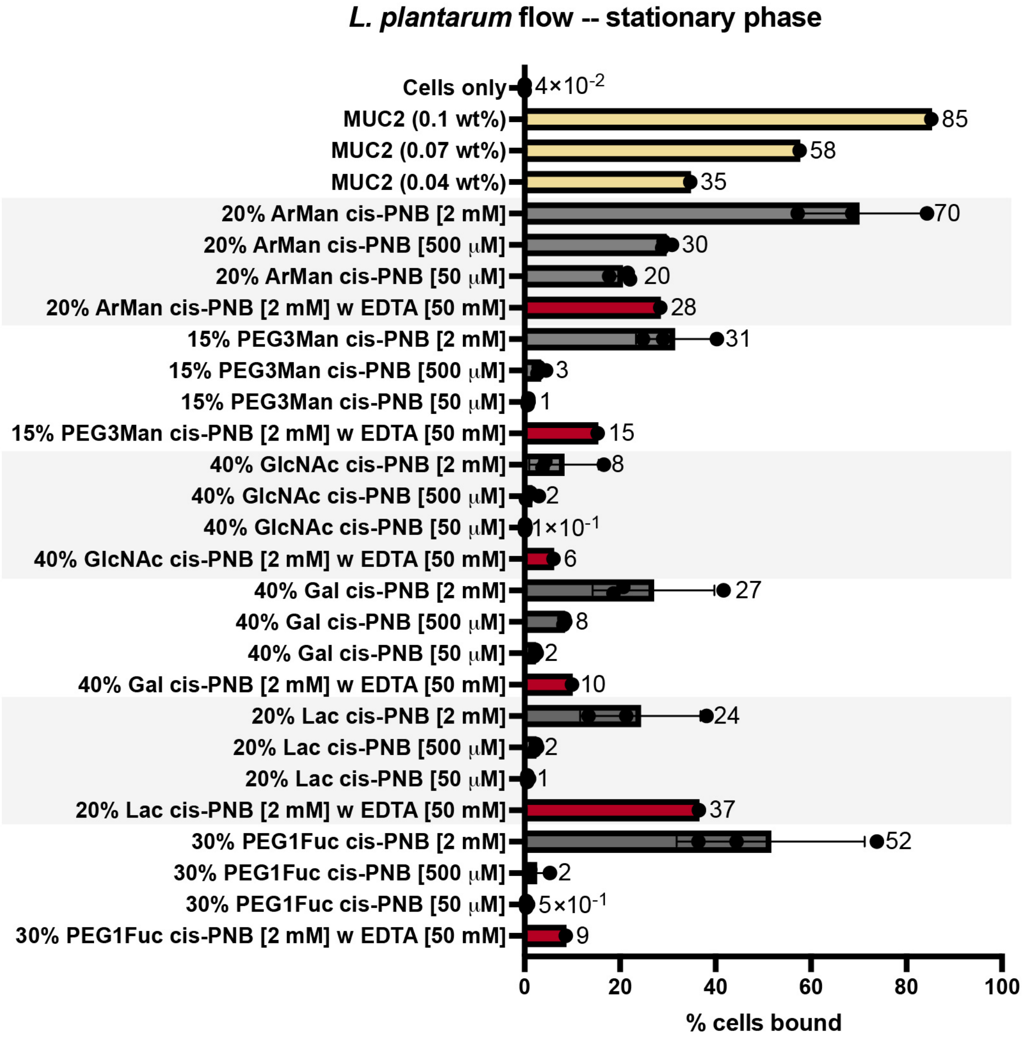

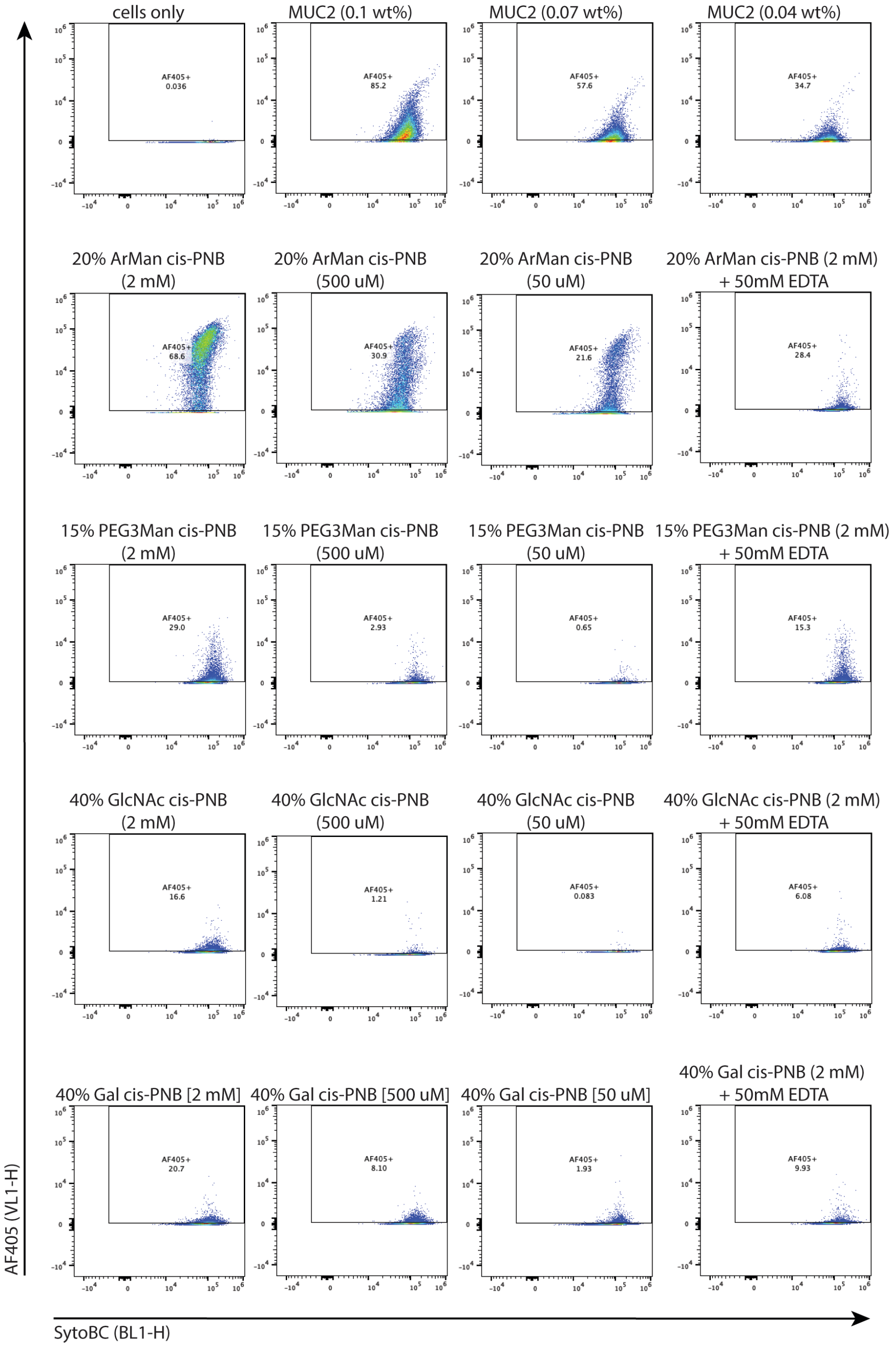
cells only
MUC2 (0.1 wt%)
MUC2 (0.07 wt%)
MUC2 (0.04 wt%)
AF405+ 0.036
AF405+ 85.2
AF405+ 57.6
AF405+ 34.7
20% ArMan cis-PNB (2 mM)
20% ArMan cis-PNB (500 uM)
20% ArMan cis-PNB (50 uM)
20% ArMan cis-PNB (2 mM) + 50mM EDTA
AF405+ 68.6
AF405+ 30.9
AF405+ 21.6
AF405+ 28.4
15% PEG3Man cis-PNB (2 mM)
15% PEG3Man cis-PNB (500 uM)
15% PEG3Man cis-PNB (50 uM)
15% PEG3Man cis-PNB (2 mM) + 50mM EDTA
AF405+ 29.0
AF405+ 2.93
AF405+ 0.65
AF405+ 15.3
40% GlcNAc cis-PNB (2 mM)
40% GlcNAc cis-PNB (500 uM)
40% GlcNAc cis-PNB (50 uM)
40% GlcNAc cis-PNB (2 mM) + 50mM EDTA
AF405+ 16.6
AF405+ 1.21
AF405+ 0.083
AF405+ 6.08
40% Gal cis-PNB [2 mM]
40% Gal cis-PNB [500 uM]
40% Gal cis-PNB [50 uM]
40% Gal cis-PNB (2 mM) + 50mM EDTA
AF405+ 20.7
AF405+ 8.10
AF405+ 1.93
AF405+ 9.93
AF405 (VL1-H)
SytoBC (BL1-H)

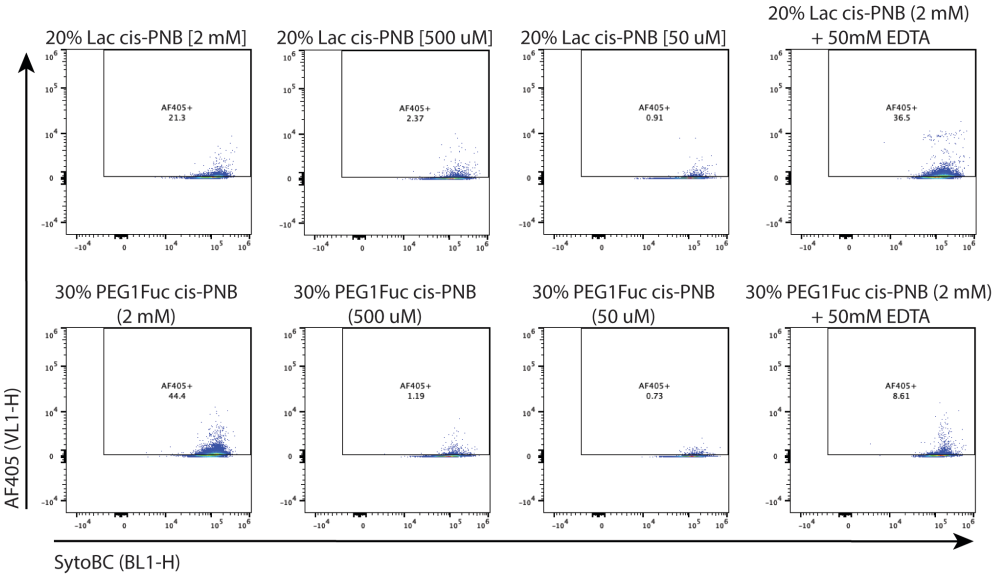
20% Lac cis-PNB [2 mM]
20% Lac cis-PNB [500 uM]
20% Lac cis-PNB [50 uM]
20% Lac cis-PNB (2 mM) + 50mM EDTA
AF405+ 21.3
AF405+ 2.37
AF405+ 0.91
AF405+ 36.5
30% PEG1Fuc cis-PNB (2 mM)
30% PEG1Fuc cis-PNB (500 uM)
30% PEG1Fuc cis-PNB (50 uM)
30% PEG1Fuc cis-PNB (2 mM) + 50mM EDTA
AF405+ 44.4
AF405+ 1.19
AF405+ 0.73
AF405+ 8.61
AF405 (VL1-H)
SytoBC (BL1-H)

**Figure S6.** Microscopy with *L. plantarum* in stationary phase. The images were collected in technical duplicate and biological duplicate. Prior to imaging, the cells and glycopolymer mixture post-flow cytometry were diluted 5:95.

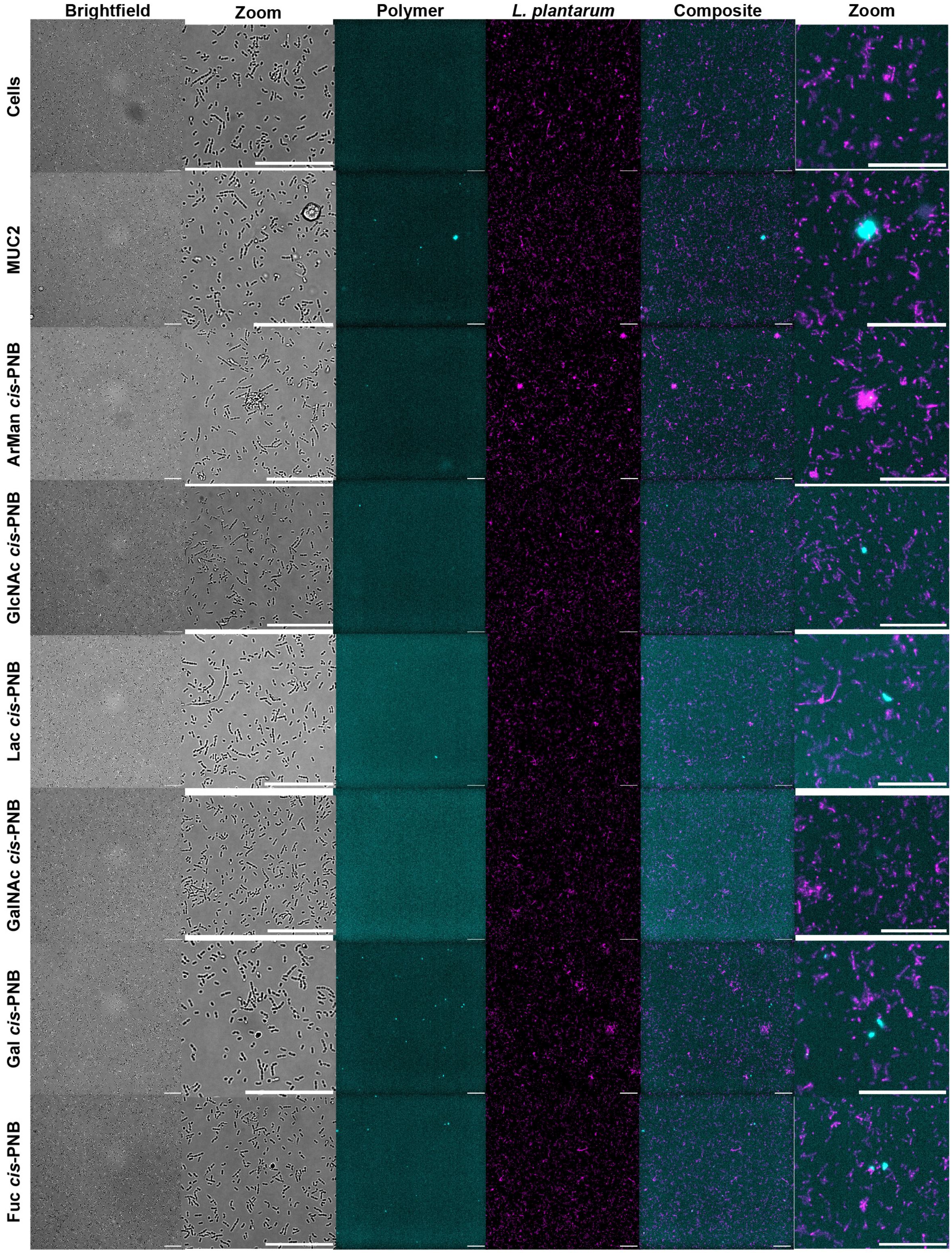

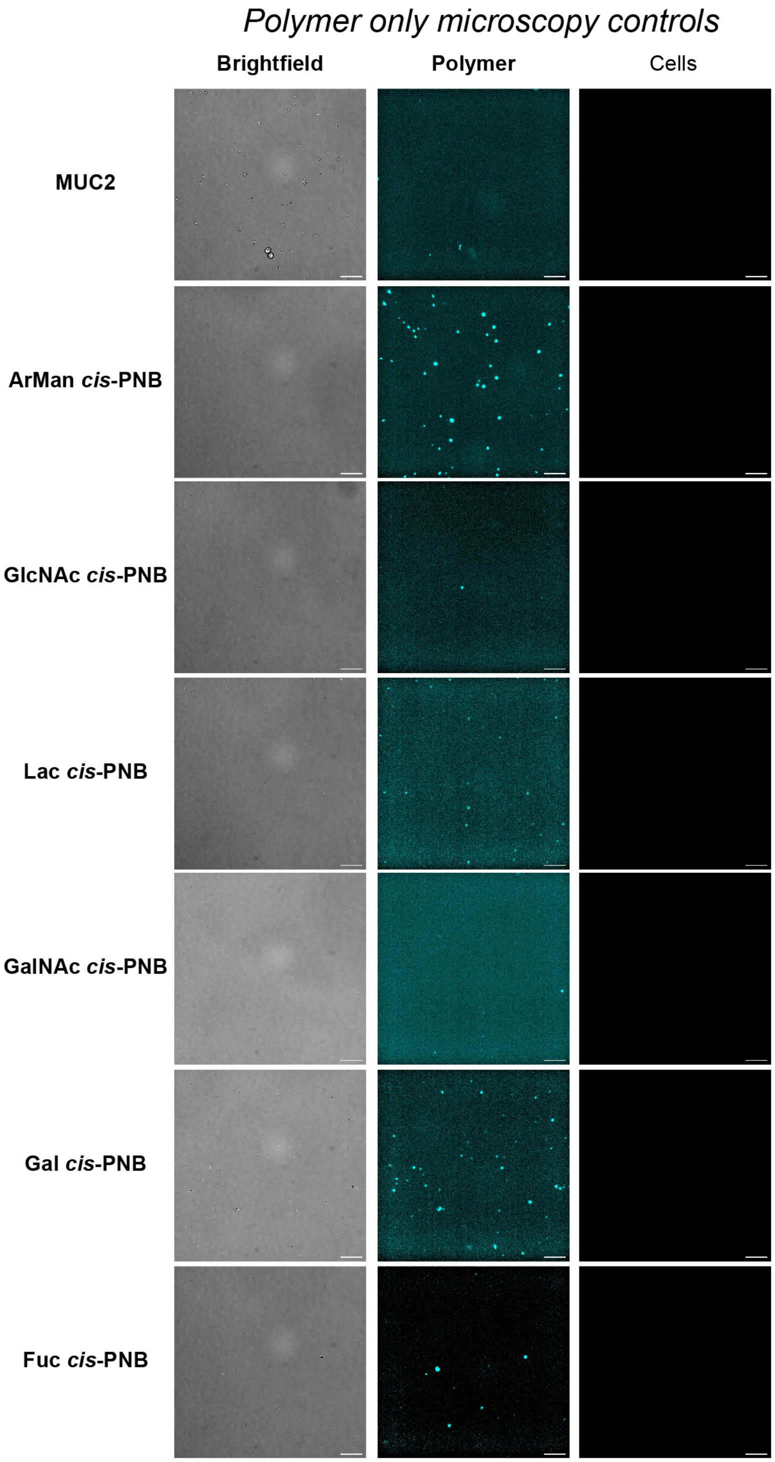

*Polymer only microscopy controls*
Brightfield
Polymer
Cells
MUC2
ArMan *cis*-PNB
GlcNAc *cis*-PNB
Lac *cis*-PNB
GalNAc *cis*-PNB
Gal *cis*-PNB
Fuc *cis*-PNB

**Figure S7.** Flow cytometry with *L. plantarum* in mid-log phase with various glycopolymer concentrations and glycan identities. The OD of *L. plantarum* after overnight was 1.79 & 1.80. The bacteria were allowed to recover for 1 hour after inoculating 1 mL of culture into 15 mL of media to OD = 0.62 & 0.61. The assay was performed in biological duplicate with technical triplicates in each plate.

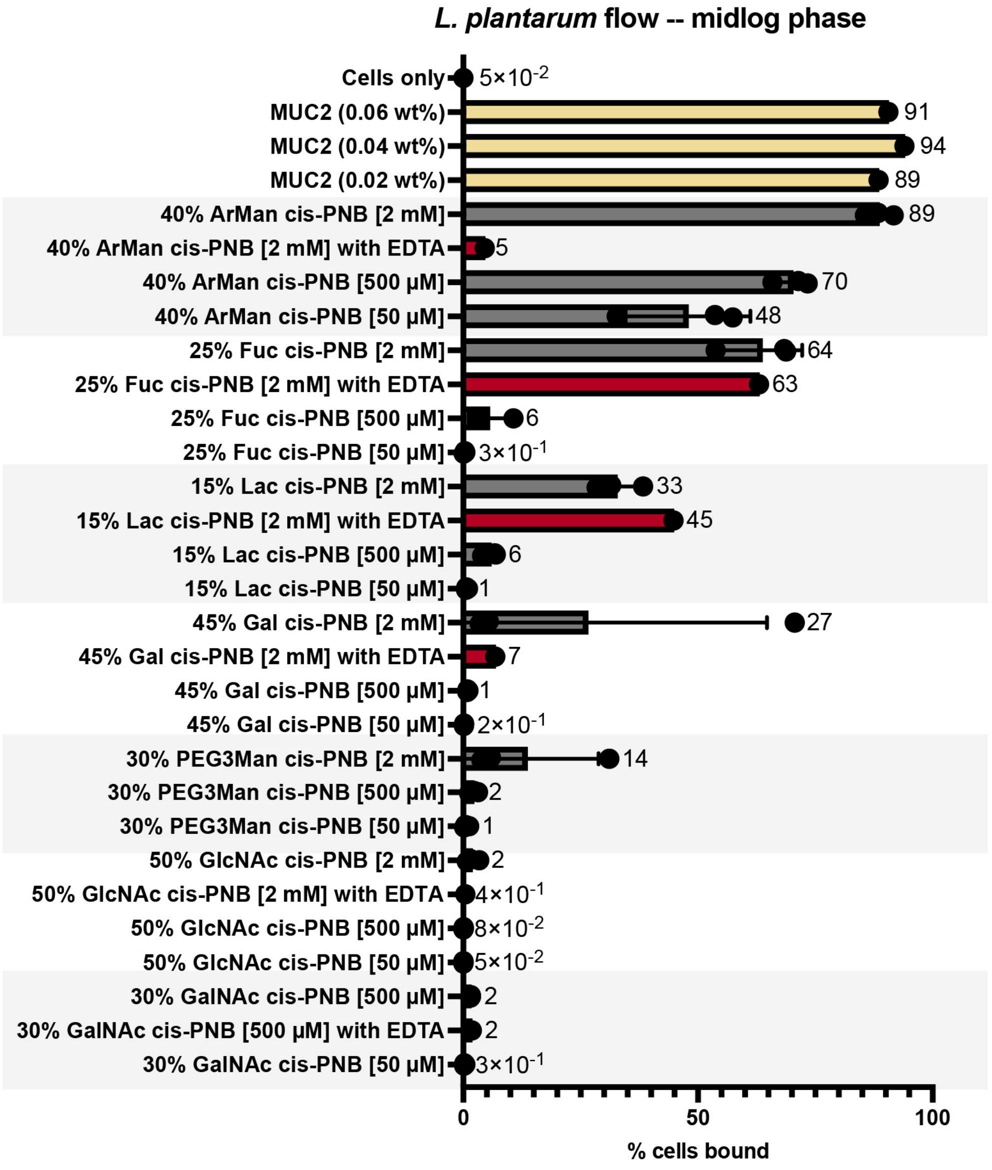

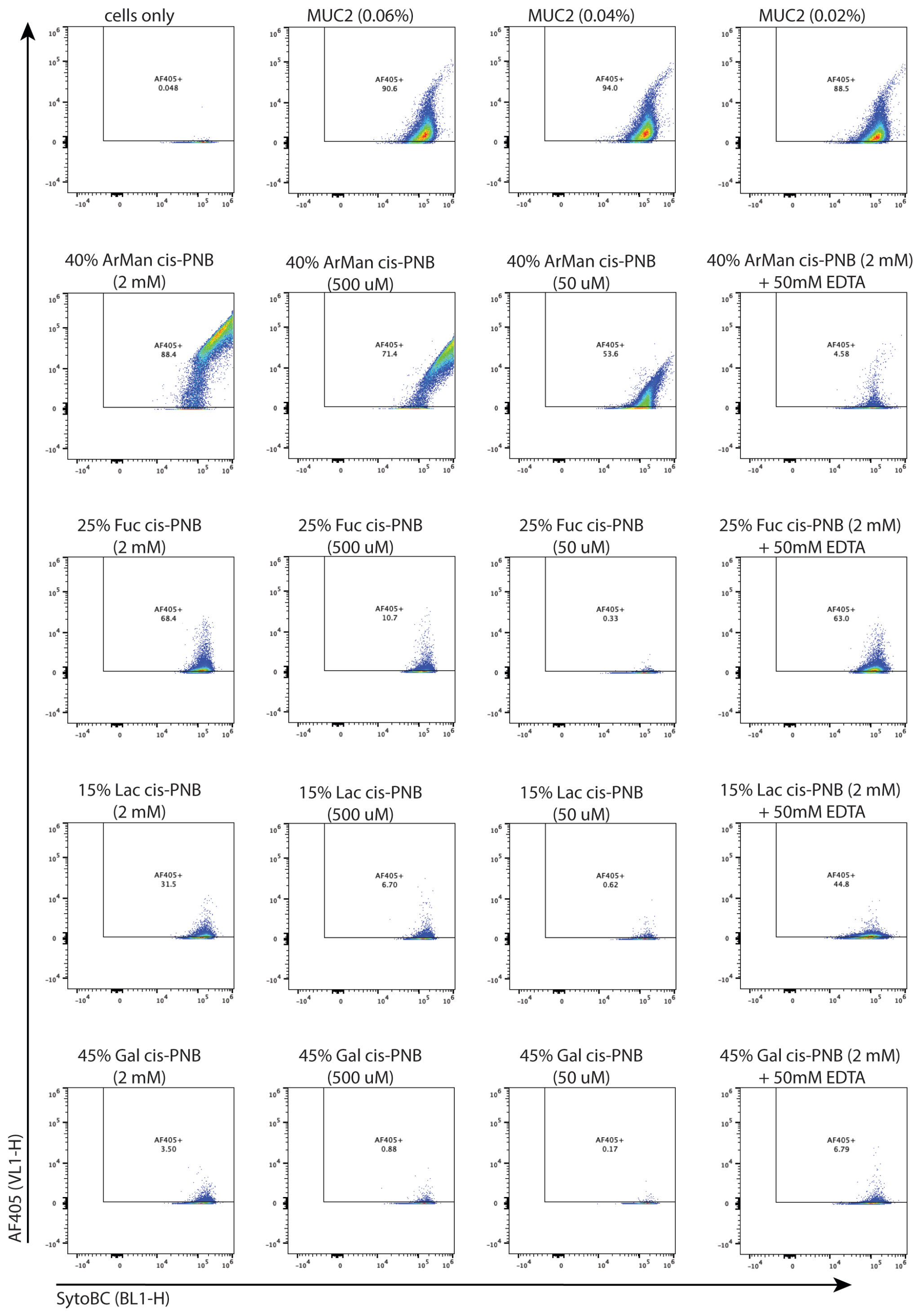
cells only
AF405+ 0.048
MUC2 (0.06%)
AF405+ 90.6
MUC2 (0.04%)
AF405+ 94.0
MUC2 (0.02%)
AF405+ 88.5
40% ArMan cis-PNB (2 mM)
AF405+ 88.4
40% ArMan cis-PNB (500 uM)
AF405+ 71.4
40% ArMan cis-PNB (50 uM)
AF405+ 53.6
40% ArMan cis-PNB (2 mM) + 50mM EDTA
AF405+ 4.58
25% Fuc cis-PNB (2 mM)
AF405+ 68.4
25% Fuc cis-PNB (500 uM)
AF405+ 10.7
25% Fuc cis-PNB (50 uM)
AF405+ 0.33
25% Fuc cis-PNB (2 mM) + 50mM EDTA
AF405+ 63.0
15% Lac cis-PNB (2 mM)
AF405+ 31.5
15% Lac cis-PNB (500 uM)
AF405+ 6.70
15% Lac cis-PNB (50 uM)
AF405+ 0.62
15% Lac cis-PNB (2 mM) + 50mM EDTA
AF405+ 44.8
45% Gal cis-PNB (2 mM)
AF405+ 3.50
45% Gal cis-PNB (500 uM)
AF405+ 0.88
45% Gal cis-PNB (50 uM)
AF405+ 0.17
45% Gal cis-PNB (2 mM) + 50mM EDTA
AF405+ 6.79
AF405 (VL1-H)
SytoBC (BL1-H)

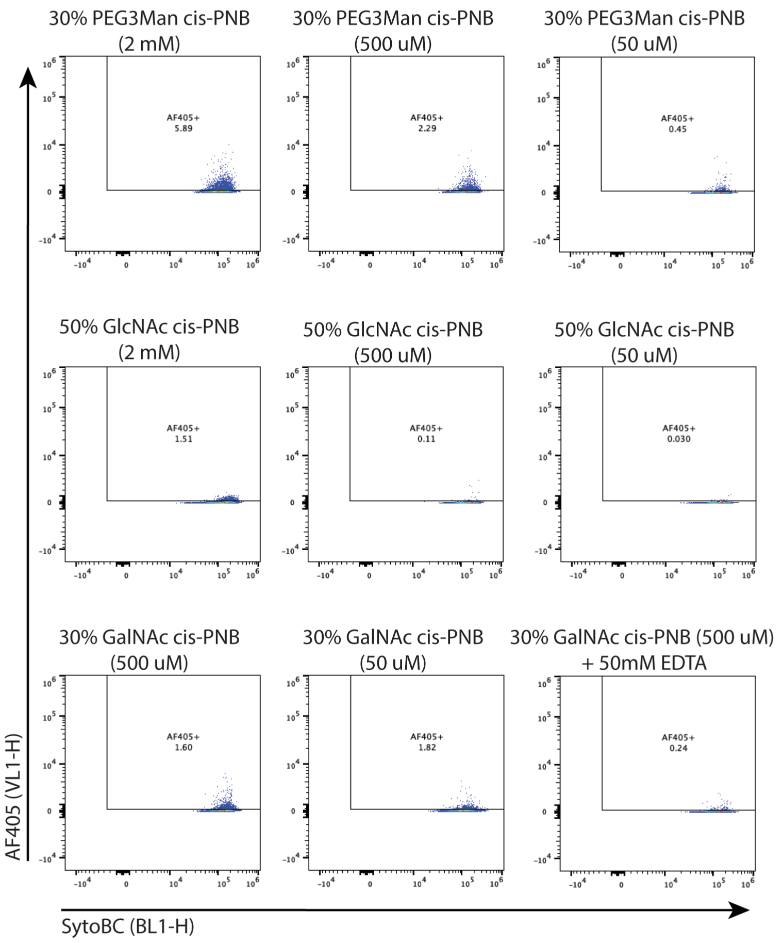
30% PEG3Man cis-PNB (2 mM)
AF405+ 5.89
30% PEG3Man cis-PNB (500 uM)
AF405+ 2.29
30% PEG3Man cis-PNB (50 uM)
AF405+ 0.45
50% GlcNAc cis-PNB (2 mM)
AF405+ 1.51
50% GlcNAc cis-PNB (500 uM)
AF405+ 0.11
50% GlcNAc cis-PNB (50 uM)
AF405+ 0.030
30% GalNAc cis-PNB (500 uM)
AF405+ 1.60
30% GalNAc cis-PNB (50 uM)
AF405+ 1.82
30% GalNAc cis-PNB (500 uM) + 50mM EDTA
AF405+ 0.24
AF405 (VL1-H)
SytoBC (BL1-H)

**Figure S8.** Flow cytometry with *L. reuteri* stationary phase and various glycopolymers at multiple concentrations. The ODs after overnight growth were 1.77, 1.496, 1.563, and 1.599. The assay has been performed in biological quadruplicate with technical triplicates in each plate.

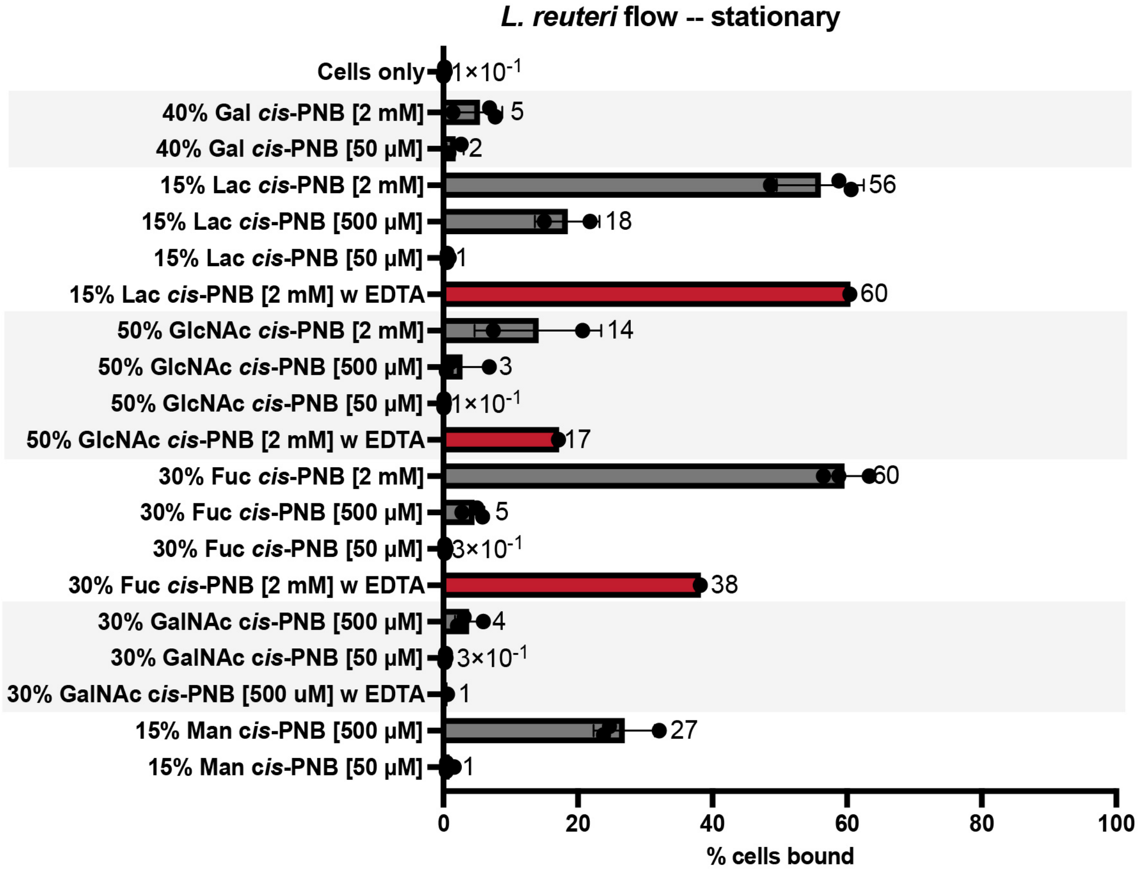

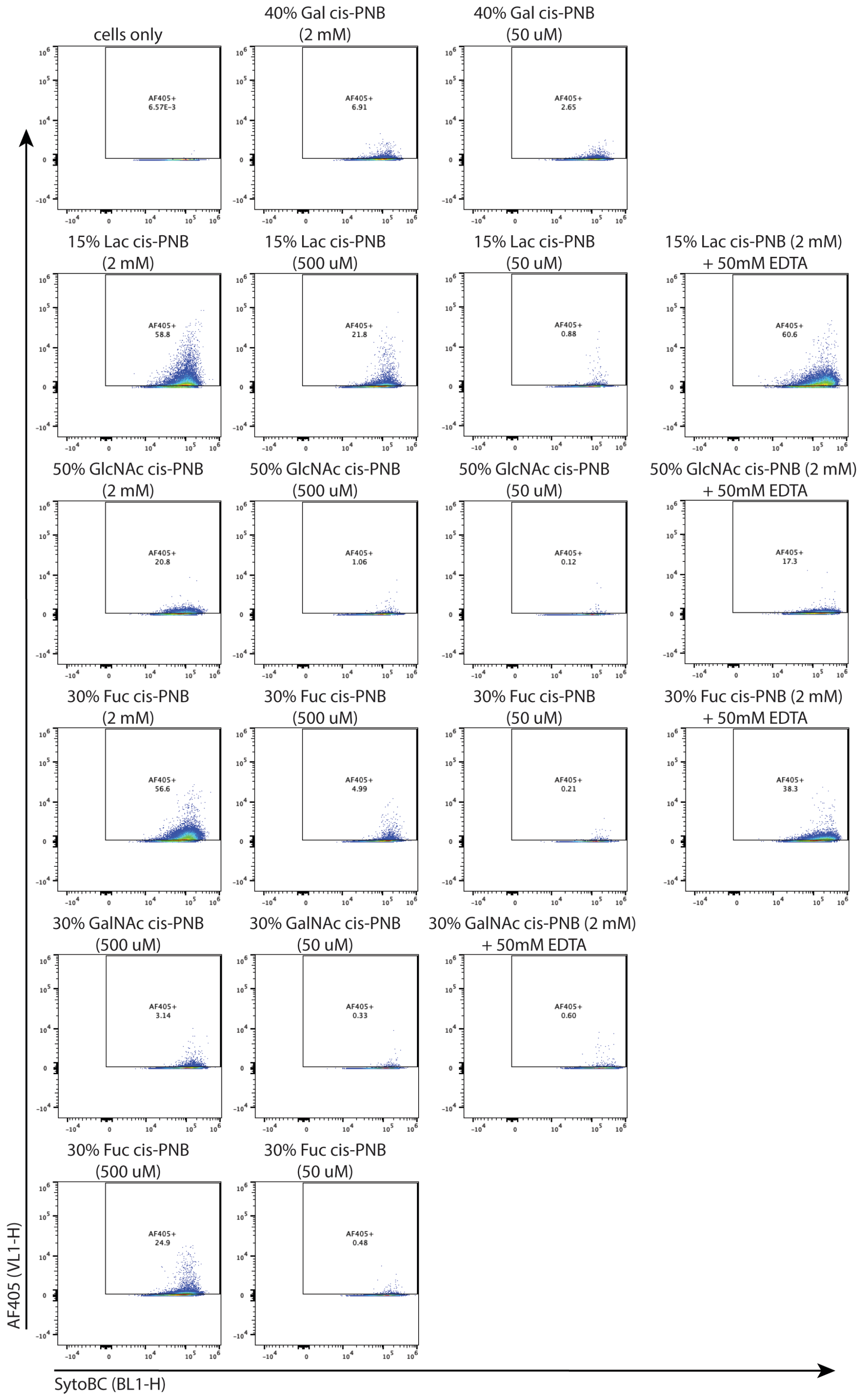
cells only
40% Gal cis-PNB (2 mM)
40% Gal cis-PNB (50 uM)
15% Lac cis-PNB (2 mM)
15% Lac cis-PNB (500 uM)
15% Lac cis-PNB (50 uM)
15% Lac cis-PNB (2 mM) + 50mM EDTA
50% GlcNAc cis-PNB (2 mM)
50% GlcNAc cis-PNB (500 uM)
50% GlcNAc cis-PNB (50 uM)
50% GlcNAc cis-PNB (2 mM) + 50mM EDTA
30% Fuc cis-PNB (2 mM)
30% Fuc cis-PNB (500 uM)
30% Fuc cis-PNB (50 uM)
30% Fuc cis-PNB (2 mM) + 50mM EDTA
30% GalNAc cis-PNB (500 uM)
30% GalNAc cis-PNB (50 uM)
30% GalNAc cis-PNB (2 mM) + 50mM EDTA
30% Fuc cis-PNB (500 uM)
30% Fuc cis-PNB (50 uM)
AF405 (VL1-H)
SytoBC (BL1-H)

**Figure S9.** Flow cytometry of *L. reuteri* in mid-log phase with various glycopolymer concentrations and glycan identities. The OD after overnight culture was 1.58, 1.56, and 1.67. The bacteria were allowed to recover for an hour in inoculations of 1 mL into 15 mL MRS media to OD = 0.44, 0.44, and 0.48. The assay was performed in biological triplicate with technical triplicates in each plate.

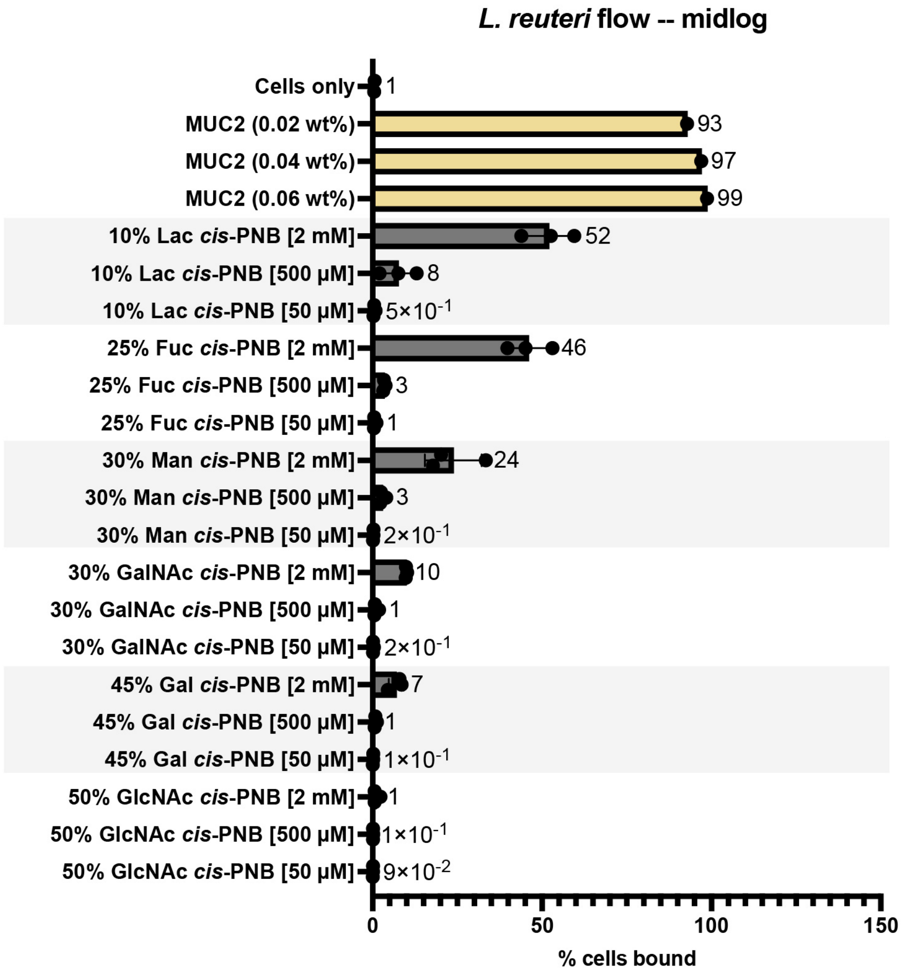

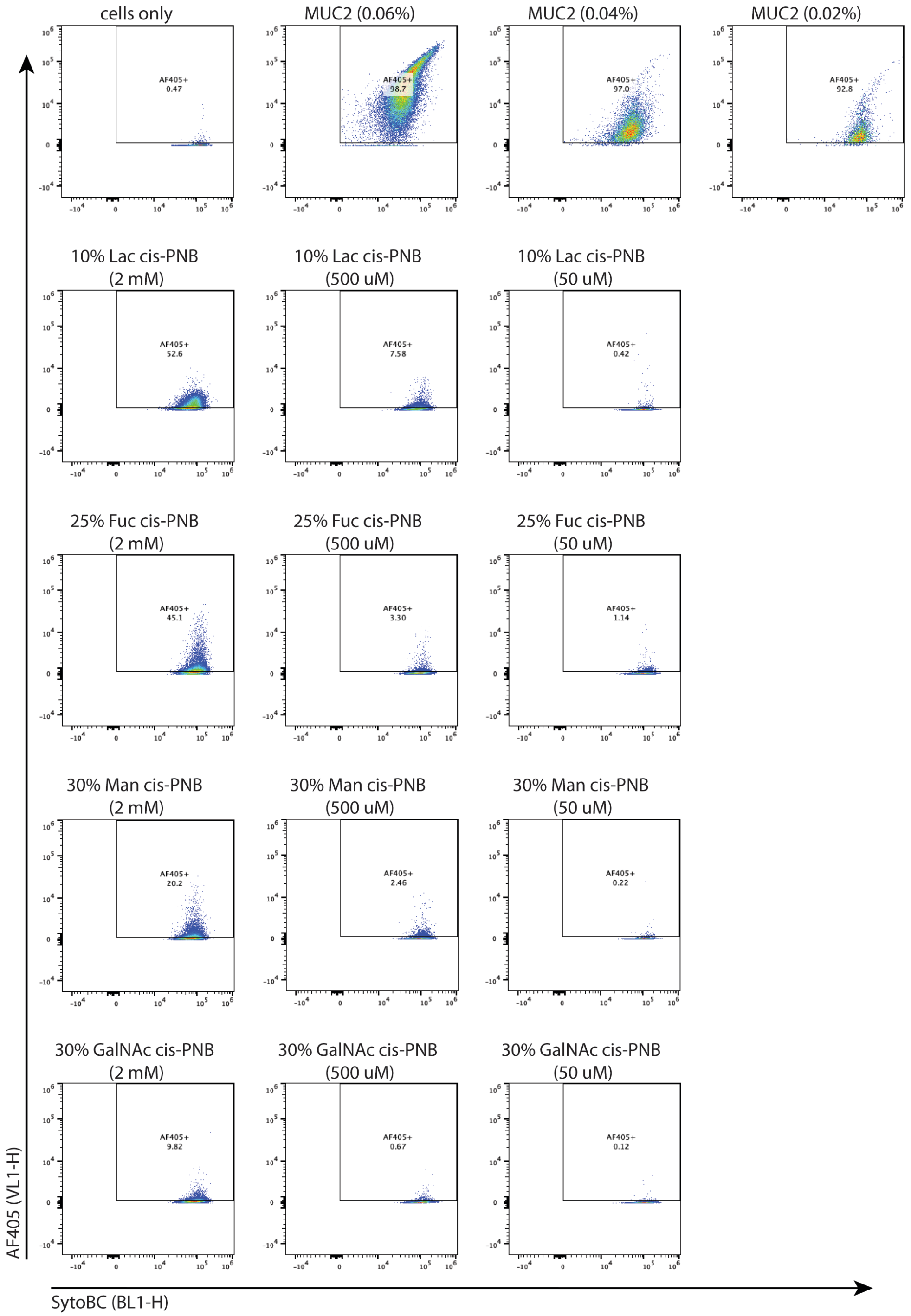
cells only
AF405+ 0.47
MUC2 (0.06%)
AF405+ 98.7
MUC2 (0.04%)
AF405+ 97.0
MUC2 (0.02%)
AF405+ 92.8
10% Lac cis-PNB (2 mM)
AF405+ 52.6
10% Lac cis-PNB (500 uM)
AF405+ 7.58
10% Lac cis-PNB (50 uM)
AF405+ 0.42
25% Fuc cis-PNB (2 mM)
AF405+ 45.1
25% Fuc cis-PNB (500 uM)
AF405+ 3.30
25% Fuc cis-PNB (50 uM)
AF405+ 1.14
30% Man cis-PNB (2 mM)
AF405+ 20.2
30% Man cis-PNB (500 uM)
AF405+ 2.46
30% Man cis-PNB (50 uM)
AF405+ 0.22
30% GalNAc cis-PNB (2 mM)
AF405+ 9.82
30% GalNAc cis-PNB (500 uM)
AF405+ 0.67
30% GalNAc cis-PNB (50 uM)
AF405+ 0.12
AF405 (VL1-H)
SytoBC (BL1-H)

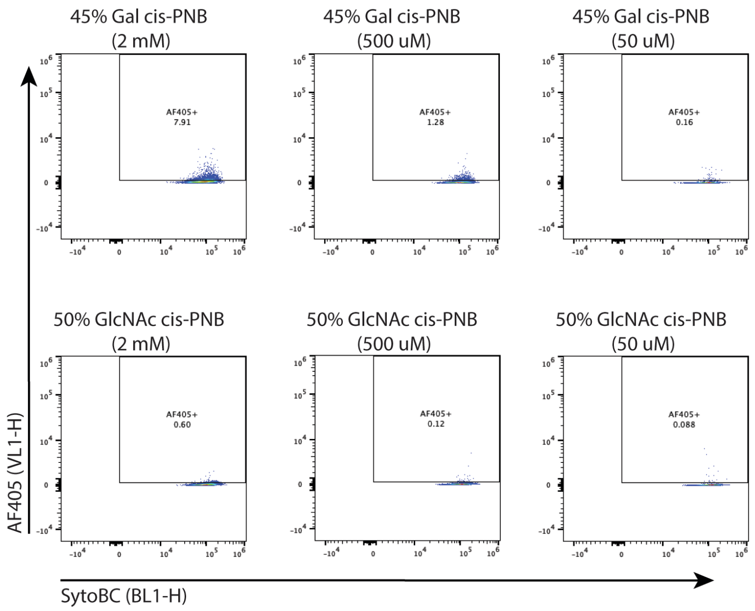


**Table S1.** Determination of minimal media for *L. reuteri* growth.

| Bacteria | Media | OD at 24 h | OD at 48 h |
|---|---|---|---|
| *L. reuteri* | MRS | 1.75 | 1.83 |
| *L. reuteri* | M9 | -0.001 | -0.001 |
| ***L. reuteri*** | **MOD-MRS** | **0.122** | **N/A** |
| *L. plantarum* | MOD-MRS | 0.051 | 0.047 |
| *L. fermentum* | MOD-MRS | 0.053 | N/A |
| *L. reuteri* | MOD-MRS without peptone or extracts | 0.001 | N/A |
| *L. plantarum* | MOD-MRS without peptone or extracts | 0.002 | N/A |
| *L. fermentum* | MOD-MRS without peptone or extracts | 0.001 | N/A |

**Figure S10.** Flow cytometry with *L. reuteri* after growth in minimal media with various glycopolymer concentrations and glycan identities. After overnight growth in MOD-MRS anaerobically, the ODs were measured as 0.122, 0.146, and 0.146. The assay was performed in biological triplicate with technical triplicates in each plate.

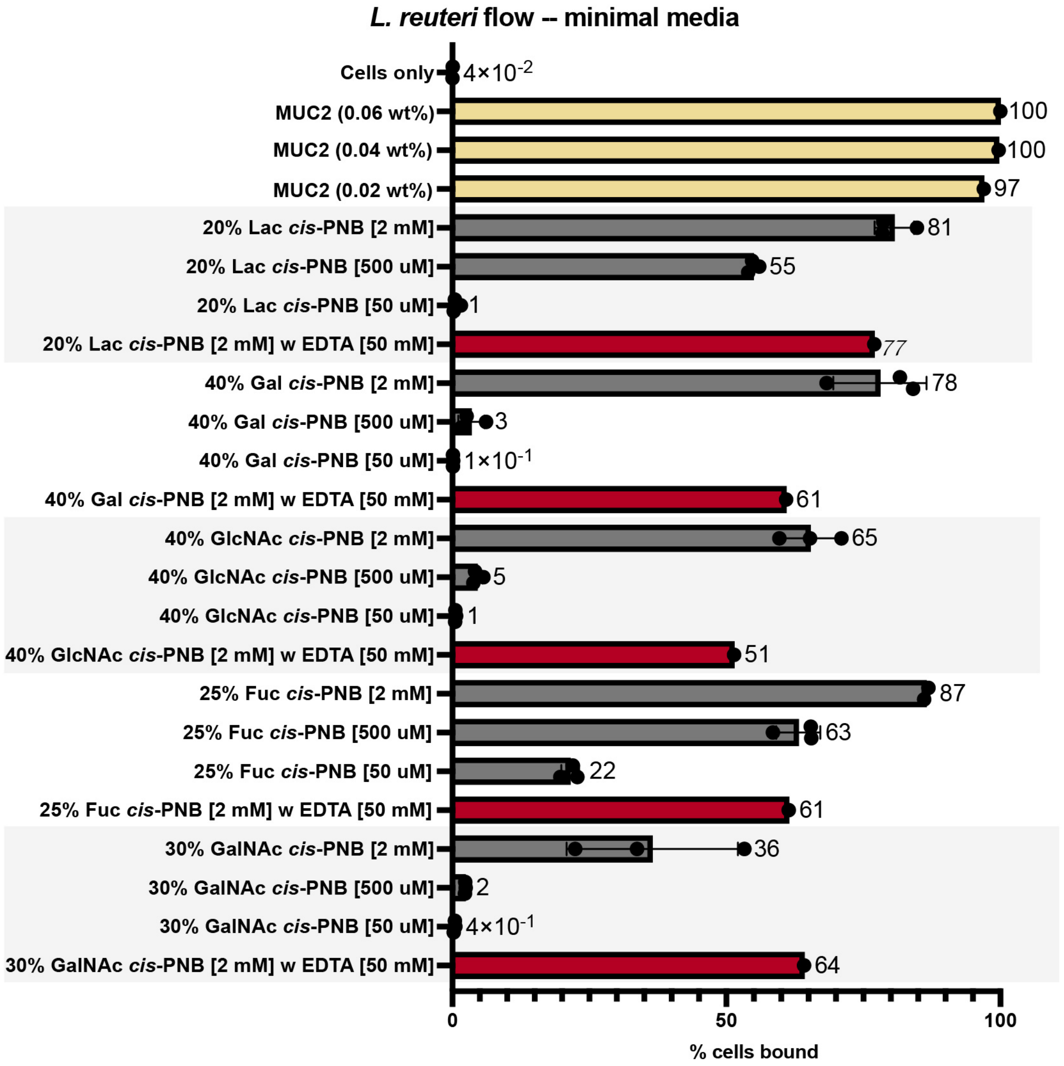

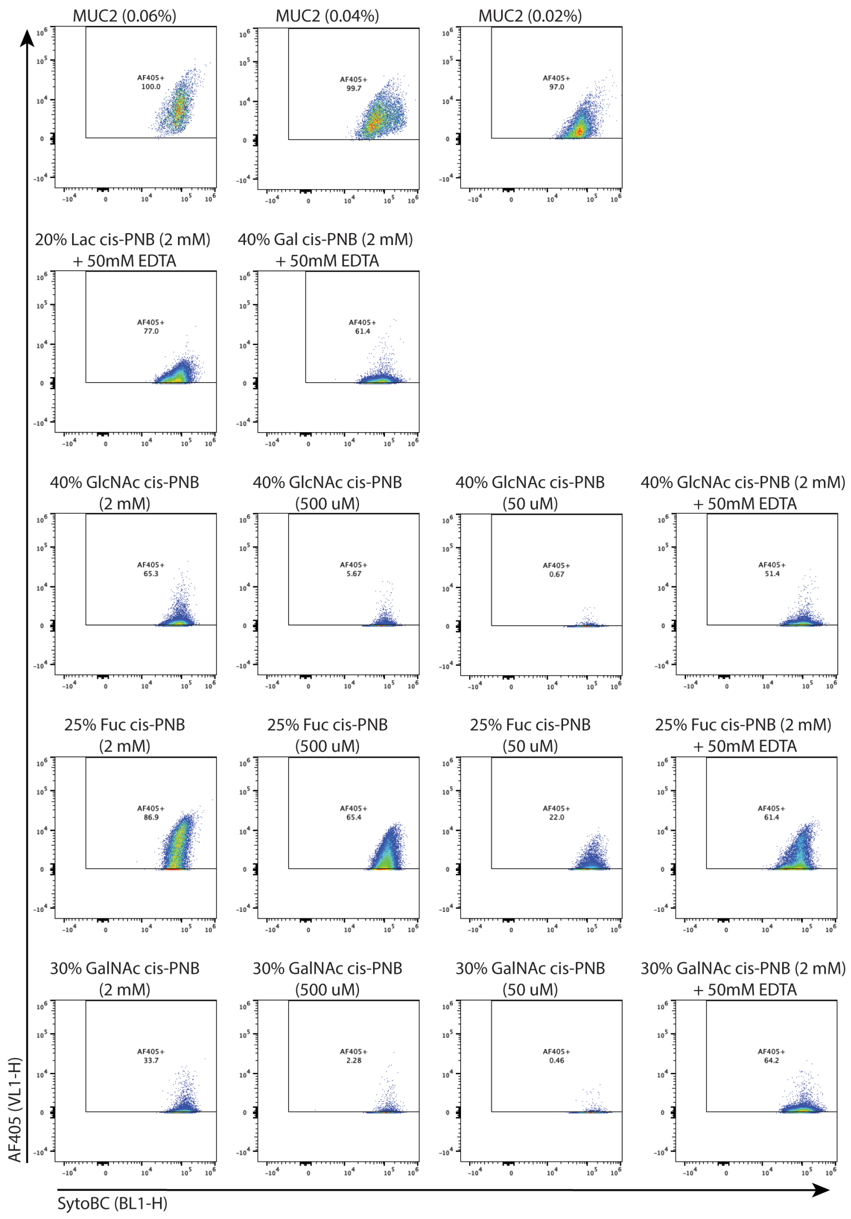
MUC2 (0.06%)
MUC2 (0.04%)
MUC2 (0.02%)
AF405+ 100.0
AF405+ 99.7
AF405+ 97.0
20% Lac cis-PNB (2 mM) + 50mM EDTA
40% Gal cis-PNB (2 mM) + 50mM EDTA
AF405+ 77.0
AF405+ 61.4
40% GlcNAc cis-PNB (2 mM)
40% GlcNAc cis-PNB (500 uM)
40% GlcNAc cis-PNB (50 uM)
40% GlcNAc cis-PNB (2 mM) + 50mM EDTA
AF405+ 65.3
AF405+ 5.67
AF405+ 0.67
AF405+ 51.4
25% Fuc cis-PNB (2 mM)
25% Fuc cis-PNB (500 uM)
25% Fuc cis-PNB (50 uM)
25% Fuc cis-PNB (2 mM) + 50mM EDTA
AF405+ 86.9
AF405+ 65.4
AF405+ 22.0
AF405+ 61.4
30% GalNAc cis-PNB (2 mM)
30% GalNAc cis-PNB (500 uM)
30% GalNAc cis-PNB (50 uM)
30% GalNAc cis-PNB (2 mM) + 50mM EDTA
AF405+ 33.7
AF405+ 2.28
AF405+ 0.46
AF405+ 64.2
AF405 (VL1-H)
SytoBC (BL1-H)

**Figure S11.** Microscopy with *L. reuteri* in stationary phase. The images were collected in technical triplicate and biological triplicate, with a dilution of 5:95 post-flow cytometry.

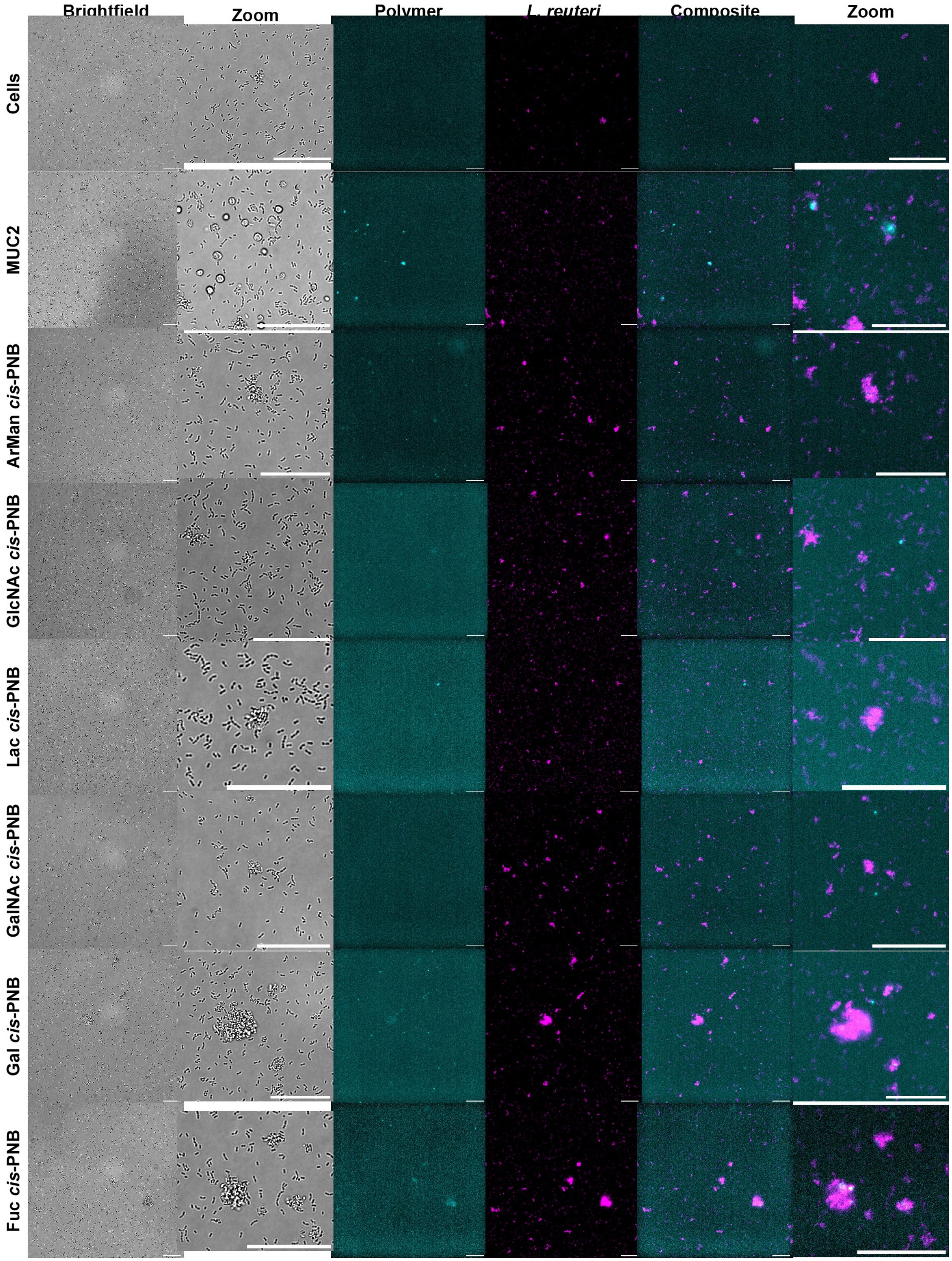

**Figure S12.** *L. fermentum* mid-log phase with multiple concentrations. OD = 1.62, 1.58, and 1.64 after overnight growth. The bacteria was allowed to recover for one hour after inoculating of 1 mL culture into 15 mL media. OD = 0.62, 0.60, and 0.56. Performed in triplicate.

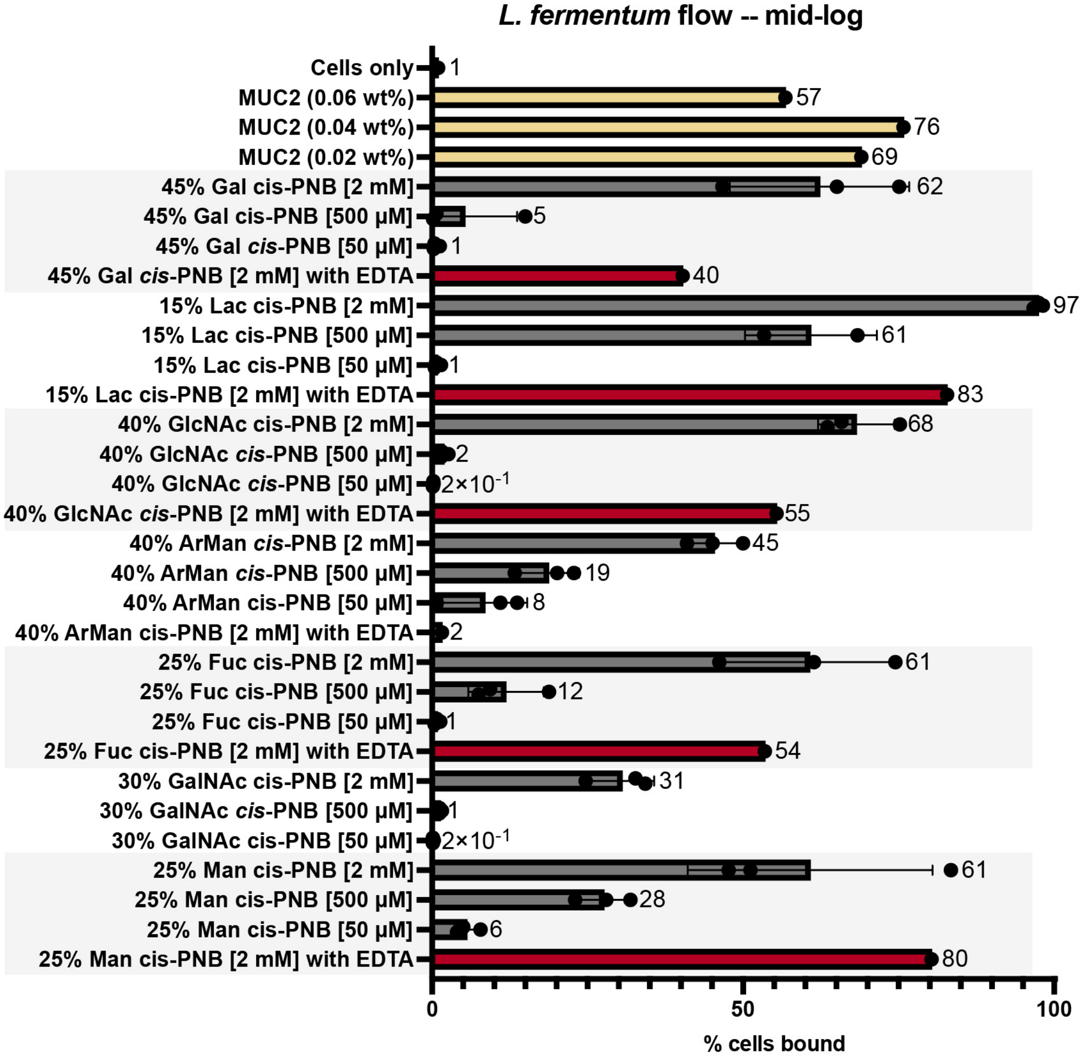

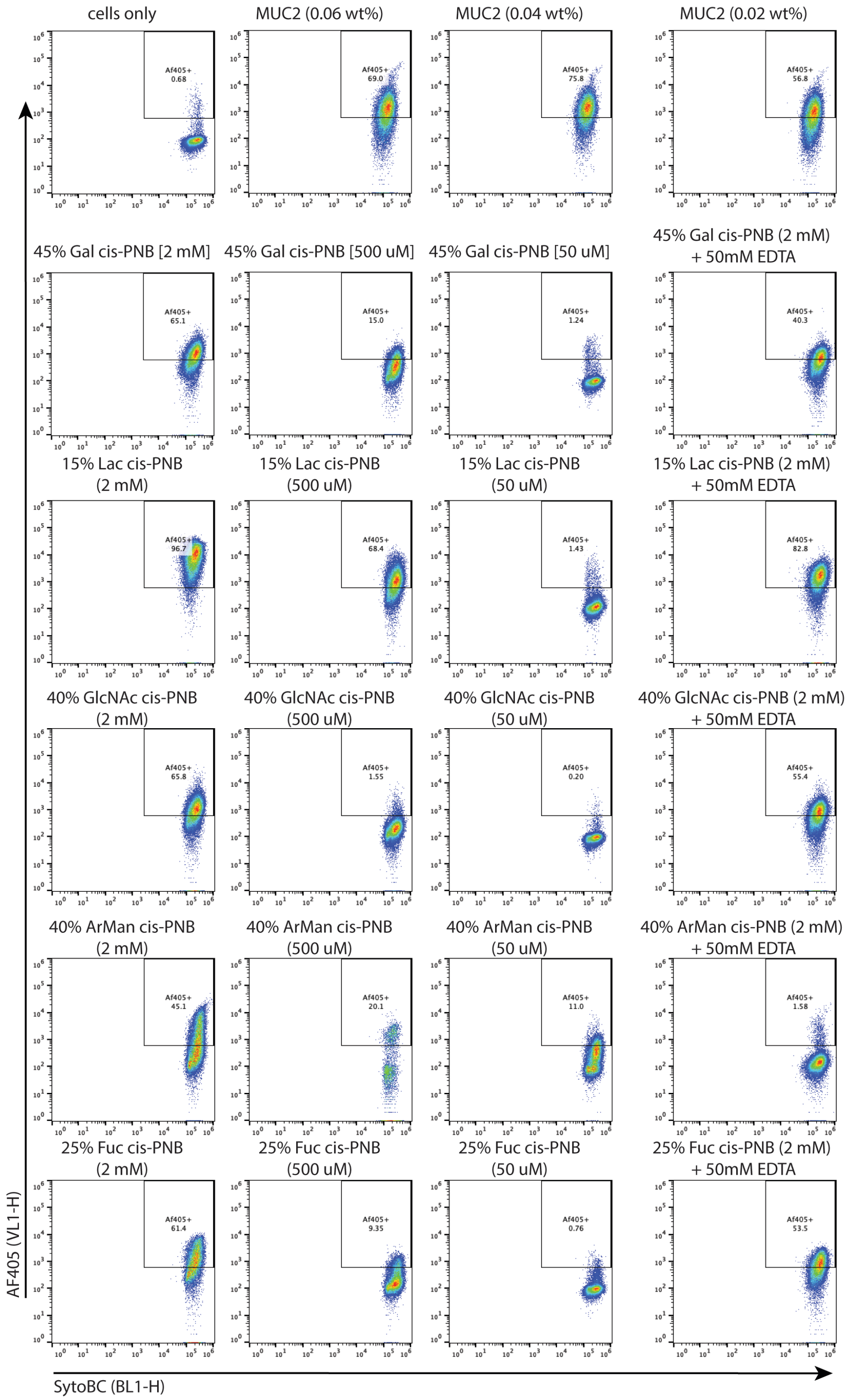
cells only
MUC2 (0.06 wt%)
MUC2 (0.04 wt%)
MUC2 (0.02 wt%)
Af405+ 0.68
Af405+ 69.0
Af405+ 75.8
Af405+ 56.8
45% Gal cis-PNB [2 mM]
45% Gal cis-PNB [500 uM]
45% Gal cis-PNB [50 uM]
45% Gal cis-PNB (2 mM) + 50mM EDTA
Af405+ 65.1
Af405+ 15.0
Af405+ 1.24
Af405+ 40.3
15% Lac cis-PNB (2 mM)
15% Lac cis-PNB (500 uM)
15% Lac cis-PNB (50 uM)
15% Lac cis-PNB (2 mM) + 50mM EDTA
Af405+ 96.7
Af405+ 68.4
Af405+ 1.43
Af405+ 82.8
40% GlcNAc cis-PNB (2 mM)
40% GlcNAc cis-PNB (500 uM)
40% GlcNAc cis-PNB (50 uM)
40% GlcNAc cis-PNB (2 mM) + 50mM EDTA
Af405+ 65.8
Af405+ 1.55
Af405+ 0.20
Af405+ 55.4
40% ArMan cis-PNB (2 mM)
40% ArMan cis-PNB (500 uM)
40% ArMan cis-PNB (50 uM)
40% ArMan cis-PNB (2 mM) + 50mM EDTA
Af405+ 45.1
Af405+ 20.1
Af405+ 11.0
Af405+ 1.58
25% Fuc cis-PNB (2 mM)
25% Fuc cis-PNB (500 uM)
25% Fuc cis-PNB (50 uM)
25% Fuc cis-PNB (2 mM) + 50mM EDTA
Af405+ 61.4
Af405+ 9.35
Af405+ 0.76
Af405+ 53.5
AF405 (VL1-H)
SytoBC (BL1-H)

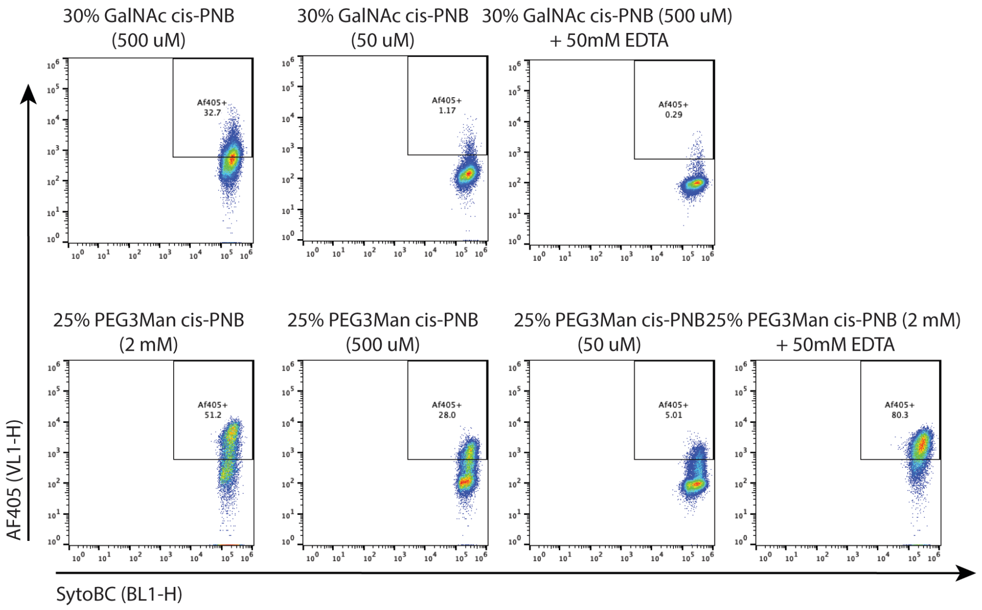

30% GalNAc cis-PNB (500 uM)
30% GalNAc cis-PNB (50 uM)
30% GalNAc cis-PNB (500 uM) + 50mM EDTA
Af405+ 32.7
Af405+ 1.17
Af405+ 0.29
25% PEG3Man cis-PNB (2 mM)
25% PEG3Man cis-PNB (500 uM)
25% PEG3Man cis-PNB (50 uM)
25% PEG3Man cis-PNB (2 mM) + 50mM EDTA
Af405+ 51.2
Af405+ 28.0
Af405+ 5.01
Af405+ 80.3
AF405 (VL1-H)
SytoBC (BL1-H)

**Figure S13.** Flow cytometry with *L. fermentum* in stationary phase with various glycopolymer concentrations and glycan identities. The ODs were 1.845, 1.826, and 1.725 after overnight growth. The assay was performed in biological triplicate with technical triplicates in each plate.

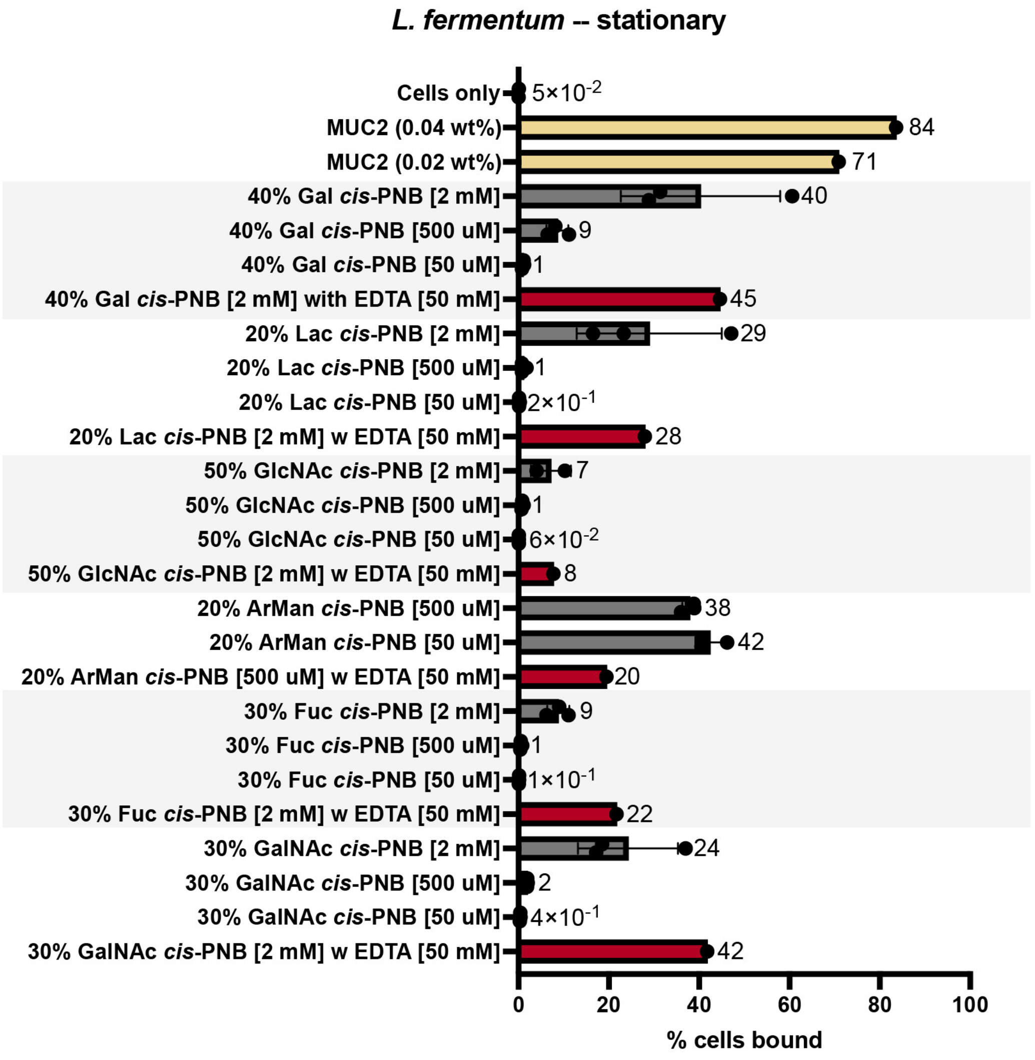

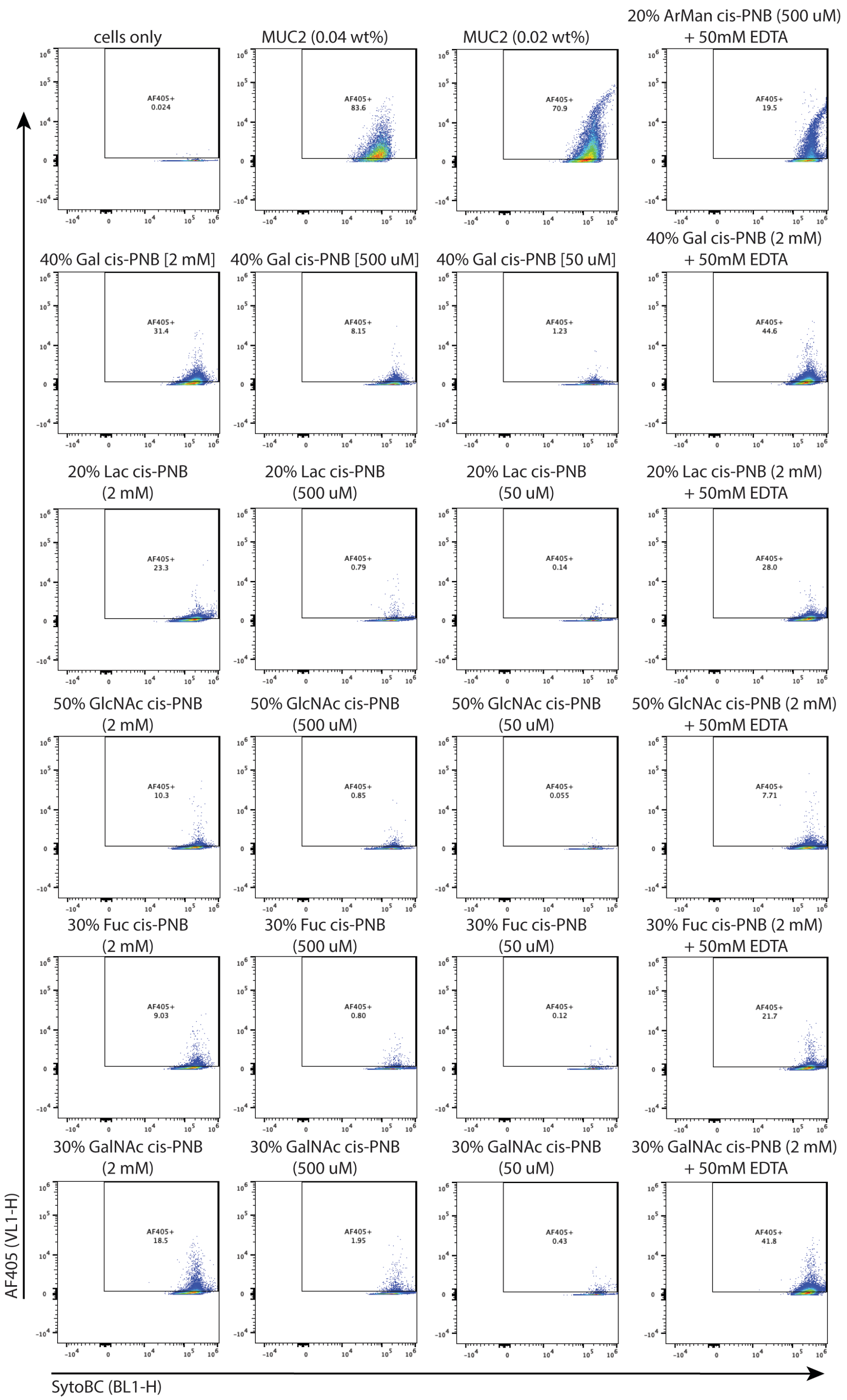
cells only
AF405+ 0.024
MUC2 (0.04 wt%)
AF405+ 83.6
MUC2 (0.02 wt%)
AF405+ 70.9
20% ArMan cis-PNB (500 uM) + 50mM EDTA
AF405+ 19.5
40% Gal cis-PNB [2 mM]
AF405+ 31.4
40% Gal cis-PNB [500 uM]
AF405+ 8.15
40% Gal cis-PNB [50 uM]
AF405+ 1.23
40% Gal cis-PNB (2 mM) + 50mM EDTA
AF405+ 44.6
20% Lac cis-PNB (2 mM)
AF405+ 23.3
20% Lac cis-PNB (500 uM)
AF405+ 0.79
20% Lac cis-PNB (50 uM)
AF405+ 0.14
20% Lac cis-PNB (2 mM) + 50mM EDTA
AF405+ 28.0
50% GlcNAc cis-PNB (2 mM)
AF405+ 10.3
50% GlcNAc cis-PNB (500 uM)
AF405+ 0.85
50% GlcNAc cis-PNB (50 uM)
AF405+ 0.055
50% GlcNAc cis-PNB (2 mM) + 50mM EDTA
AF405+ 7.71
30% Fuc cis-PNB (2 mM)
AF405+ 9.03
30% Fuc cis-PNB (500 uM)
AF405+ 0.80
30% Fuc cis-PNB (50 uM)
AF405+ 0.12
30% Fuc cis-PNB (2 mM) + 50mM EDTA
AF405+ 21.7
30% GalNAc cis-PNB (2 mM)
AF405+ 18.5
30% GalNAc cis-PNB (500 uM)
AF405+ 1.95
30% GalNAc cis-PNB (50 uM)
AF405+ 0.43
30% GalNAc cis-PNB (2 mM) + 50mM EDTA
AF405+ 41.8
AF405 (VL1-H)
SytoBC (BL1-H)

**Figure S14.** After stationary flow cytometry was assessed, the polymer and *L. fermentum* samples were analyzed by microscopy. Two images were taken per well for technical duplicate and all three biological replicates were imaged, with a dilution of 5:95 post-flow cytometry

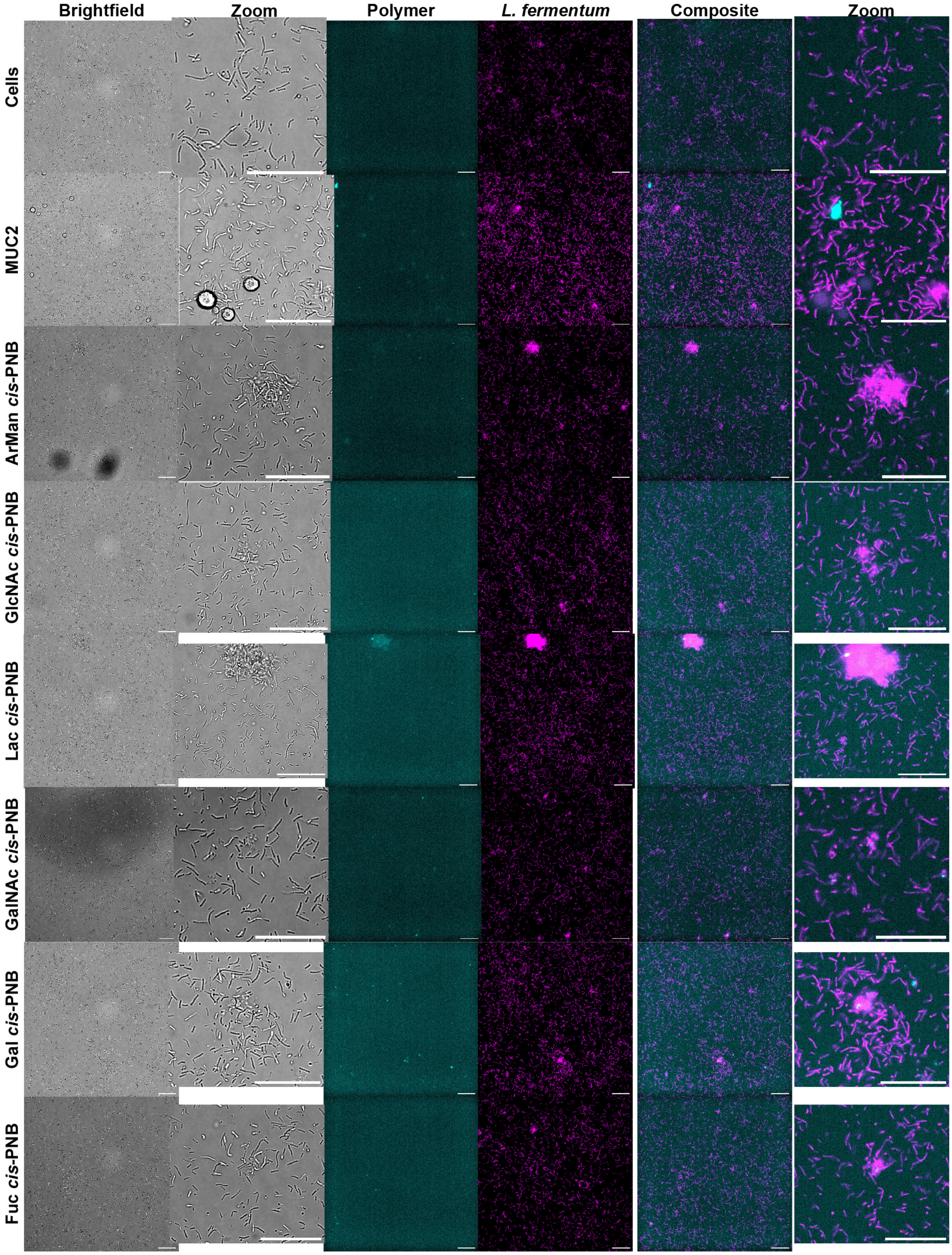

**Figure S15.** Clustering of *L. plantarum* cells with glycopolymer and native mucins in biological triplicate with technical duplicates. Native mucin, fucosylated polymer, galactosylated polymer, and aryl mannosylated polymer clustered the cells best.

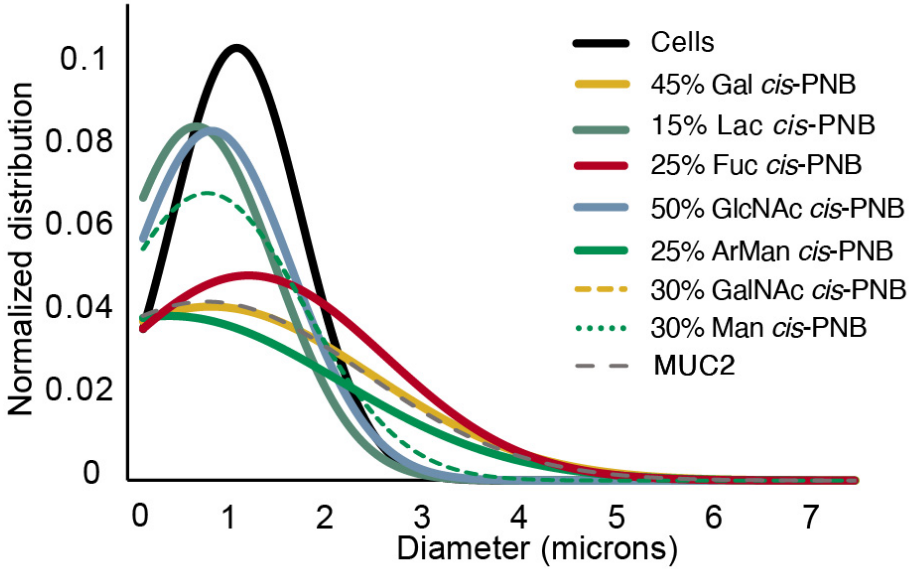


**Figure S16.** Clustering of *L. reuteri* cells with glycopolymer and native mucins in biological quadruplicate with technical duplicates. Native mucin, lactosylated polymer, and fucosylated polymer clustered the cells best. Cell counts ranged from 300 to 10,000 per image with one image per curve.

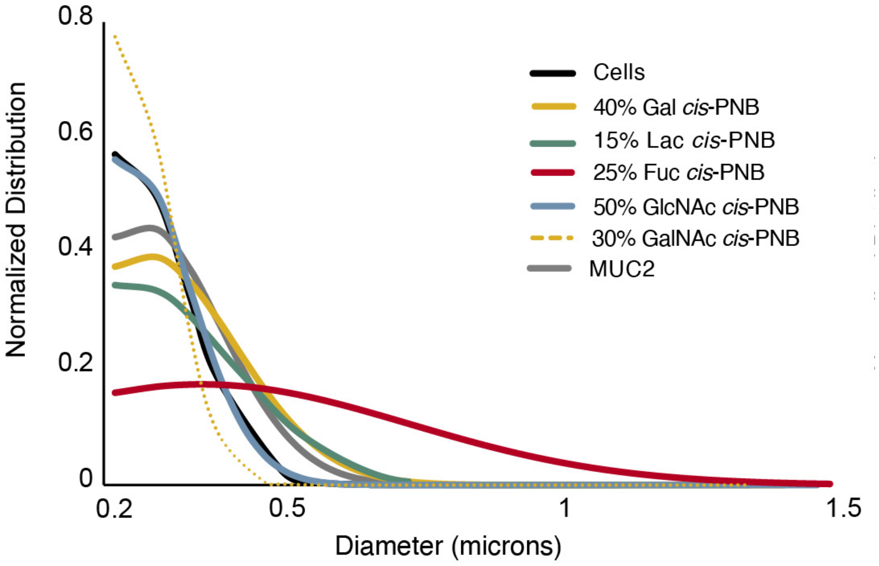

**Figure S17.** Clustering of *L. fermentum* cells with glycopolymer and native mucins in biological triplicate with technical duplicates. Galactosylated polymer, fucosylated polymer, and aryl mannosylated polymer clustered the cells best. Cell counts per image ranged from 1,000 to 20,000 with one image per curve.

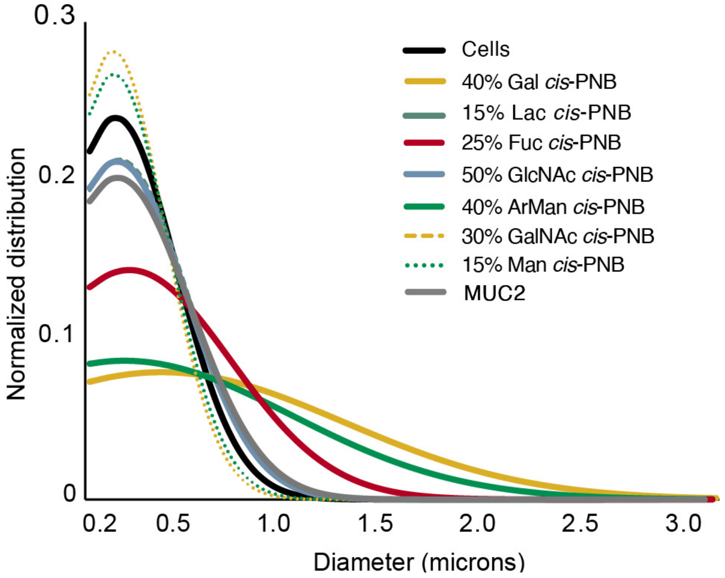


**Figure S18.** Confirmation of staining efficacy using staining via esterases without crosstalk of the AF405 fluorophore on the polymer. Based on this data, 100 µM cFDA was used to stain the cells for adhesion. Additionally, the loading of MUC2-NHS was evaluated and determined that incubated the amine-coated plate with 30% MUC2-NHS was best for *L. reuteri* commensal retainment.

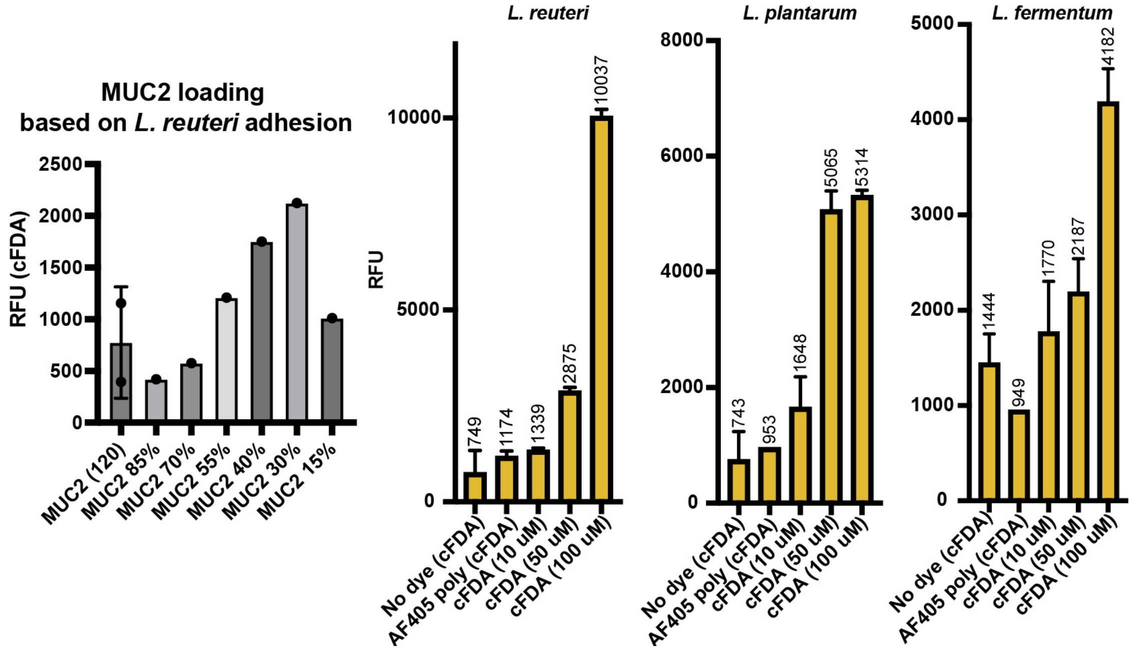

**Figure S19.** Commensal adhesion of *L. fermentum* to MUC2-coated wells in the presence and absence of synthetic mucins. The assay was performed in biological duplicate with technical replicates (duplicate or triplicate) in each plate. Significance was calculated using an ANOVA Dunnett test to compare the mean of the MUC2-only well (yellow) relative to the glycopolymer additive wells (dark grey). Relative commensal populations are shown on the left by cFDA fluorescence. Glycopolymer presence is shown on the right by AF405 fluorescence. While the cFDA fluorescence should be considered quantitative, the AF405 fluorescence should be considered qualitative.

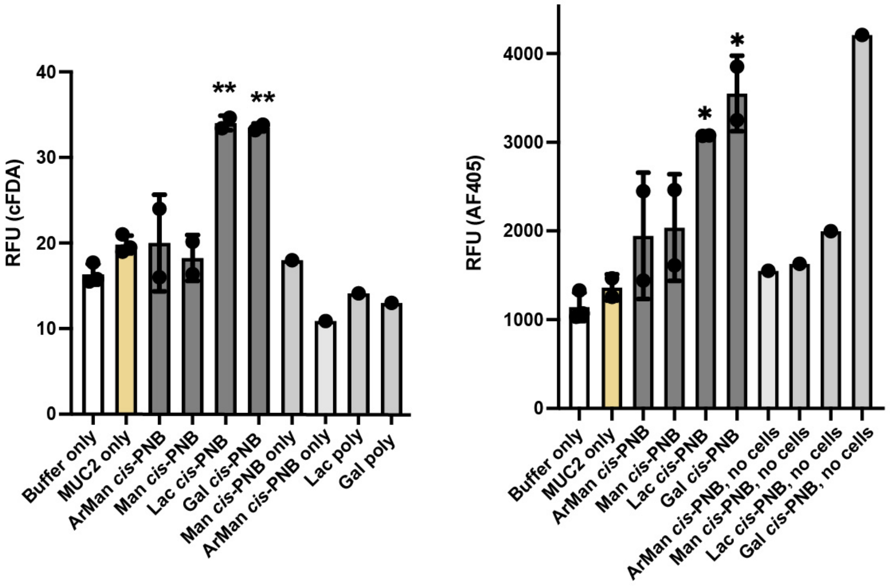

**Figure S20**. *L. fermentum* was grown in MRS broth overnight at 37 °C aerobically. OD600 was taken to ensure the bacteria had reached saturation prior to dilution in MOD-MRS to OD = 0.1. The cells were added as 10 µL to each well of a clear 96-well plate and further diluted with MOD-MRS and glycan (10 µL in MilliQ water). Final concentrations were: 0.2 mM for monosaccharides, disaccharides, for synthetic mucin [with respect to glycan], and 0.1 mg/mL for MUC2 with 100 µL total volume. The assays included a buffer-only well and cells-only well as negative controls. Absorbance at 600 nm was measured by a SpectraMax M5 plate reader over 14 hours with readings every 15 minutes. The plate reader was kept at 37 °C and the plate was sealed with a gas-permeable sticker. The results represent technical quadruplicates with six biological replicates. ANOVA tests show significance with respect to the cells-only control growth.

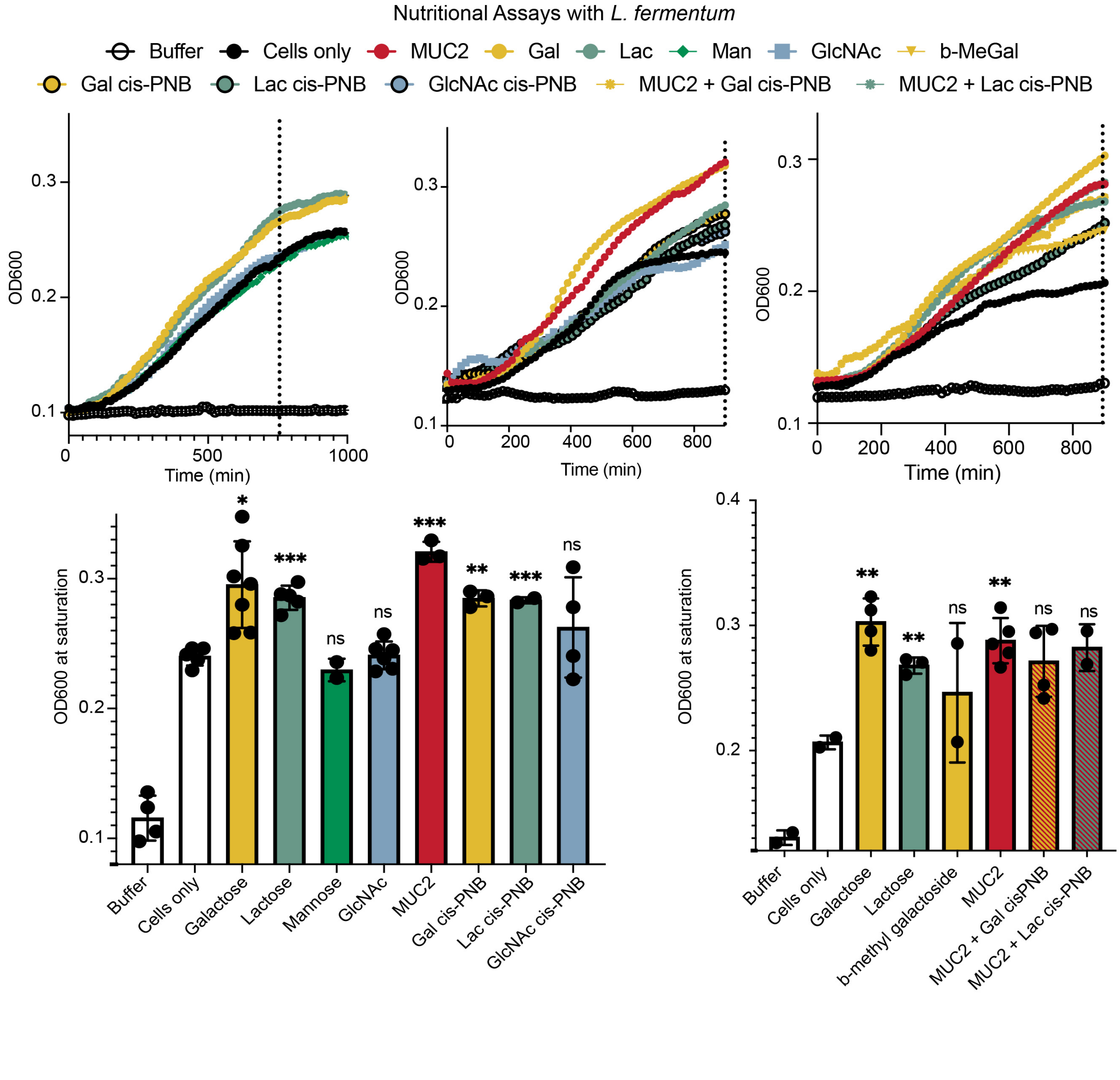

**Figure S21.** 4-methylumbelliferyl glycans were procured for β-glucose, N-acetyl -β-D-glucosamine, a-mannose, β-galactose, and a-fucose (Fisher Scientific). The 4MU-glycans were dissolved in 10% DMSO/90% MilliQ water at [40 mM]. The bacteria were grown anaerobically or aerobically in MRS broth to saturation overnight at 37 °C. The bacteria were pelleted and transferred to MOD-MRS or HEPES with added calcium at an OD = 1. The final concentrations within the black 96-well plate were cells [OD = 0.1] and 4MU-Gly [4 mM] at a total volume of 100 μL. Negative controls included buffer-only, cells-only, and 4MU-Gly-only wells. The plate was read by a SpectraMax M5 plate reader for excitation at 360 nm and emission at 448 nm over

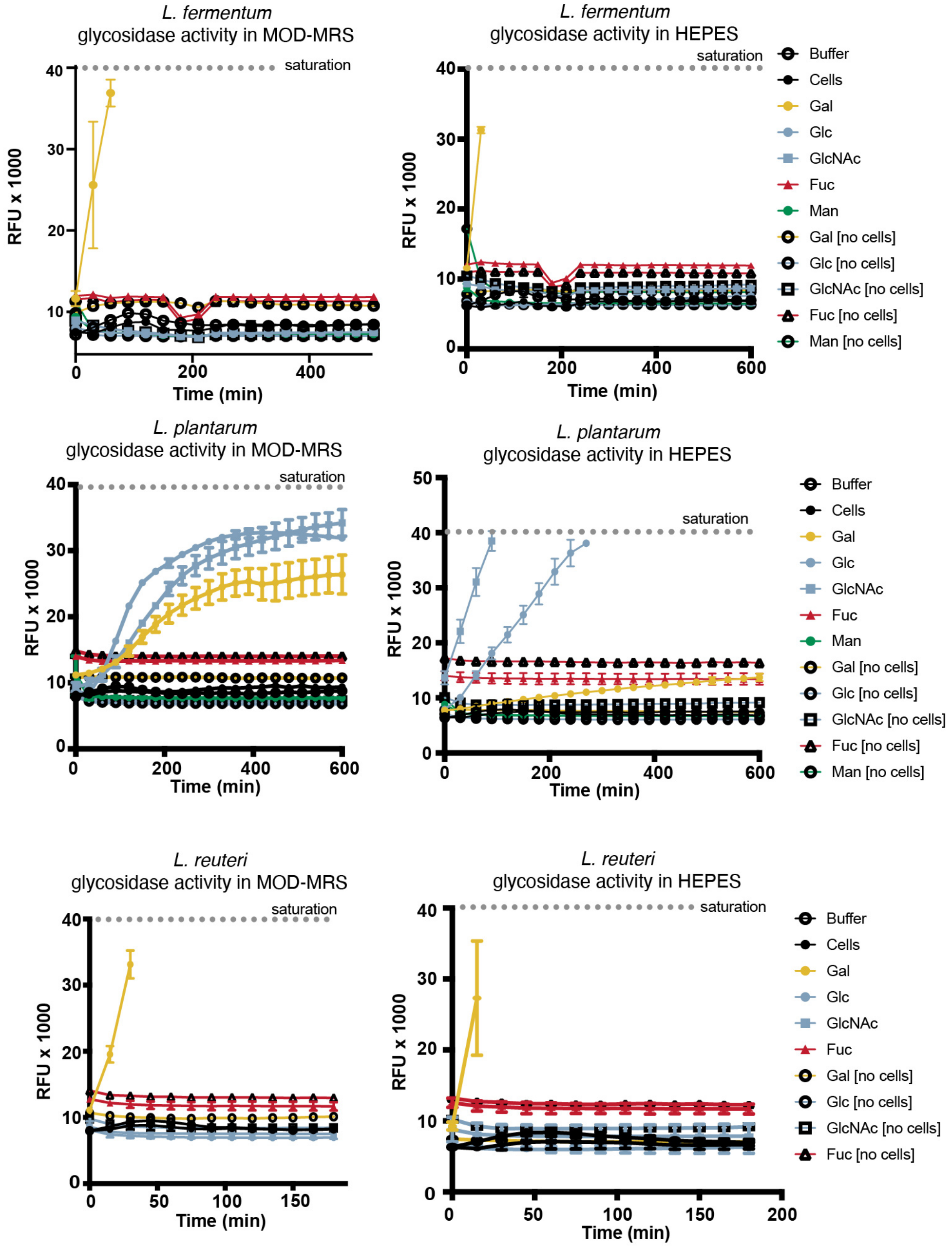


10+ hours. Saturation of the plate reader was observed if fluorescence was >40,000 a.u. The assay was completed once for each bacteria with technical triplicates within each plate.

**Table S2.** Real time qPCR primer sequence

| Gene | Gene ID | Forward Primer | Reverse Primer |
|---|---|---|---|
| *L. fermentum* beta-galactosidase small subunit | 83715431 | ACGTGGAAGGGTTGCTGGTG | CTGGTGTACGGCAAGCACGA |

**Figure S22.** Quantitative real-time PCR analysis of galactosidase expression in *L. fermentum* evaluating the impact of media conditions. Growth conditions were compared for *L. fermentum* grown overnight in nutrient-rich media (MRS broth, OD = 0.775) and *L. fermentum* grown in MUC2-enhanced nutrient-rich media (MRS broth with MUC2 at 1 mg/15 mL, OD = 1.069). Galactosidase expression is normalized against the 16S gene. The error bars represent standard error of the mean. Welch's t-test was used for data analysis (**P* < 0.05, ** *P* < 0.01).

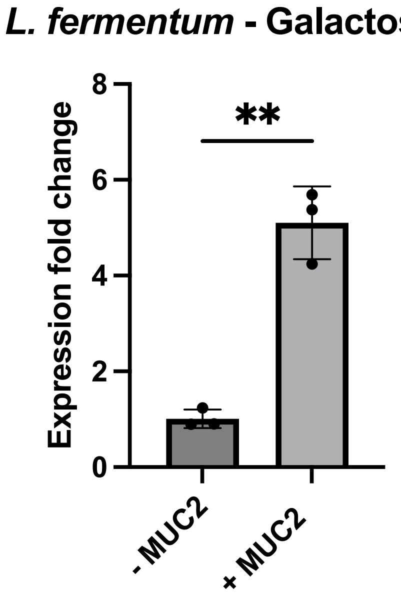

**Figure S23.** Quantitative real-time PCR analysis of galactosidase expression in *L. fermentum* evaluating the impact of media conditions. Growth conditions were compared for *L. fermentum* grown overnight in nutrient-rich media (MRS broth, OD = 1.41) and *L. fermentum* grown in synthetic mucin-enhanced nutrient-rich media (MRS broth with 30% Gal cis-PNB at 72 µM, OD = 1.24 or MRS broth with 15% Lac *cis*-PNB at 72 µM, OD = 1.07). Galactosidase expression is normalized against the 16S gene. The error bars represent standard error of the mean. One-way ANOVA with Dunnett's multiple comparisons test was used for data analysis (*$P < 0.05$, ** $P < 0.01$).

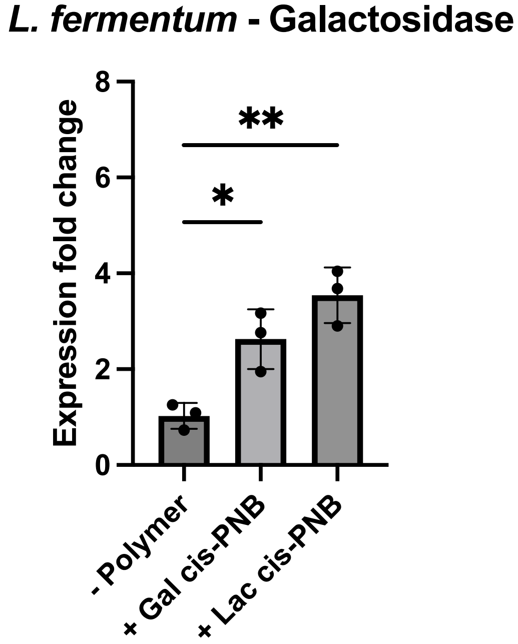


### 3. Ligand and Polymer Syntheses

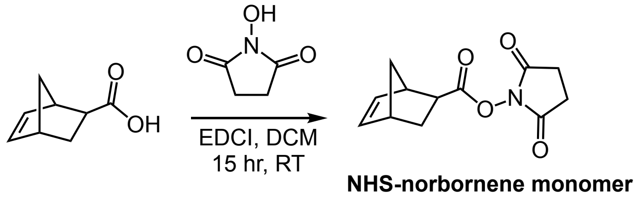


***N*-hydroxysuccinimidyl norbornene monomer** was synthesized according to known procedures.[3,4]

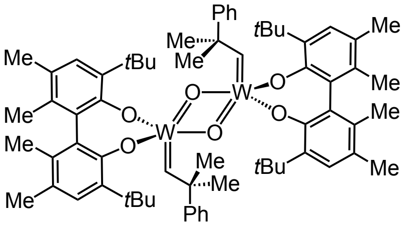


[W] catalyst

**Schrock *cis*-ROMP catalyst**: Catalyst was synthesized according to the procedure for compound *rac*-2a in the cited reference.[5]

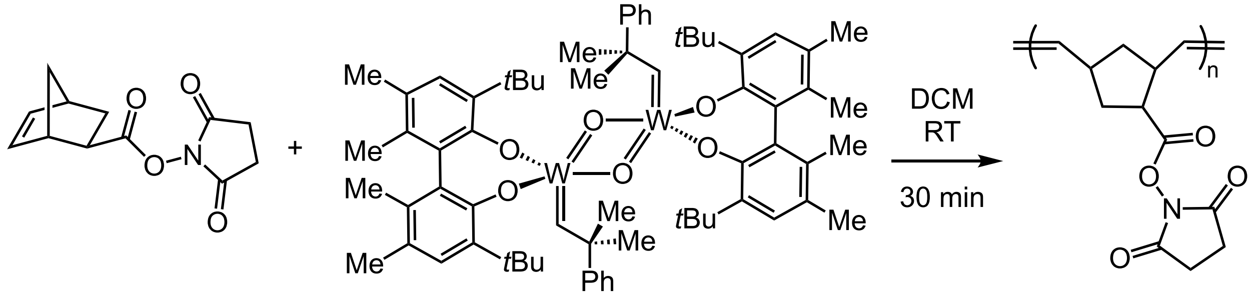


NHS monomer [W] catalyst *cis*-PNB

***Cis*-PNB (100-mer):** Polymer was synthesized according to the modified procedure.[3,5] In a nitrogen-filled glovebox, 450 mg NHS-monomer (1.9 mmol, 100 equiv.) was added to a flame-dried vial equipped with a stir bar. In a separate flame-dried vial, [W] catalyst was weighed out in the glovebox (26 mg, 19 μmol, 1 equiv.). Dry DCM was added in ½ volume to each vial (3 mL per, [0.3 M overall]). The catalyst solution was then added quickly via syringe to the monomer solution whilst stirring. Upon reaction, the solution appeared orange-red. The reaction was allowed to proceed for 30 minutes before removing from the glovebox. The reactant solution was precipitated dropwise into MeOH (100 mL) and decanted to isolate polymer. The product was obtained as a white solid from precipitation. Additional reprecipitations were conducted as necessary based on $^1$H NMR after isolation. $^1$H NMR (400 MHz, methylene chloride-$d_2$) agrees with prior reports. **GPC** ($CHCl_3$, RI by PS standards): $M_n$ = 22 kDa, $Đ$ = 1.64.

***Cis*-PNB (250-mer)**: The same procedure was performed as described above for *cis*-PNB (100-mer). The stoichiometry was adjusted to NHS-monomer (142 mg, 0.60 mmol, 200 equiv) and [W] catalyst (4.13 mg, 3 μmol, 1 equiv.). $^1$H NMR (400 MHz, methylene chloride-$d_2$) agrees with prior reports. **GPC** ($CHCl_3$, RI by PS standards): $M_n$ = 58 kDa, $Đ$ = 1.76.

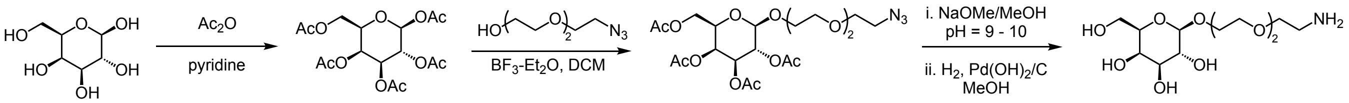


**2-[2-(2-aminoethoxy)ethoxy]ethyl-β-D-galactopyranoside (PEG3Gal):** Synthesis and $^1$H NMR were in agreement with prior reports.[3,6–8] $^1$H NMR: (500 MHz, $D_2O$) δ 4.44 – 4.41 (m, 1H), 4.12 – 4.04 (m, 1H), 3.92 (d, $J$ = 3.4 Hz, 1H), 3.83 (dq, $J$ = 11.8, 3.4 Hz, 1H), 3.80 – 3.63 (m, 11H), 3.59 (t, $J$ = 5.4 Hz, 1H), 3.53 (dd, $J$ = 9.9, 7.9 Hz, 1H), 2.82 (dt, $J$ = 11.7, 5.4 Hz, 2H).

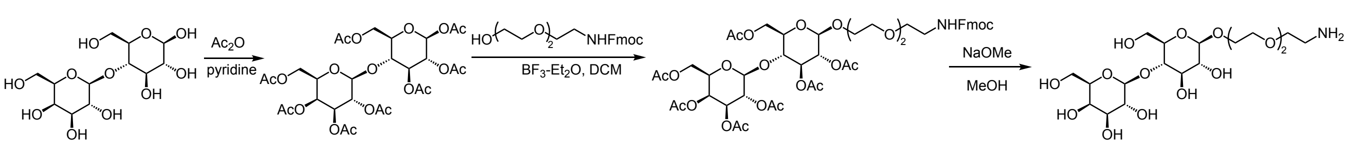


**1-[2-{2-(2-aminoethoxy)ethoxy}ethyl]β-lactoside (PEG3Lac):** Synthesis and $^1$H NMR were in agreement with current literature.[9] $^1$H NMR (400 MHz, $D_2O$): δ 4.45 (d, J = 8.1 Hz, 1H), 4.37 (d, J = 7.8 Hz, 1H), 3.99 (h, J = 4.5 Hz, 1H), 3.90 (d, J = 12.1 Hz, 1H), 3.85 (d, J = 3.4 Hz, 1H), 3.81 – 3.43 (m, 22 H), 3.35 – 3.22 (m, 2H), 3.14 (t, J = 5.0 Hz, 1H), 3.08 (t, J = 5.7 Hz, 2H), 1.70 (p, J = 5.8 Hz, 2H), 1.59 (q, J = 5.7 Hz, 1H).

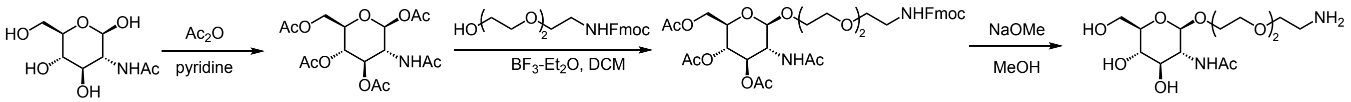

**1-[2-{2-aminoethoxy)ethoxy}ethyl] 2-acetamido-2-deoxy-β-D-glucopyranoside (PEG3GlcNAc):** Synthesis and $^{1}$H NMR is in agreement with current literature.[9] $^{1}$H NMR (600 MHz, $D_2O$) δ 4.56 (d, *J* = 8.5 Hz, 1H), 4.04 (ddd, *J* = 11.6, 6.0, 3.1 Hz, 1H), 3.94 (dd, *J* = 12.2, 1.7 Hz, 1H), 3.82 – 3.67 (m, 11H), 3.59 – 3.53 (m, 1H), 3.49 – 3.44 (m, 2H), 3.22 (dd, *J* = 5.6, 4.6 Hz, 2H), 3.19 – 3.15 (m, 1H), 2.06 (s, 3H), 1.79 (p, J = 5.8 Hz, 1H), 1.68 (tq, J = 9.1, 4.5 Hz, 1H).

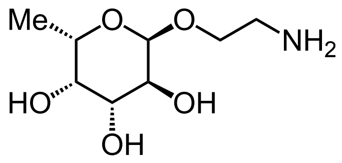

**2-Aminoethyl-α-L-fucopyranoside (PEG1Fuc):** procured by Synthose (CAS: 153252-87-0, product number: AF924). $^{1}$H NMR upon arrival confirmed the structure.

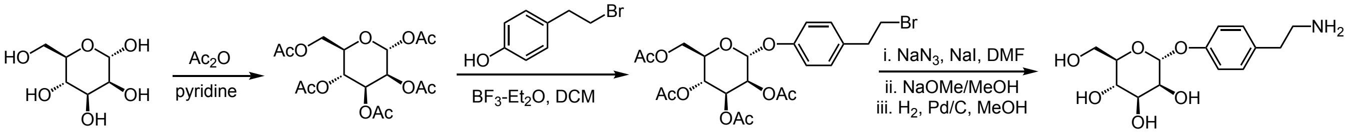

**α-1-O-(4-hydroxyphenylethylamino)-D-mannoside (ArMan):** For the synthesis of aryl mannoside glycoligand, literature procedure was followed.[10] $^{1}$H NMR (400 MHz, $D_2O$) δ 7.31 (d, 2H), 7.17 (d, 2H), 5.61 (d, 1H), 4.18 (dd, 1H), 4.05 (dd, 1H), 3.83-3.71 (m, 4H), 3.26 (t, 2H), 3.97 (t, 2H). 13C NMR (101 MHz, $D_2O$) δ 130.13, 117.32, 98.40, 91.31, 73.33, 70.39, 69.89, 66.57, 60.65, 47.40, 41.88, 36.56.

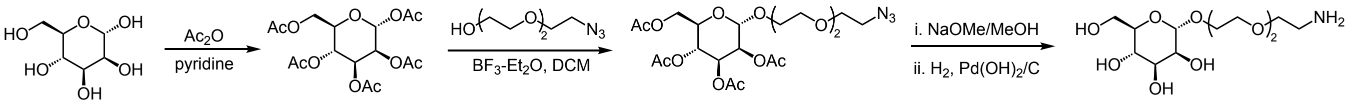

**2-[2-(2-aminoethoxy)ethoxy]ethyl-α-D-mannopyranoside (PEG3Man):** was prepared according to literature procedures.[7] $^{1}$H NMR (400 MHz, $D_2O$) δ ppm 4.90 (d, J = 1.7 Hz, 1H), 3.97 (dd, J = 3.5, 1.8 Hz, 1H), 3.92-3.86 (m, 2H), 3.85-3.80 (m, 1H), 3.78-3.62 (m, 14 H), 2.93 (t, J = 5.3 Hz, 2H). $^{13}$C NMR (101 MHz, $D_2O$) δ 99.91, 72.73, 70.46, 70.38, 69.92, 69.63, 69.48, 69.36, 66.71, 66.30, 60.89, 39.59.

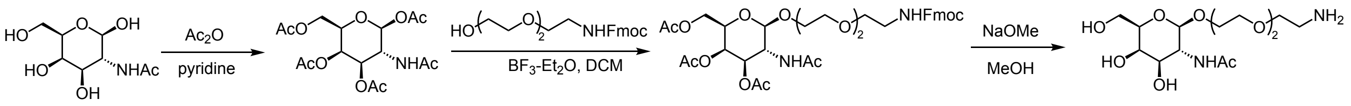

**2-(2-(2-(2-aminoethoxy)ethoxy)-ethoxy)-4,5-dihydroxy-6-(hydroxymethyl)tetrahydro-2H-pyran-3-yl)acetamide (PEG3GalNAc):** Synthesis and $^{1}$H NMR agree with prior report.[11] $^{1}$H

NMR (400 MHz, $D_2O$) δ 4.35 (dd, *J* = 7.9, 1.4 Hz, 1H), 4.06 – 3.94 (m, 1H), 3.84 (d, *J* = 3.4 Hz, 1H), 3.79 – 3.52 (m, 13H), 3.51 – 3.37 (m, 1H), 3.08 (t, *J* = 5.5 Hz, 4H), 1.83 (d, J = 1.5 Hz, 1H), 1.69 (s, 3H), 1.59 (q, *J* = 5.7 Hz, 2H).

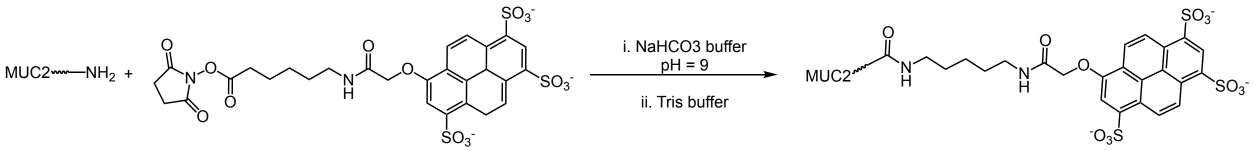


**AF405-labeled MUC2:** The protocol was modified according to a prior report.[12] To fluorescently label the mucin, 0.9 mg was solubilized in 500 µL of [0.1 M] sodium bicarbonate buffer (pH 9) overnight at 4 °C. Alexa Fluor 405 succinimidyl ester (ThermoFisher Scientific) dissolved in DMSO at 10 mg/mL was added to the mucin solution (10 µL), and the mixture was incubated at 4 °C overnight with inverting. Tris(hydroxymethyl)aminomethane [50 mM] was added (10 µL) to quench the reaction and the mixture was left at 4 °C for 3 hours with inverting. The volume was diluted to 20 mL with PBST (PBS with Tween 20) in a Corning Spin-X 100 kDa molecular weight cutoff ultrafiltration concentrator (Fisher) and centrifuged at 3000 G for 25 min at 4 °C; the flow-through was discarded. These steps (dilute to 20 mL with PBST, centrifuge, discard flow-through) were repeated twice more; the volume remaining after each centrifugation was approximately 5 mL. The labeled mucin was then diluted to the appropriate concentration, as measured by a ThermoFisher Nanodrop spectrometer.

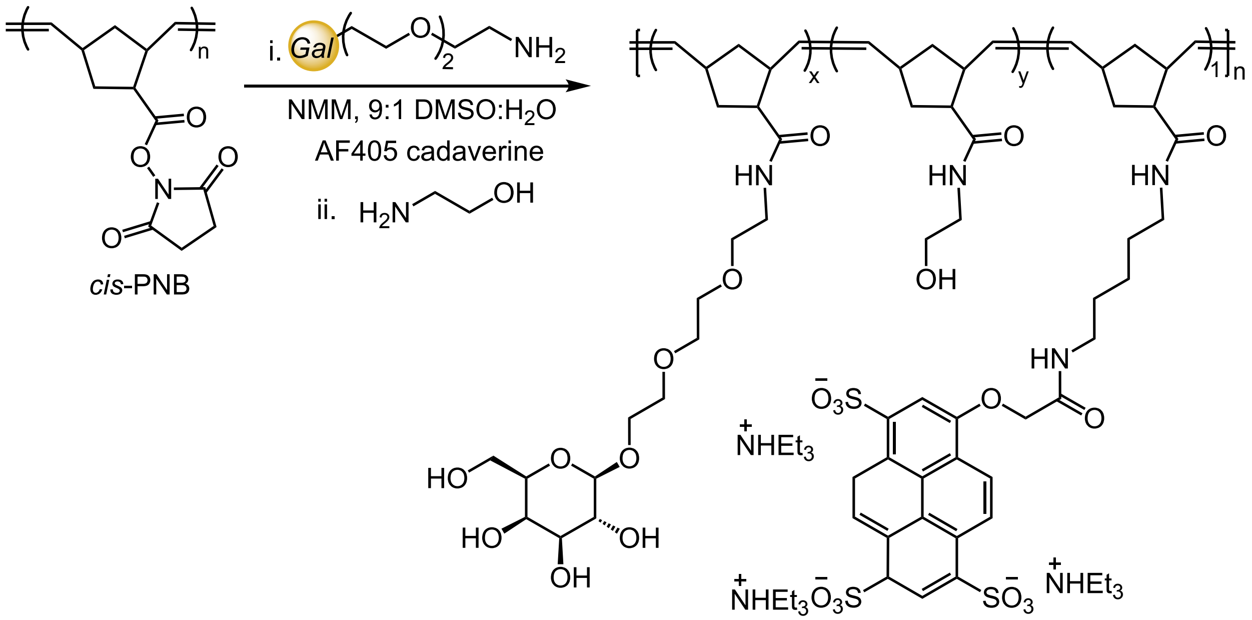


**Gal *cis*-PNB:** To a 1-dram vial equipped with a stir bar, *cis*-PNB 100-mer (10 mg, 0.043 mmol, 1 equiv.) was added. The polymer was dissolved in DMSO (0.85 mL) with stirring. In a separate vial, galactoside (12 mg, 0.032 mmol, 0.75 equiv.) was weighed out and dissolved in DI $H_2O$ (0.15 mL). The galactoside solution was added to the polymer solution to yield a [0.05 M] overall concentration wrt polymer. *N*-methyl morpholine was added (0.12 mL, 1.9 mmol, 25 equiv.) to adjust the pH to 9. Alexa Fluor 405 cadaverine was added via a 1 mg/100 µL solution in DMSO to account for 1 fluorophore per polymer chain (0.28 mg, 28 µL, 0.01 equiv.). The reaction was covered in aluminum foil and stirred overnight. Ethanolamine (0.1 mL) was then added to quench any remaining NHS esters, with stirring for an additional 3 hours. The polymer was purified via dialysis exchange in DI $H_2O$ through 10,000 MWCO dialysis tubing, with 4-6 exchanges over 48 hours. The polymer was isolated as a white fluffy material post-lyophilization. $^1$H NMR determined the extent of galactose incorporation into the polymer, where the vinyl protons at 5.2 ppm were set to integrate to 2 H and the integral of the anomeric peak measured glycan incorporation. $^1$H NMR was in agreement with prior report.[3] $^1$H NMR for 40% Gal *cis*-PNB (400 MHz, $d_2$-DMSO) δ 7.64 (brm, 0.85 H), 5.19 (brm, 2H), 4.84 (s, 0.45 H), 4.64 (m, 5 H), 4.36 (s, 0.35 H), 4.10 (d, $J$ = 6.5 Hz, 0.54 H), 3.85 (d, $J$ = 7.5 Hz, 0.43 H), 3.02–3.60 (brm, 12 H), 2.95 (s, 2 H), 2.33 (m, 1 H), 1.93 (s, 3 H), 1.45 (s, 1 H), 1.23 (s, 0.16 H), 1.02 (s, 1 H).

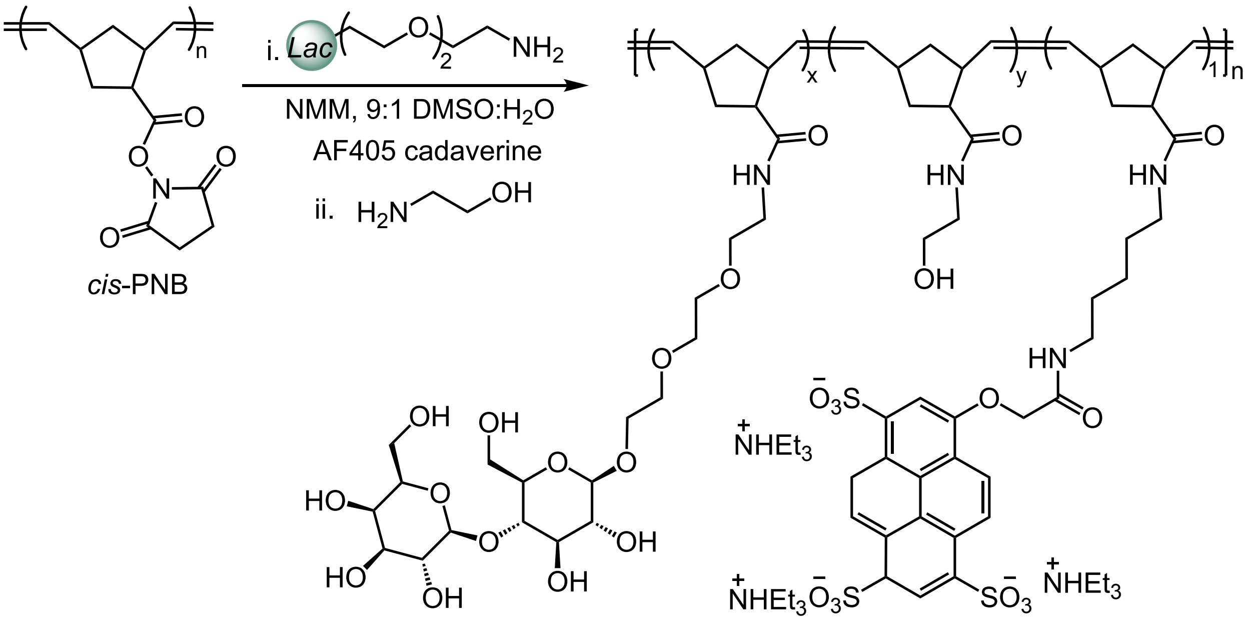


**Lac *cis*-PNB:** To a 1-dram vial equipped with a stir bar, cis-PNB 100-mer (10 mg, 0.043 mmol, 1 equiv.) was added. The polymer was dissolved in DMSO (0.85 mL) with stirring. In a separate vial, lactoside (7 mg, 0.013 mmol, 0.3 equiv.) was weighed out and dissolved in DI $H_2O$ (0.15 mL). The lactoside solution was added to the polymer solution to yield a [0.05 M] overall concentration wrt polymer. *N*-methyl morpholine was added (0.12 mL, 1.08 mmol, 25 equiv.) to adjust the pH to 9. Alexa Fluor 405 cadaverine was added via a 1 mg/100 μL solution in DMSO to account for 1 fluorophore per polymer chain (0.28 mg, 28 μL, 0.01 equiv.). The reaction was covered in aluminum foil and stirred overnight. Ethanolamine (0.1 mL) was then added to quench any remaining NHS esters, with stirring for an additional 3 hours. The polymer was purified via dialysis exchange in DI $H_2O$ through 10,000 MWCO dialysis tubing, with 4-6 exchanges over 48 hours. The polymer was isolated as a white fluffy material post-lyophilization. $^1$H NMR determined the extent of lactose incorporation into the polymer, where the vinyl protons at 5.2 ppm were set to integrate to 2 H and the integral of the anomeric peak measured glycan incorporation. $^1$H NMR for 10% Lac *cis*-PNB (400 MHz, DMSO) δ 7.84 – 7.48 (m, 0.68 H), 5.46 – 4.94 (m, 2 H), 4.86 – 4.47 (m, 0.62 H), 4.22 (ddt, $J$ = 9.2, 7.5, 1.6 Hz, 0.23

H), 3.92 – 3.71 (m, 0.19 H), 3.66 – 2.79 (m, 12 H), 2.44 – 2.27 (m, 0.85 H), 2.07 – 1.76 (m, 2 H), 1.60 – 1.33 (m, 1 H), 1.25 (d, *J* = 3.4 Hz, 0.47 H), 1.16 – 0.93 (m, 0.66 H).

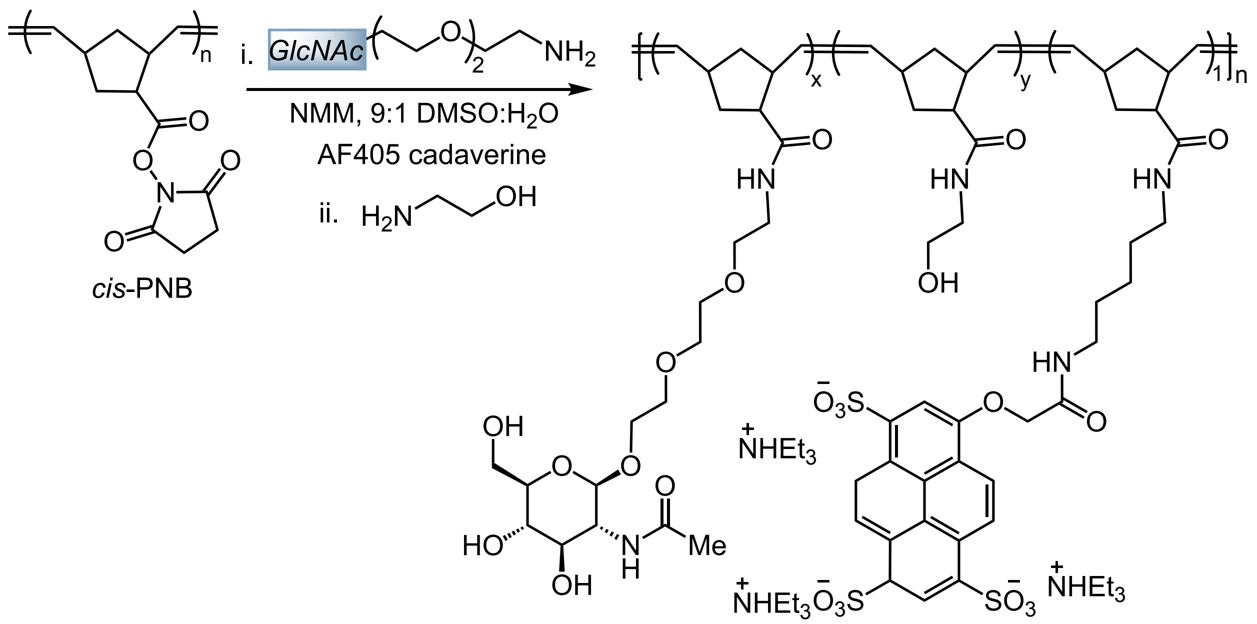


**GlcNAc *cis*-PNB:** To a 1-dram vial equipped with a stir bar, *cis*-PNB 100-mer (10 mg, 0.043 mmol, 1 equiv.) was added. The polymer was dissolved in DMSO (0.85 mL) with stirring. In a separate vial, *N*-acetyl glucosaminoside (9 mg, 0.022 mmol, 0.5 equiv.) was weighed out and dissolved in DI $H_2O$ (0.15 mL). The glycan solution was added to the polymer solution to yield a [0.05 M] overall concentration wrt polymer. *N*-methyl morpholine was added (0.12 mL, 1.08 mmol, 25 equiv.) to adjust the pH to 9. Alexa Fluor 405 cadaverine was added via a 1 mg/100 μL solution in DMSO to account for 1 fluorophore per polymer chain (0.28 mg, 28 μL, 0.01 equiv.). The reaction was covered in aluminum foil and stirred overnight. Ethanolamine (0.1 mL) was then added to quench any remaining NHS esters, with stirring for an additional 3 hours. The polymer was purified via dialysis exchange in DI $H_2O$ through 10,000 MWCO dialysis tubing, with 4-6 exchanges over 48 hours. The polymer was isolated as a white fluffy material post-lyophilization. $^1$H NMR determined the extent of GlcNAc incorporation into the polymer, where the vinyl protons at 5.2 ppm were set to integrate to 2 H and the integral of the anomeric peak measured glycan incorporation. $^1$H NMR for 50% GlcNAc *cis*-PNB (400 MHz, DMSO): δ 7.69 (d, *J* = 9.0 Hz, 0.67 H), 5.20 (bs, 2 H), 5.02 – 4.87 (bm, 0.81 H), 4.64 (bs, 0.45 H), 4.54 (bs, 0.58 H), 4.33 (d, *J* = 8.3 Hz, 0.58 H), 3.80 (d, *J* = 5.4 Hz, 0.55 H), 3.73 – 3.63 (m, 0.64 H), 3.58 – 2.78 (m, 12 H), 2.43 – 2.23 (m, 1 H), 1.92 (q, *J* = 17.6 Hz, 2 H), 1.61 – 1.30 (m, 2 H), 1.24 (s, 0.35 H), 1.03 (bs, 1H).

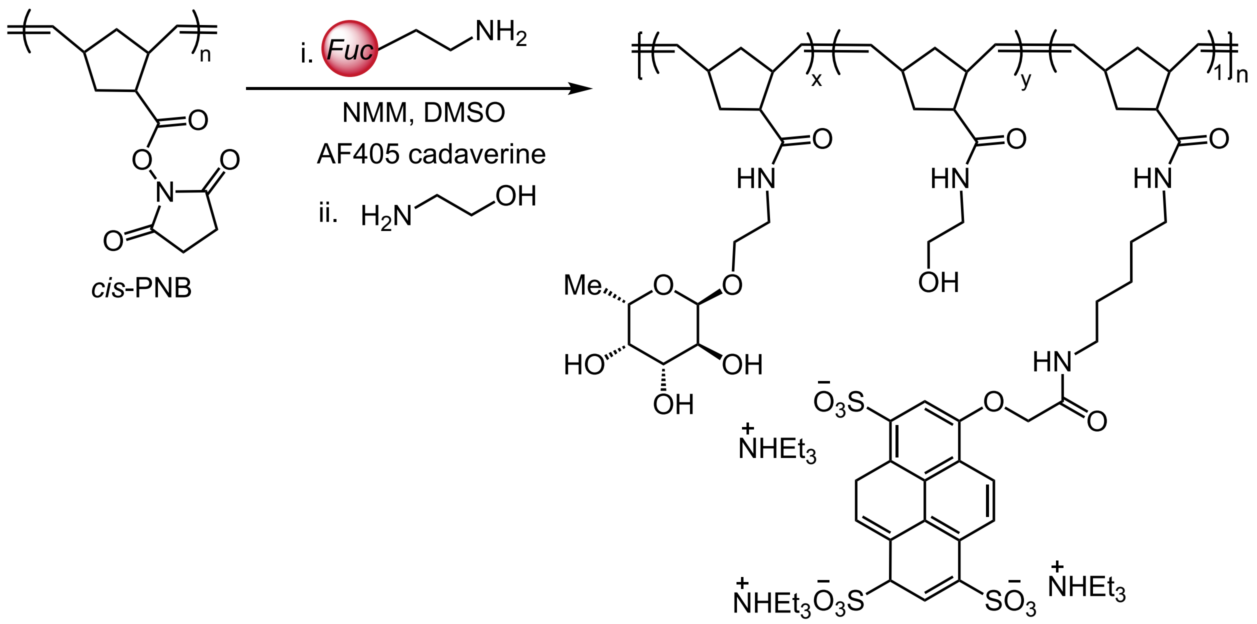


**Fuc *cis*-PNB:** To a 1-dram vial equipped with a stir bar, *cis*-PNB 100-mer (10 mg, 0.043 mmol, 1 equiv.) was added. The polymer was dissolved in DMSO (0.85 mL) with stirring. Upon arrival from Synthose™, fucoside was dissolved in DMSO to generate a 25 mg/mL solution. The glycan solution (4.5 mg, 0.18 mL) was added to the polymer solution. *N*-methyl morpholine was added (0.12 mL, 1.08 mmol, 25 equiv.) to adjust the pH to 9. Alexa Fluor 405 cadaverine was added via a 1 mg/100 mL solution in DMSO to account for 1 fluorophore per polymer chain (0.28 mg, 28 µL, 0.01 equiv.). The reaction was covered in aluminum foil and stirred overnight. Ethanolamine (0.1 mL) was then added to quench any remaining NHS esters, with stirring for an additional 3 hours. The polymer was purified via dialysis exchange in DI $H_2O$ through 10,000 MWCO dialysis tubing, with 4-6 exchanges over 48 hours. The polymer was isolated as a white fluffy material post-lyophilization. $^1$H NMR determined the extent of Fuc incorporation into the polymer, where the vinyl protons at 5.2 ppm were set to integrate to 2 H and the integral of the anomeric peak measured glycan incorporation. $^1$H NMR for 30% Fuc *cis*-PNB (400 MHz, DMSO): δ 7.66 (s, 0.80 H), 5.32 – 5.04 (m, 2 H), 4.62 (d, *J* = 15.0 Hz, 1 H), 4.49 (s, 0.25 H), 4.40 (s, 0.32 H), 4.31 (s, 0.23 H), 3.78 (d, *J* = 8.5 Hz, 0.33 H), 3.60 – 3.45 (m, 1 H), 3.23 – 2.83 (m, 0.37 H), 1.93 (d, *J* = 15.4 Hz, 2 H), 1.57 – 1.33 (m, 1 H), 1.07 (q, *J* = 5.8 Hz, 1 H).

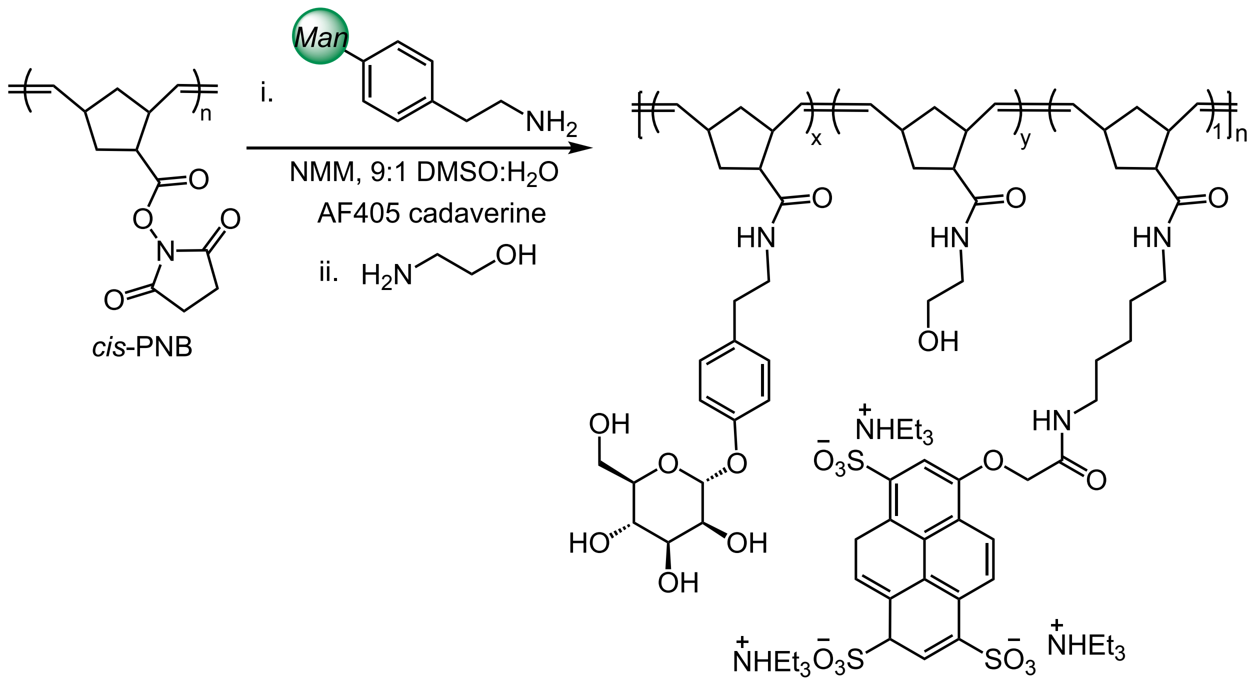


**ArMan *cis*-PNB:** To a 1-dram vial equipped with a stir bar, *cis*-PNB 100-mer (10 mg, 0.043 mmol, 1 equiv.) was added. The polymer was dissolved in DMSO (0.85 mL) with stirring. In a separate vial, the mannoside was weighed out (5 mg, 0.017 mmol, 0.4 equiv.) and dissolved in DI $H_2O$ (0.1 mL). The glycan solution was added to the polymer solution. *N*-methyl morpholine was added (0.12 mL, 1.08 mmol, 25 equiv.) to adjust the pH to 9. Alexa Fluor 405 cadaverine was added via a 1 mg/100 mL solution in DMSO to account for 1 fluorophore per polymer chain (0.28 mg, 28 µL, 0.01 equiv.). The reaction was covered in aluminum foil and stirred overnight. Ethanolamine (0.1 mL) was then added to quench any remaining NHS esters, with stirring for an additional 3 hours. The polymer was purified via dialysis exchange in DI $H_2O$ through 10,000 MWCO dialysis tubing, with 4-6 exchanges over 48 hours. The polymer was isolated as a white fluffy material post-lyophilization. $^1H$ NMR determined the extent of ArMan incorporation into the polymer, where the vinyl protons at 5.2 ppm were set to integrate to 2 H and the integral of the anomeric peak measured glycan incorporation. $^1H$ NMR for 40% ArMan *cis*-PNB (400 MHz, DMSO): δ 7.84 – 7.53 (m, 1.17 H), 7.18 – 6.85 (m, 1.73 H), 5.20 (bs, 2 H), 4.97 (s, 0.15 H), 4.65 (bs, 0.44 H), 4.45 (bs, 0.22 H), 3.82 (bs, 0.49 H), 3.67 (s, 0.44 H), 2.97 (bs, 2 H), 1.94 (bs, 3 H), 1.46 (bs, 1 H), 1.03 (s, 1H).

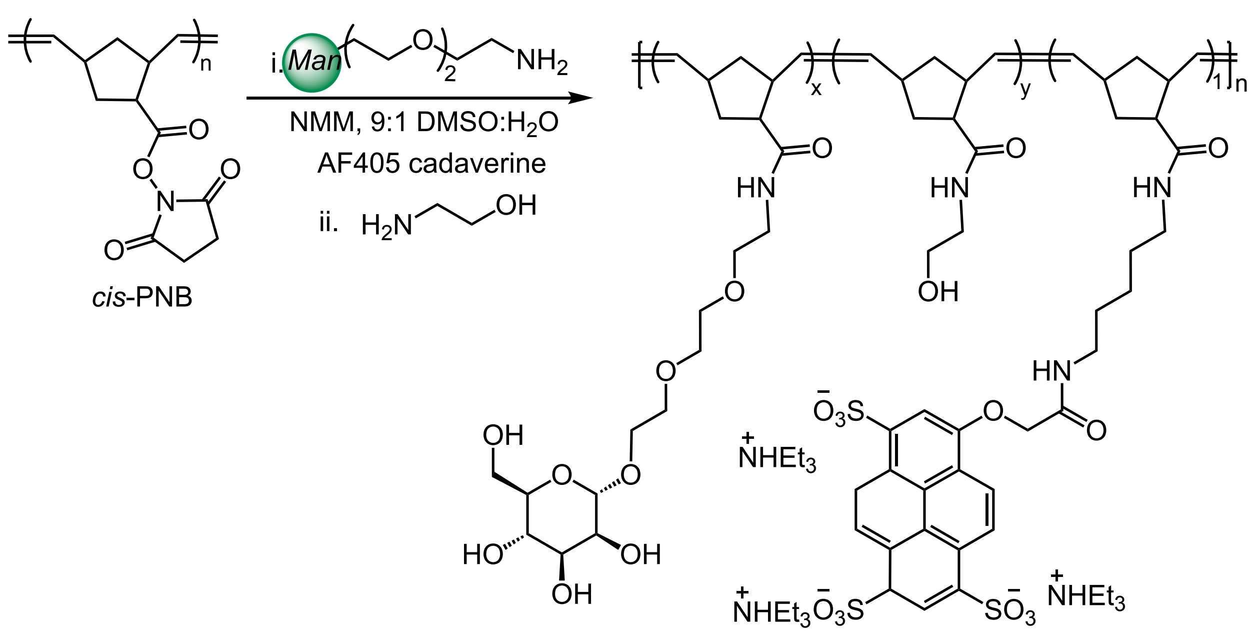


**Man *cis*-PNB:** To a 1-dram vial equipped with a stir bar, *cis*-PNB 100-mer (10 mg, 0.043 mmol, 1 equiv.) was added. The polymer was dissolved in DMSO (0.85 mL) with stirring. In a separate vial, the mannoside was weighed out (8 mg, 0.022 mmol, 0.5 equiv.) and dissolved in DI $H_2O$ (0.1 mL). The glycan solution was added to the polymer solution. *N*-methyl morpholine was added (0.12 mL, 1.08 mmol, 25 equiv.) to adjust the pH to 9. Alexa Fluor 405 cadaverine was added via a 1 mg/100 mL solution in DMSO to account for 1 fluorophore per polymer chain (0.28 mg, 28 μL, 0.01 equiv.). The reaction was covered in aluminum foil and stirred overnight. Ethanolamine (0.1 mL) was then added to quench any remaining NHS esters, with stirring for an additional 3 hours. The polymer was purified via dialysis exchange in DI $H_2O$ through 10,000 MWCO dialysis tubing, with 4-6 exchanges over 48 hours. The polymer was isolated as a white fluffy material post-lyophilization. $^1$H NMR determined the extent of Man incorporation into the polymer, where the vinyl protons at 5.2 ppm were set to integrate to 2 H and the integral of the anomeric peak measured glycan incorporation. $^1$H NMR for 30% Man *cis*-PNB (600 MHz, DMSO): δ 7.85 – 7.55 (m, 1.13 H), 5.30 – 5.03 (m, 2 H), 4.76 – 4.69 (m, 0.66 H), 4.64 (bs, 0.49 H), 4.56 (bs, 0.24 H), 4.45 (bs, 0.33 H), 3.71 – 2.70 (m, 12 H), 2.37 (bs, 1 H), 2.05 – 1.78 (m, 2 H), 1.55 – 1.36 (m, 1 H), 1.04 (bs, 1 H).

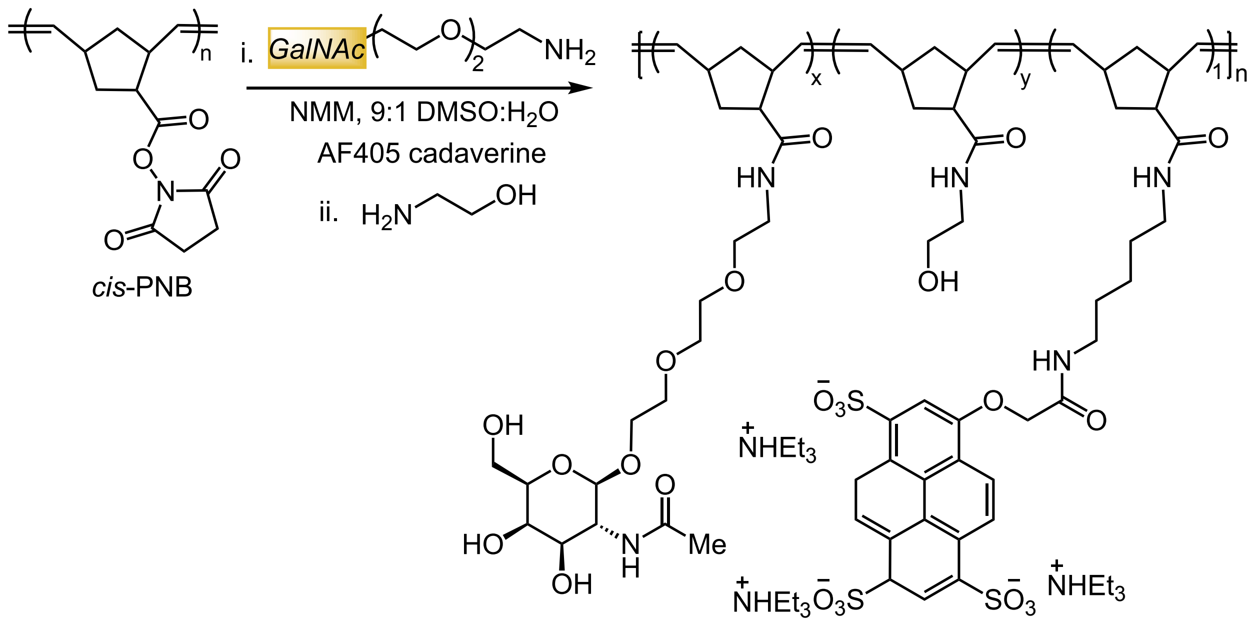


**GalNAc *cis*-PNB:** To a 1-dram vial equipped with a stir bar, *cis*-PNB 100-mer (10 mg, 0.043 mmol, 1 equiv.) was added. The polymer was dissolved in DMSO (0.85 mL) with stirring. In a separate vial, the *N*-acetyl galactosaminoside was weighed out (9 mg, 0.022 mmol, 0.5 equiv.) and dissolved in DI $H_2O$ (0.1 mL). The glycan solution was added to the polymer solution. *N*-methyl morpholine was added (0.12 mL, 1.08 mmol, 25 equiv.) to adjust the pH to 9. Alexa Fluor 405 cadaverine was added via a 1 mg/100 mL solution in DMSO to account for 1 fluorophore per polymer chain (0.28 mg, 28 µL, 0.01 equiv.). The reaction was covered in aluminum foil and stirred overnight. Ethanolamine (0.1 mL) was then added to quench any remaining NHS esters, with stirring for an additional 3 hours. The polymer was purified via dialysis exchange in DI $H_2O$ through 10,000 MWCO dialysis tubing, with 4-6 exchanges over 48 hours. The polymer was isolated as a white fluffy material post-lyophilization. $^1$H NMR determined the extent of GalNAc incorporation into the polymer, where the vinyl protons at 5.2 ppm were set to integrate to 2 H and the integral of the anomeric peak measured glycan incorporation. $^1$H NMR for 30% GalNAc *cis*-PNB (400 MHz, DMSO): δ 7.63 (s, 0.44 H), 5.20 (bs, 2 H), 4.85 (bs, 0.26 H), 4.72 – 4.57 (m, 0.73 H), 4.37 (bs, 0.24 H), 4.10 (d, *J* = 6.2 Hz, 0.28 H), 3.84 (bs, 0.32 H), 3.67 – 2.72 (m, 12 H), 1.91 (bs, 2 H), 1.66 – 1.32 (m, 2 H), 1.01 (s, 1 H).

**4. Flow Cytometry Protocol.** After overnight culture, the OD600 was measured to ensure the cells reached stationary phase. Depending on the experiment, the cells were either used immediately or allowed to recover into mid-log phase after dilution. The cells were pelleted at 3,000 G for 10 min and growth media was aspirated. The cells were resuspended in HEPES with added calcium to OD = 1. Cells were added to a Delta Nunclon 96-well plate to an OD = 0.4 per well. To each well, fluorophore-labeled synthetic or native mucins were added at 20% aqueous DMSO stocks as high as [20 mM] with respect to glycan, varying concentration to test dose-dependence: [2 mM], [500 µM], and [50 µM] final concentrations in HEPES with added calcium or HEPES with EDTA. The polymers and cells were incubated at 4 °C for 1 hour with shaking at 160 rpm. After incubation, the cells were pelleted at 3,000 G for 5 min and the supernatant was aspirated to remove unbound polymers. The cells were resuspended in HEPES with calcium or HEPES with EDTA. SYTO-BC (Invitrogen™) was dissolved to [100 mM] in DMSO and added to the wells as 1 μL. The plate was allowed to shake for 10-30 minutes at 4 °C and 160 rpm for the cells to be sufficiently labeled.

The fluorophore labeled cells and polymers were analyzed using an Attune NxT Flow Cytometer (405 nm, 488 nm, 561 nm, and 640 nm lasers). Flow cytometry analysis was performed in triplicate, and representative scatter plots are shown. The dye only, cells only, and polymer only controls were analyzed first to set gates. Data were analyzed using the FlowJo software package (FlowJo LLC). Percent cells bound was determined by cells that displayed SYTO and AF405 fluorescence signal.

**5. Microscopy Protocol**

For microscopy, 5 μl of each sample were spotted onto a glass-bottomed microwell dish (Nunc 501) and diluted with 95 μL of HEPES with added calcium or EDTA. Images were collected on a MetaXpress confocal HTAI microscope (40x water immersion lens). Brightness and contrast were identically adjusted with the open-source Fiji distribution of ImageJ. Images were then converted to an RGB format to preserve normalization and then assembled into panels.

**6. Clustering Protocol**

Images were collected on a MetaXpress confocal HTAI microscope (40x water immersion lens). Brightness and contrast of the SYTO channel were identically adjusted with the open-source Fiji distribution of ImageJ. The threshold was set to mask the particles and analyze their area, with less than 2% of the population appearing so that the software could analyze the particle edges well. In Excel, the area measurement was transformed into diameter and the distribution of cell diameters was measured by a normal distribution probability function, according to a literature procedure.[13] The diameter was then plotted against the distribution to show a size dispersity after the cells incubated with glycopolymer.

**7. Adhesion Protocol**

The following was modified from a previously reported procedure.[14] Within the MIT.nano cleanroom facility, a black, flat-bottomed 96-well polystyrene plate was oxygenated using soft lithography of oxygen plasma for 1 min. Aminopropyltriethoxysilane (APTES) was added in a 5% (v/v) in ethanol solution to each well (250 μL). The reaction of hydroxyls and APTES installed primary amines on the surface of the plate. Separately, MUC2 (10 mg) was dissolved in PBS (1 mL) and then subjected to EDC (0.775 mg in 0.1 mL PBS, 5 equiv.). MUC2 and EDC were inverted in an epitube at 4 °C for 5 minutes. Then, NHS (1.18 mg in 0.08 mL PBS, 10 equiv.) was added to the tube and the reaction was inverted for another 5 minutes at 4 °C. After 15 minutes or more, the APTES solution was aspirated from the 96-well plate in a chemical hood and each well was washed with 250 μL ethanol 2x. After the final wash, the plate was dried under air stream. The MUC2-NHS was further diluted to 30% in PBS (+2.7 mL). 120 μL of 30% MUC2-NHS was added to each well. Control wells were filled with PBS instead. The plate was sealed with parafilm and rocked overnight at 4 °C.

Bacteria were grown overnight in MRS broth to stationary phase. The growth was measured by OD600 and the bacteria were pelleted out of MRS broth at 3,000 G for 10 minutes at 4 °C. The bacteria were resuspended to OD = 1 in PBS. Carboxyfluorescein diacetate (cFDA) was added to a final concentration of [100 μM]. The bacteria were incubated with cFDA for 4 h at 37 °C with 190 rpm in falcon tubes. Excess fluorophore was washed away through pelleting and resuspending in PBS 3x, with centrifugation set at 3,000 G for 5 min at 4 °C. The fluorescent bacteria were finally resuspended in PBS to OD = 1.

MUC2-NHS was aspirated from the 96-well plate. Each well was filled with 100 μL of PBS buffer, fluorescent bacterial cells, and/or AF405-labelled glycopolymer. The bacteria were incubated on the MUC2-coated plate for 4 h at 37 °C with shaking. The plate was then washed 3x with PBS aspirations. The plate was read by a SpectraMax plate reader for commensal cell counts (cFDA fluorescence 493 → 517 nm) and glycopolymer presence (AF405 fluorescence 401 → 421 nm).

**8. Quantitative Real-Time PCR Protocol**

Glycosidase expression was measured by quantitative real-time PCR (qPCR) following growth in media conditions supplemented with and without native mucin (MUC2). To investigate whether glycosidase expression changed in different growth conditions, the bacteria of interest (*L. reuteri, L. plantarum,* or *L. fermentum*) were grown in minimal media (MOD-MRS), MRS, or MRS supplemented with MUC2 (1 wt%). Total RNA was extracted with Trizol (ThermoFisher Scientific) using a Direct-zol RNA mini-prep kit (Zymo Research). cDNA was generated using GoScript Reverse Transcriptase with Random Primers (Promega). PCR amplification was performed in the presence of SYBR green (BioRad) in a CFX96 real-time PCR detection system (BioRad). Specific primers were designed using the PrimerQuest tool (Integrated DNA Technologies, Inc., **Supplementary Table 2**). Expression of specific genes was normalized to 16S expression (ΔCt), and expression fold change was calculated using the delta Ct method ($2^{-(\Delta Ct\ +MUC2\ -\ \Delta Ct\ -MUC2)}$.

### 9. Polymer NMRs and GPCs

**Figure S24.** *Cis*-PNB NMR (100-mer) in deuterated dichloromethane.

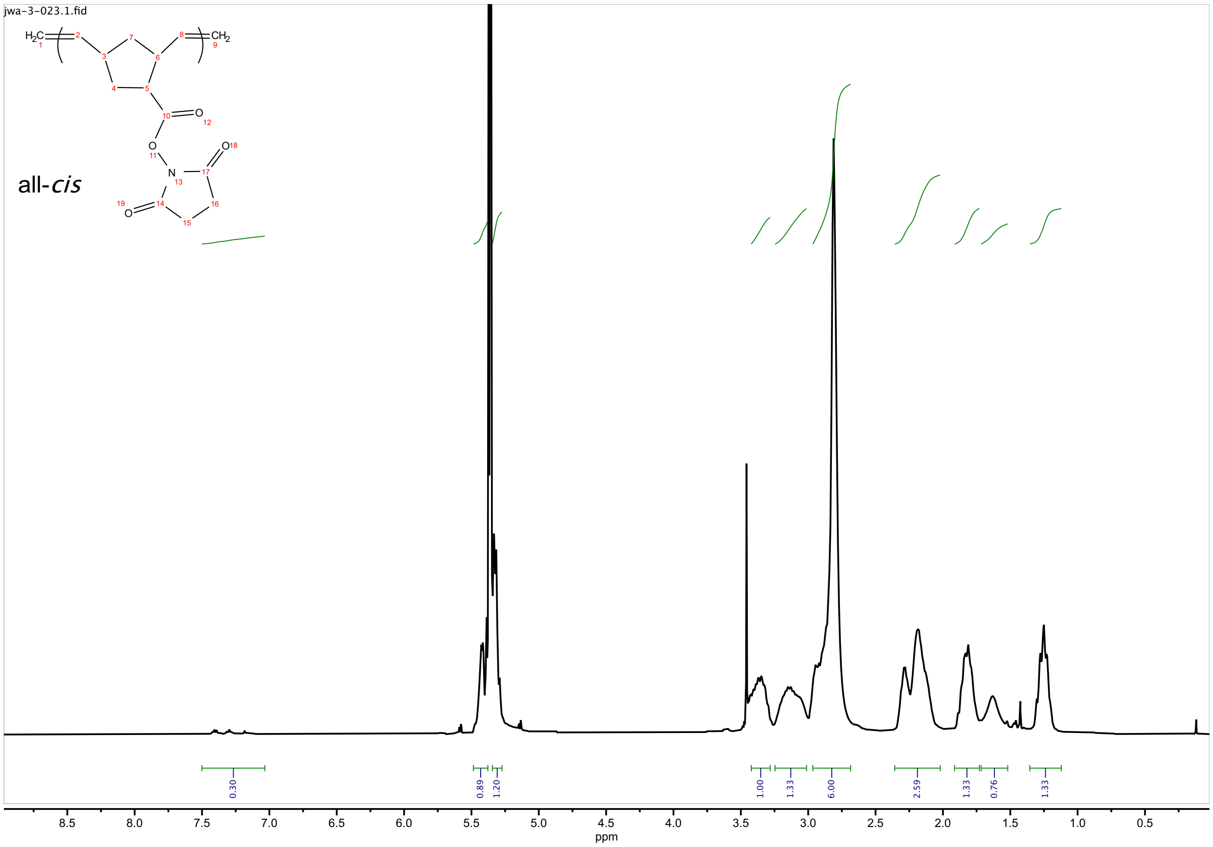

**Figure S25.** *Cis*-PNB GPC (100-mer)

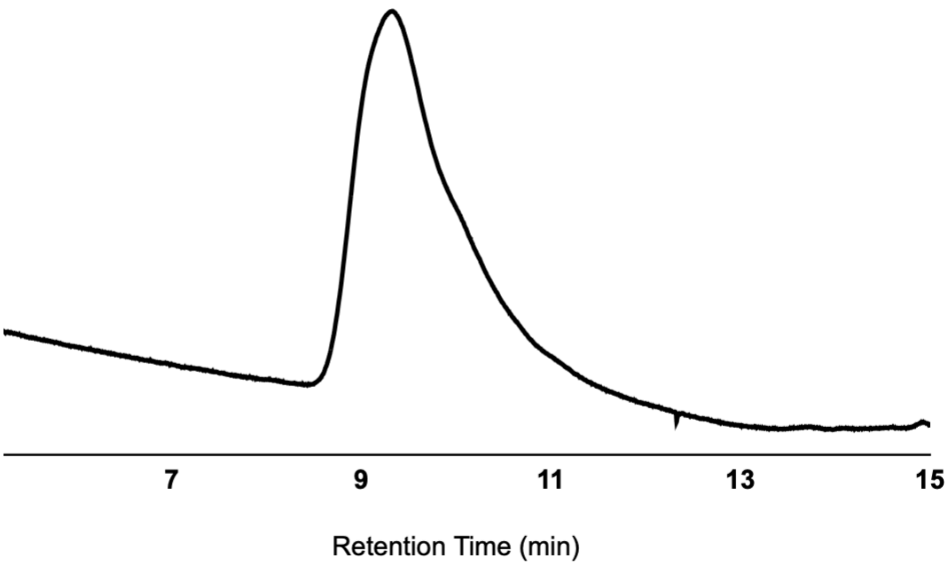


**Figure S26.** *Cis*-PNB GPC (250-mer)

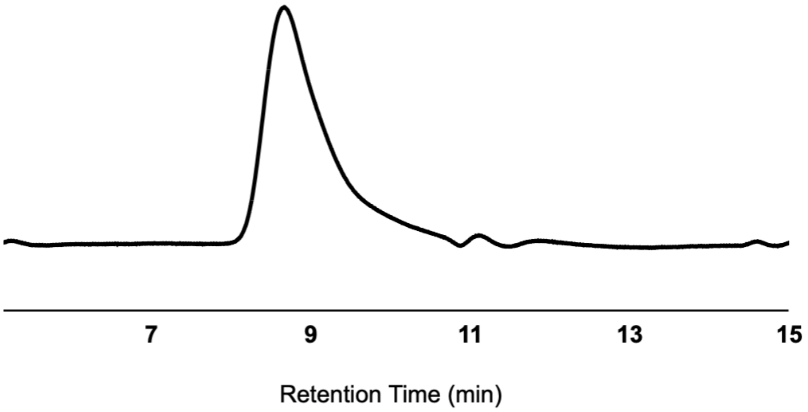

**Figure S27.** Representative $^1$H NMR of galactosylated *cis*-PNB in d-DMSO: 40% Gal cis-PNB NMR (100-mer)

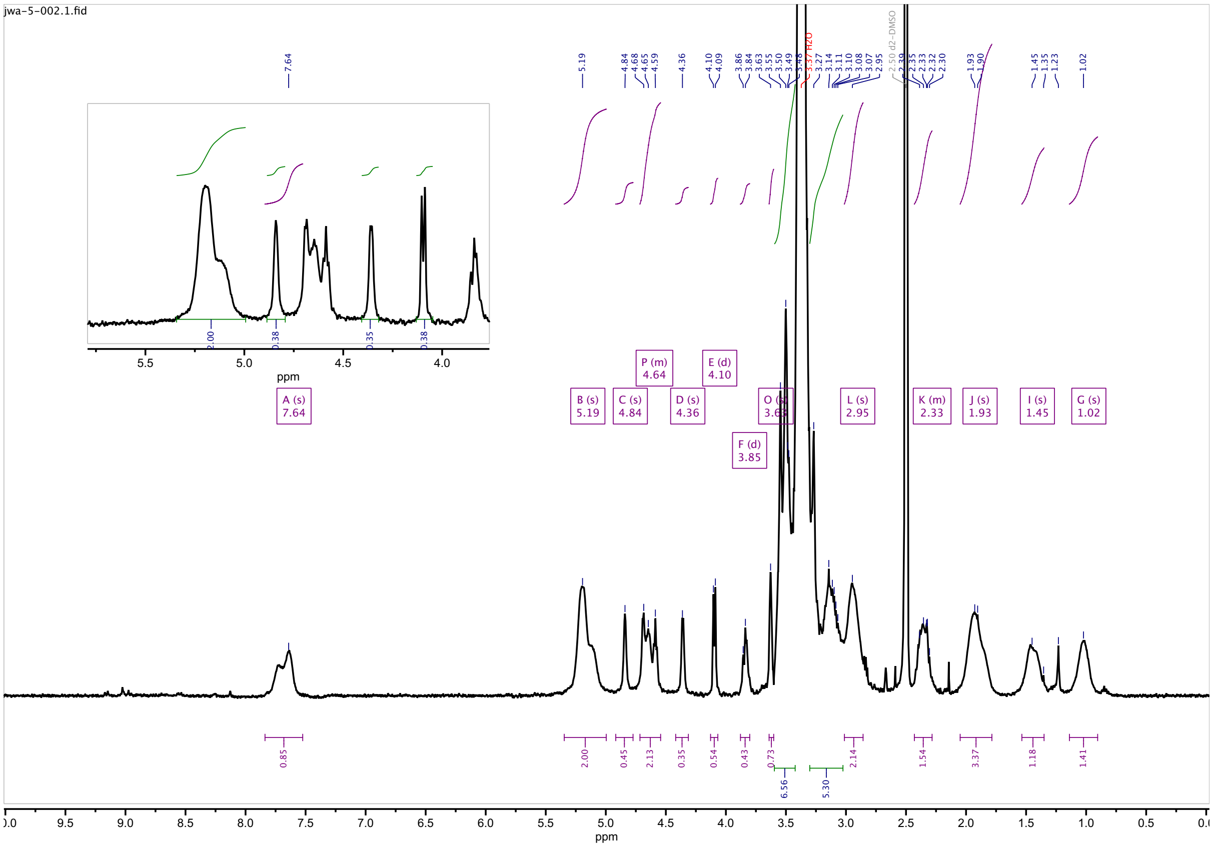

**Figure S28.** Representative $^1$H NMR of lactosylated *cis*-PNB in d-DMSO: 10% Lac cis-PNB NMR (100-mer)

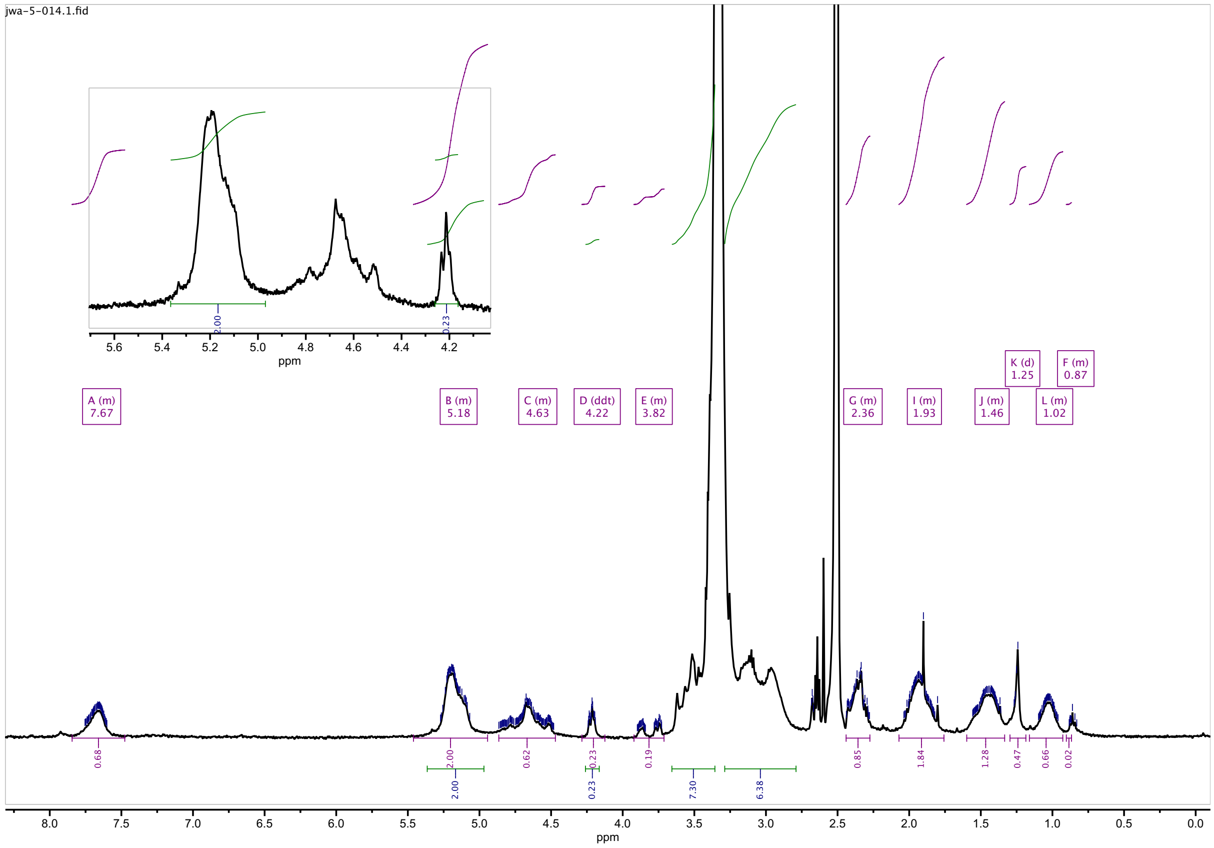

**Figure S29.** Representative $^1$H NMR of *N*-acetyl glucosaminated *cis*-PNB in d-DMSO: 50% GlcNAc *cis*-PNB (100-mer)

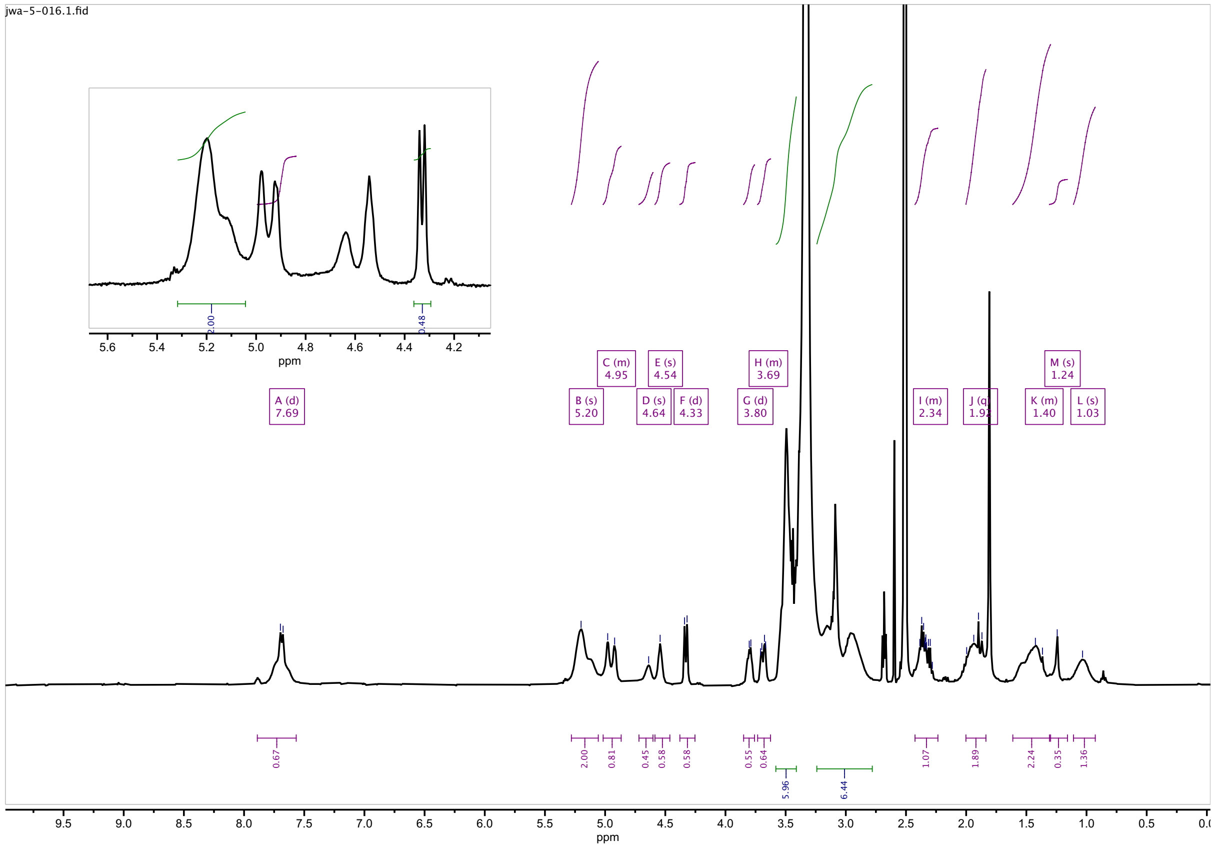

**Figure S30.** Representative $^1$H NMR of *N*-acetyl galactosaminated *cis*-PNB in d-DMSO: 30% GalNAc *cis*-PNB (100-mer)

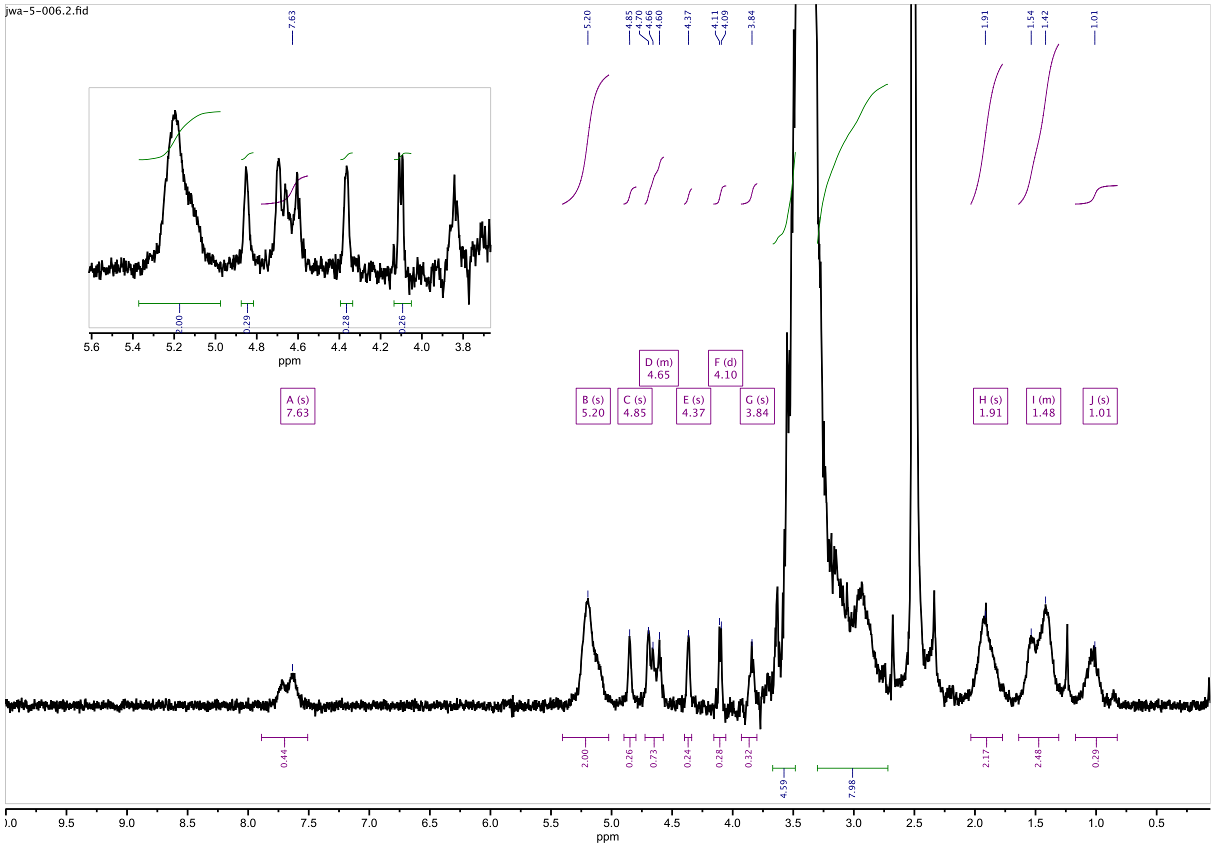

**Figure S31.** Representative $^{1}$H NMR of fucosylated *cis*-PNB in d-DMSO: 30% Fuc *cis*-PNB (100-mer)

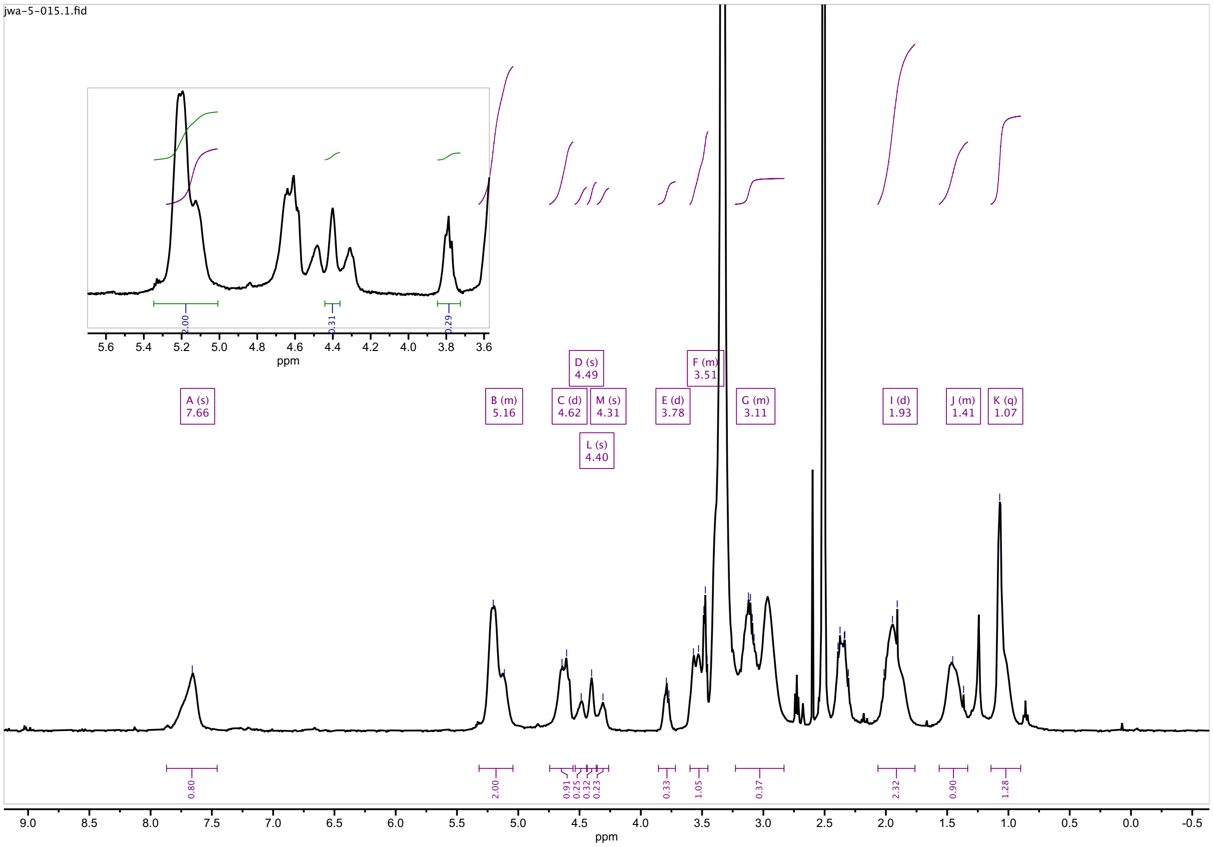

**Figure S32.** Representative $^1H$ NMR of mannosylated *cis*-PNB in d-DMSO: 30% PEG3Man *cis*-PNB (100-mer)

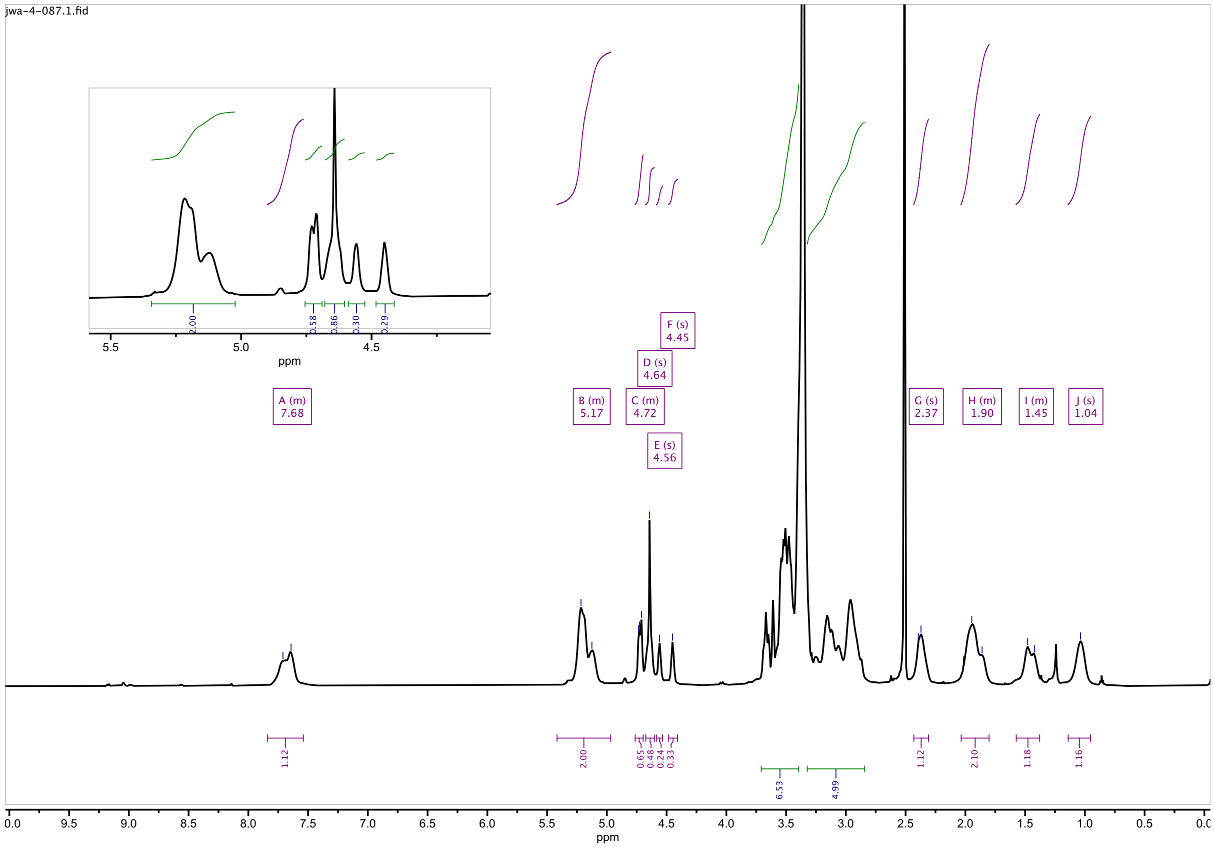

**Figure S33.** Representative $^{1}$H NMR of mannosylated *cis*-PNB in d-DMSO: 40% ArMan *cis*-PNB (100-mer)

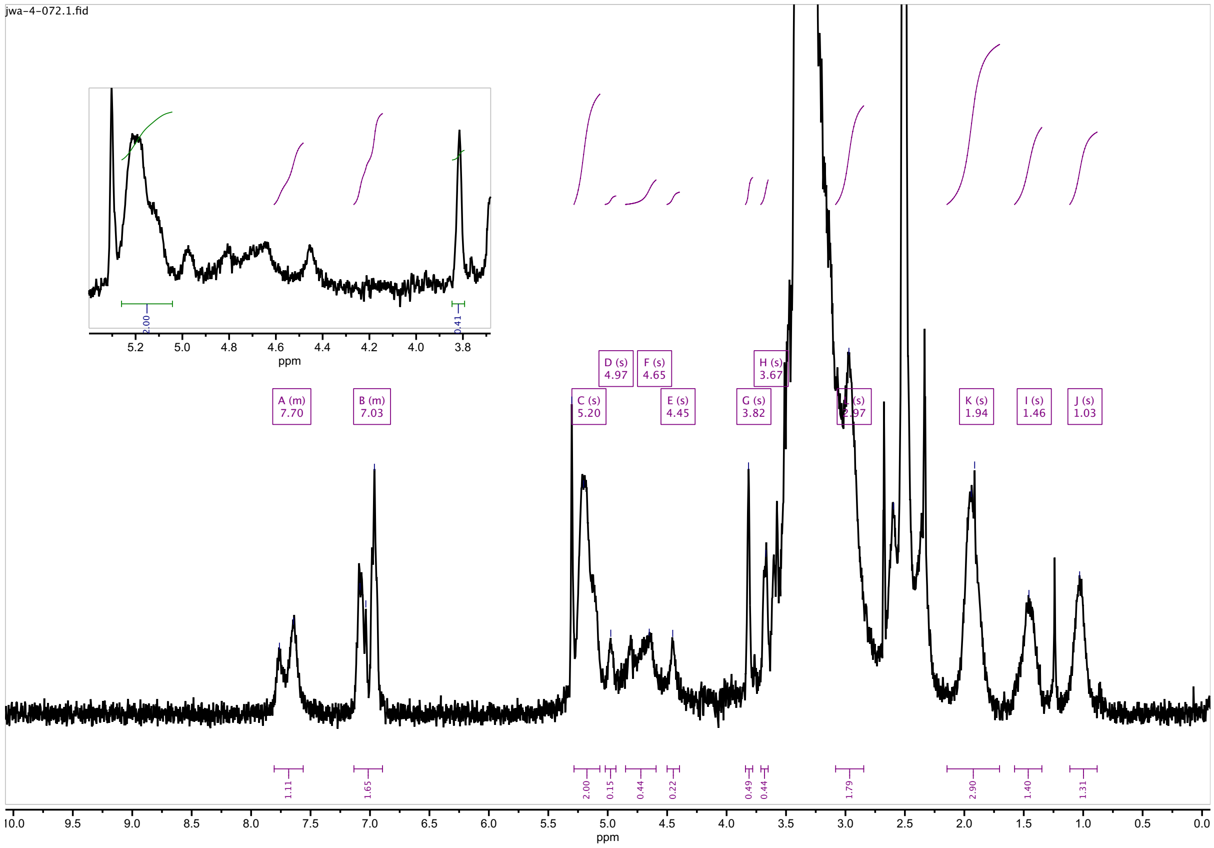
